# Robust somatic copy number estimation using coarse-to-fine segmentation

**DOI:** 10.1101/2021.08.06.455329

**Authors:** Luka Culibrk, Jasleen K Grewal, Erin D Pleasance, Laura Williamson, Karen Mungall, Janessa Laskin, Marco A Marra, Steven JM Jones

## Abstract

Cancers routinely exhibit chromosomal instability that results in copy number variants (CNVs), namely changes in the abundance of genomic material. Unfortunately, the detection of these variants in cancer genomes is difficult. We present Ploidetect, a software package that effectively identifies CNVs within whole-genome sequenced tumors. Ploidetect utilizes a coarse-to-fine segmentation approach which yields highly contiguous segments while allowing for focal CNVs to be detected with high sensitivity. We benchmark Ploidetect against popular CNV tools using synthetic data, cell line data, and real-world metastatic tumor data, and find an improvement in sensitivity and accuracy while maintaining low levels of oversegmentation. We show that the improved CNV sensitivity enables Ploidetect to recover recurrent homozygous deletions and genes associated with chromosomal instability in a multi-cancer cohort of 687 patients. Using highly contiguous CNV calls afforded by Ploidetect, we also demonstrate the use of segment N50 as a novel metric for the measurement of chromosomal instability within tumor biopsies. We propose that increasingly accurate determination of CNVs is critical for their productive study in cancer, and our work demonstrates advances made possible by progress in this regard.

## Introduction

Copy number variation (CNV) is a major class of somatic mutations often observed in the context of cancer. CNV has been shown to impact the majority of cancer genomes (***Zack et al., 2013***) and is associated with poor clinical outcomes (***Hieronymus et al., 2018***). Homozygous deletions are CNVs that result in the complete loss of genomic loci, and recurrent homozygous deletions are known to be enriched in tumor suppressor genes (***Cheng et al., 2017***). Homologous recombination deficiency (HRD), a cause of CNV, has been positively associated with clinical platinum therapy response (***Melinda et al., 2016***), and other phenotypes such as an increased prevalence of tandem duplications have been described with relevance to biological processes (***Menghi et al., 2018***; ***Macintyre et al., 2018***). Chromosomal instability (CIN) driving the accumulation of CNVs is associated with poor patient outcome in multiple cancer types (***Heilig et al., 2010***; ***Orsetti et al., 2014***; ***Choi et al., 2009***). Overall, CNV is a common class of mutations with relevance in numerous facets of tumor biology and clinical management of cancer. Although CNV is a highly pervasive mechanism by which tumors accrue somatic mutation, these variants are considerably more difficult to detect accurately compared to other types of mutations.

While small mutations can be determined to a high degree of accuracy through base changes embedded within aligned sequence reads, CNVs are variations in DNA quantity and are typically determined through measurement of relative DNA abundance by way of sequence read depth (***Xi et al., 2016***; ***Favero et al., 2015***). Due to random noise and sequence bias, raw read depth data is unreliable and therefore must be noise corrected. Segmentation, the process of clustering genomic loci into contiguous regions of constant copy number, is often used for this noise correction in CNV data. This allows aggregation of numerous sequential read depth observations and aids in overcoming the inherent noise in the data. There have been a number of segmentation methods developed to tackle this issue in both next-generation sequencing and array comparative genomic hybridization data (***Olshen et al., 2004***; ***Nilsen et al., 2012***; ***Xi et al., 2016***). Circular binary segmentation (CBS) was introduced in 2004 and applied a “top-down” approach to CNV segmentation by iteratively introducing breakpoints into a CNV profile to explain changes in read depth, and this has been applied in numerous modern CNV software including Sequenza and CNVkit (***Olshen et al., 2004***; ***Favero et al., 2015***; ***Talevich et al., 2016***). Top-down segmentation was also used in the allele-specific piecewise constant fitting (ASPCF) algorithm used in ASCAT and PURPLE (***Van Loo et al., 2010***; ***Cameron et al., 2019***). Overall top-down segmentation approaches are overwhelmingly the method of choice for most CNV calling algorithms. Other methodologies have been applied, including a bottom-up approach in haar wavelet segmentation (***Ben-Yaacov and Eldar, 2008***) and hidden markov models (***Shah et al., 2006***), although these are used less frequently compared to CBS or ASPCF. Tools for somatic CNV detection tend to make errors considerably more often than small variant callers, owing to the stark difference in signal-to-noise ratios (***Zhang et al., 2019***). While highly sensitive CNV detection for identifying tumor suppressor deletions and oncogene amplifications is desirable, it is also vital to ensure that CNV segmentation is precise. Highly fragmented segmentation results preclude the use of metrics such as CNV count burden as phenotypic features related to CIN in cancer biology because false positive breakpoints cause an inflated estimate of the magnitude of CIN. Inherently noisy data impedes the ability to separate signal from noise, necessarily limiting the degree of positive, publishable findings that can be generated from CNV data (***201, 2019***).

Clinical genomics is a rising area of interest in cancer research. Studies that evaluate the utility of a personalized approach to identifying potential cancer therapeutics have been reported in recent years (***Robinson et al., 2017***; ***Pleasance et al., 2020***). Studies that broadly attempt to genomically characterize large cohorts of tumor biopsies are also becoming increasingly common-place (***Nik-Zainal et al., 2016***; ***Campbell et al., 2020***). As the cost of sequencing continues to decrease and large-scale sequencing studies of tumor biopsies become more commonplace, highly precise and accurate tools to interrogate these genomes will be required for their effective interpretation.

Here we present Ploidetect, a tool that performs copy number variation analysis of cancer genomes. Ploidetect contributes to the field of somatic CNV detection by using a novel segmentation approach to better segment genomes compared to contemporary methods. Compared to typical top-down segmentation algorithms which identify where to place breakpoints into a CNV profile, Ploidetect performs copy number segmentation using a bottom-up, subclone-aware iterative segmentation algorithm which tackles different challenges in CNV segmentation separately. We exhaustively benchmarked Ploidetect against contemporary CNV callers using first synthetic tumors across the full spectrum of tumor purity, real cancer cell line data subjected to synthetic dilution to a full spectrum of tumor purities, and finally a cohort of several hundred real-world metastatic tumor biopsies (***Laskin et al., 2015***; ***Pleasance et al., 2020***). This benchmark demonstrates that Ploidetect is able to detect focal copy number variants at all levels of tumor purity in a manner with greater sensitivity and specificity than any other method. Using the cohort of 687 metastatic tumor biopsies from patients with treatment-resistant cancers sequenced as part of the Personalized OncoGenomics (POG) program at BC Cancer (***Laskin et al., 2015***; ***Pleasance et al., 2020***), we demonstrated and propose a novel metric for the quantification of chromosomal instability that takes advantage of the accuracy of modern CNV software. We also identified patterns of homozygous deletions in these tumors congruent with prior studies, albeit using a smaller and more diverse cohort of tumors.

## Results

### Estimation of genomic copy number in cancer data

Ploidetect is comprised of a three step process to determine copy number in genomes: data preprocessing, purity/ploidy estimation, and finally copy number segmentation and estimation. Using depth of coverage of the normal, variable-width bins are generated such that an equal number of the normal’s mapped read bases are contained within each bin to account for variation in mappability. Next, tumor depth of coverage and *β*-allele frequency (BAF) data are calculated over these variable-width bins. These data are then used for tumor purity and ploidy estimation by Ploidetect.

Details of the purity/ploidy estimation, and copy number estimation are described in the supplemental methods. Briefly, depth of coverage and BAF data are segmented using a novel iterative approach with parameters intentionally set to ensure oversegmentation, rather than potential undersegmentation. These segments are used to infer useful statistics of the underlying coverage data, including the variance along segments. Using a series of Gaussian Mixture Models (GMMs), a number of models for explaining tumor purity and ploidy are tested and scored using numerous metrics including GMM likelihood and segmentation quality. CNV calling is performed using a coarse-to-fine approach; low-resolution data are segmented according to the purity and ploidy estimation, and the resulting segments are mapped to a higher-resolution binning of the same data. Because the only differences in CNV landscape expected at each iteration are focal CNVs smaller than the previous resolution, these can be identified by breaking segments at suspicious loci and re-applying the segmentation algorithm to repair falsely introduced breaks while retaining true focal CNVs. This procedure is repeated until completion or the contiguity of segmentation sharply declines between iterations, whereupon it is assumed that the limit of resolution has been reached and the previous iterations’ segments are returned (Figure 1). Once the segments are determined, allele-specific copy number estimation and zygosity estimation is performed on each segment using a GMM. Ploidetect has also been applied to perform CNV calling in single-sample contexts without a matched normal by using fixed-width bins of 5kb each, as well as using long read technology, such as from Oxford Nanopore instruments (Supplemental Figure 1) (***Boerkoel et al., 2022***), although we do not discuss this preliminary capability here.

**Figure 1.**
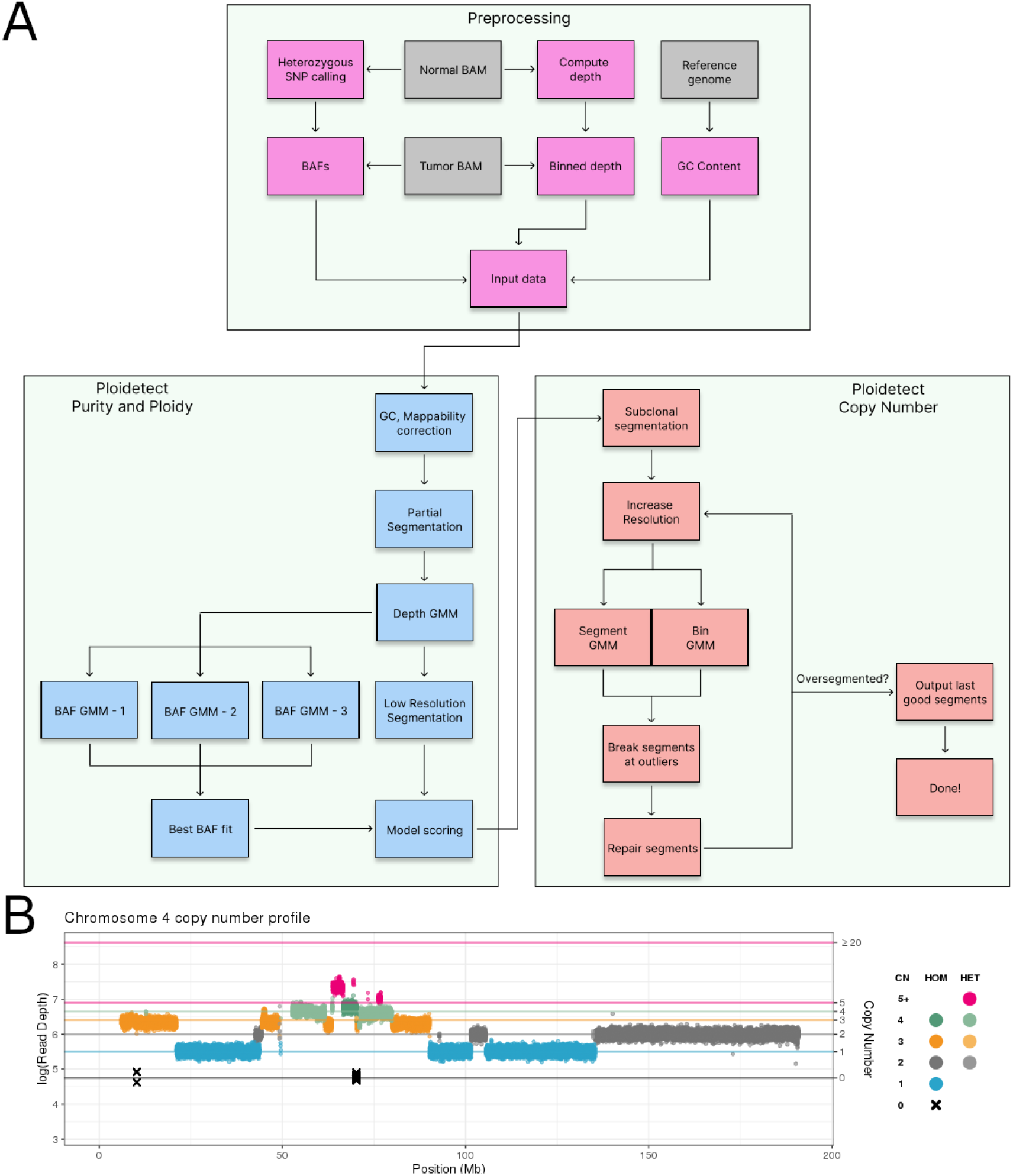
**A**. Outline of Ploidetect. Aligned sequence coverage data from tumor and matched normal are summarized into depth of coverage and beta-allele frequency (BAF) data, and GC content is computed from the reference genome to obtain input data. The input data are subjected to GC bias and mappability correction and subjected to partial low fidelity segmentation. Gaussian mixure models explaining the depth data are fitted to the data, and three GMMs describing the BAFs are also tested. The top scoring BAF model, describing the most likely ploidy is used alongside the depth GMM to score the model of tumor purity and ploidy. Once purity and ploidy are estimated, copy number estimation is performed. De novo segmentation in a subclone-aware manner is first performed in low resolution data to identify subclones with high prevalence. Data resolution are then increased, and bins are tested to identify potential focal CNVs followed by segment breakage and resegmentation. This process is continued until oversegmentation is detected and the algorithm completes. **B**. Example Ploidetect CNV outputs. Ploidetect was used to identify CNVs in a biopsy of a metastatic breast tumour from the POG cohort. CNV results from chromosome four are shown

### Synthetic tumor benchmarking

We assessed Ploidetect’s ability to identify CNVs in a number of experiments. We noted that published CNV data largely lacks gold standard CNV calls, which imposes difficulty in benchmarking using these data. We began by simulating five genomes at 80-fold tumor coverage and 40-fold matched normal coverage by introducing SNVs from NA12878 (***Zook et al., 2016***) as well as both chromosomal and segmental CNVs (Supplemental Table 1) in each simulated tumor and performed *in-silico* dilutions of tumor sequence data to generate aligned sequence data corresponding to tumors with simulated tumor purities between 10% and 100% (Supplemental Figure 2). To prevent biases, we generated the simulated CNV profiles without parameter tuning, and without post-hoc modification of the CNV profiles for the purpose of optimizing results. Using these simulated data, Ploidetect was benchmarked alongside PURPLE (***Cameron et al., 2019***), Battenberg (***Nik-Zainal et al., 2012***), Sequenza (***Favero et al., 2015***), HATCHet (***Zaccaria and Raphael, 2020***), ascatNGS (***Raine et al., 2016***), and CNVkit (***Talevich et al., 2016***) as contemporary CNV callers with similar functionality. Ploidetect performed comparably well compared to other methods at purity and ploidy estimation (Figure 2A-B). In terms of CNV breakpoint calling, Ploidetect routinely exhibited superior precision, recall, or both compared to other methods (Figure 2C). Across the spectrum of tumor purities, Ploidetect typically either had the highest performance or was comparable to ascatNGS (Figure 2D), and this observation was mirrored when assessed on a case-by-case basis (Figure 2E). Synthetic data are useful for providing benchmarks with a known gold standard, however realistic performance of software can rarely be fully appraised without using real biological data.

**Figure 2.**
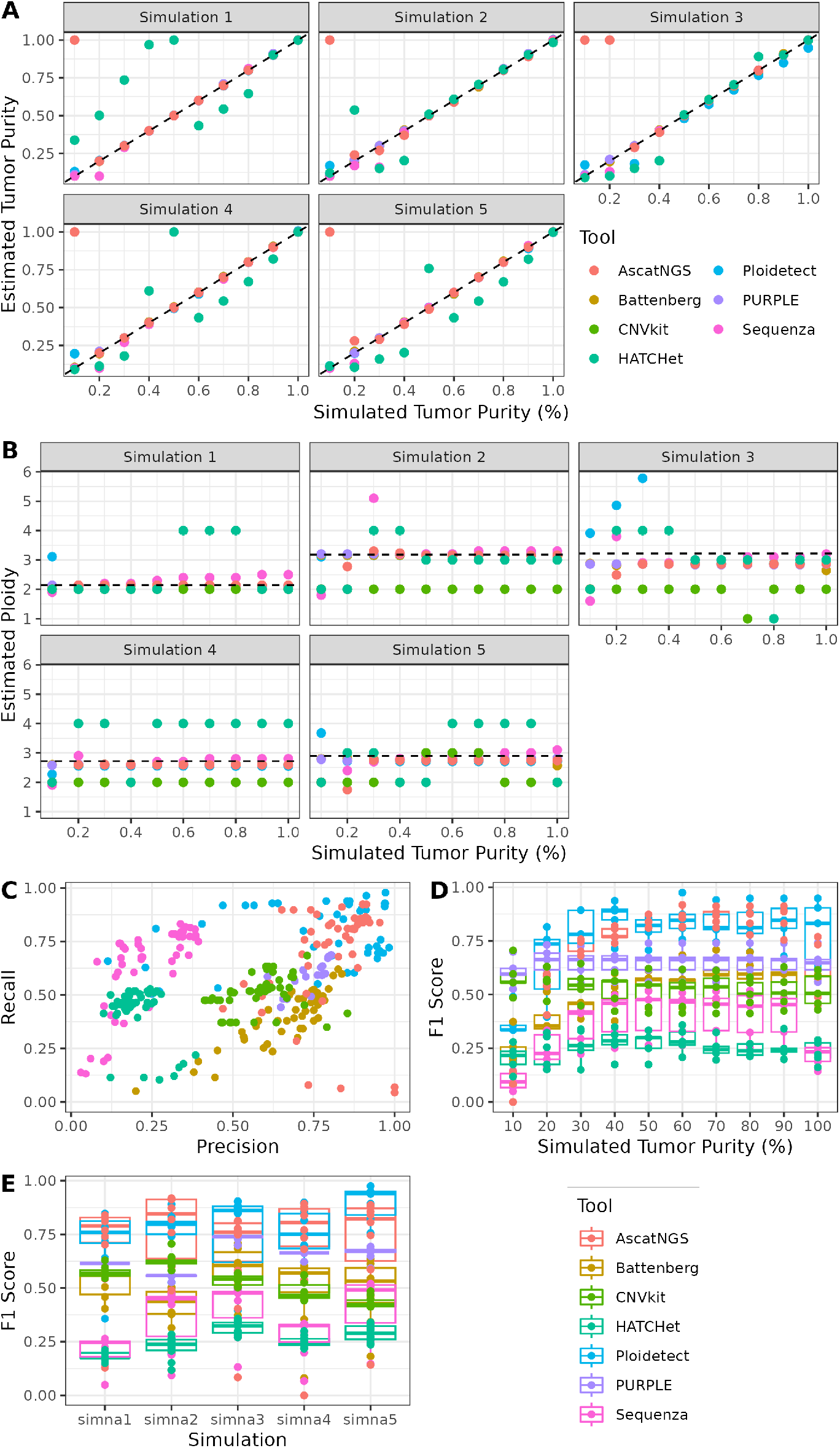
Synthetic tumor benchmarking. AscatNGS, Battenberg, CNVkit, HATCHet, Ploidetect, PURPLE, and Sequenza were used to determine CNVs in five synthetic tumor/normal genomes at ten different artificial tumor/normal mixture proportions. **A**. Tumor purity estimation across the five synthetic tumors. **B**. Tumor ploidy across the five synthetic tumors. **C**. Precision and recall of CNV calls from each synthetic tumor/normal admixture. **D**. F1-Scores of all synthetic tumors at each tumor/normal mixture purity. **E**. F1-Scores of all tumor/normal mixture purities for each synthetic tumor.

### Cell line benchmarking

We sequenced COLO829, an aneuploid skin carcinoma cell line along with its matched normal COLO829BL and performed in-silico dilution of the sequence data as before to obtain a set of COLO829 data at a range of tumor purities between 10% and 100%. Contrasting the synthetic data results, all of PURPLE, Battenberg, Sequenza and ascatNGS struggled to consistently model the tumor purity and ploidy of COLO829, whereas Ploidetect correctly modeled these features in all cases (Figure 3A-B). CNVkit does not estimate tumor purity and ploidy, and we were unsuccessful in running HATCHet on our COLO829 data without errors.

**Figure 3.**
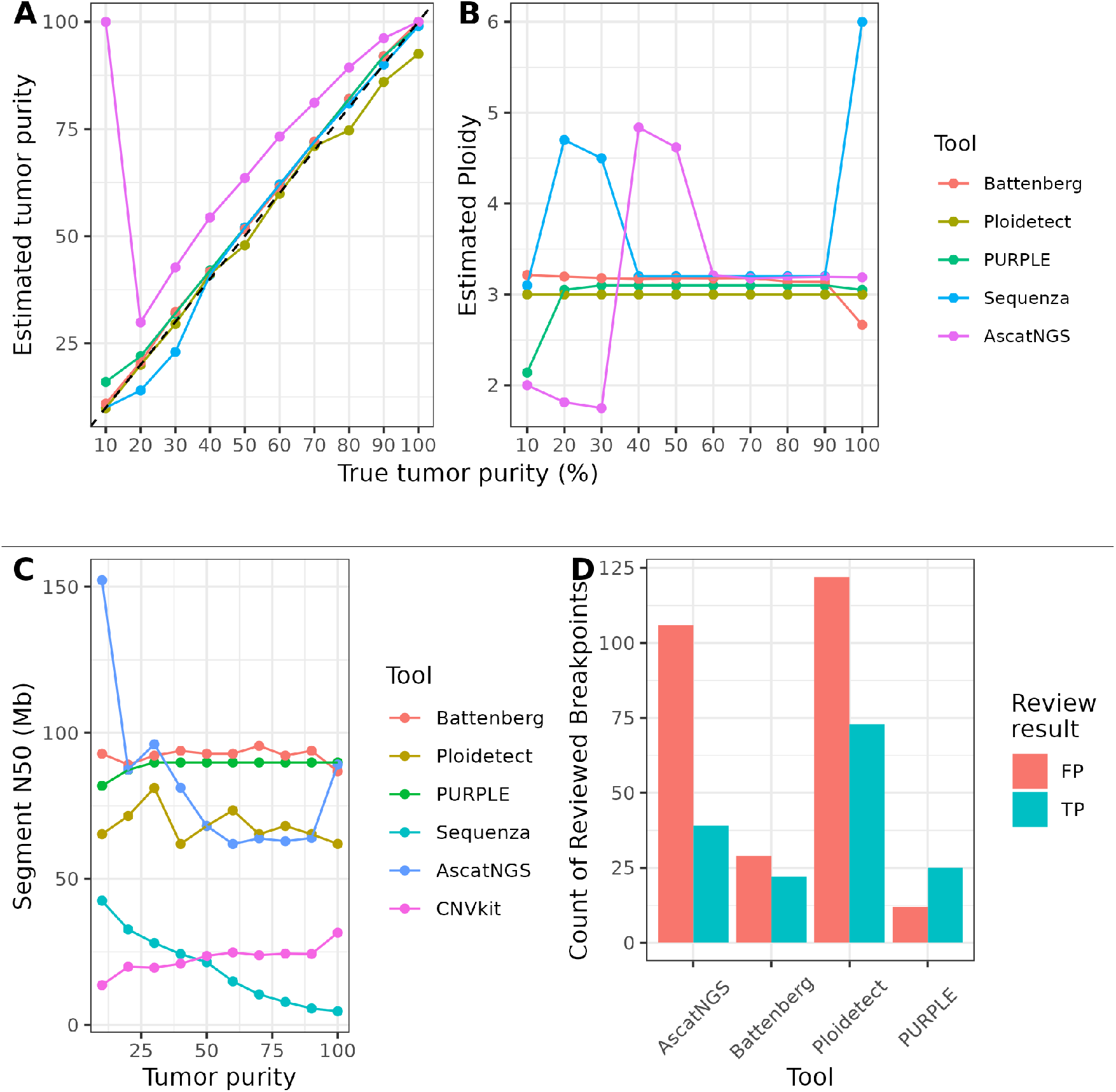
WGS data from the COLO829 cell line were mixed with matched normal COLO829BL at known proportions. AscatNGS, Battenberg, CNVkit, Ploidetect, PURPLE, and Sequenza were used to identify CNVs within each mixture. **A**. Tumor purity estimation error across all *in-silico* tumor/normal mixtures of COLO829 with COLO829BL for each tool which estimates tumor purity. **B**. Tumor ploidy estimation across all *in-silico* tumor/normal mixtures of COLO829 with COLO829BL for each tool which estimates tumor ploidy. COLO829 is known to be triploid (***Craig et al., 2016***). **C**. Contiguity of each software’s CNV profile for each COLO829 tumor/normal mixture. **D**. Results of manual review of AscatNGS, Battenberg, Ploidetect, and PURPLE CNV breakpoints in COLO829. FP = False Positive, TP = True Positive.

Benchmarking of CNV calling in real biological data is considerably more difficult due to a lack of gold standard call sets. Lacking gold standard calls, when comparing CNV profiles obtained from different software for the same underlying biological data, the ideal CNV profile is the one which recalls all relevant CNVs while also minimizing oversegmentation. The excessive introduction of segment breakpoints to model a CNV landscape is known as oversegmentation and results in reduced precision of CNV calls, as many calls are false positives introduced by the segmentation algorithm. Oversegmentation of CNV profiles allows for the identification of focal CNV events with high sensitivity, as even small changes in sequence data, whether the result of noise or a real CNV results in a breakpoint call. However, the consequent excess of segment breakpoints arising from this interferes with analysis of CIN as well as causing a mislabeling of genomic loci with an incorrect copy number. We used the N50 of segment lengths to measure the overall contiguity of a CNV profile. Oversegmentation of a CNV profile impacts the measured segment N50 by causing more of the genome to be contained within smaller segments. Focal CNVs, as commonly found in cancer also reduce the segment N50 because the otherwise long, “quiet” genomic segments are broken up by focal events. We surmised that ideally, a CNV caller should simultaneously maximize the segment N50 while also maximizing recall of CNV, two measures that would each affect the other inversely. We found that Ploidetect consistently provided more contiguous CNV calls than Sequenza and CNVkit, was comparable with AscatNGS, and its calls were consistently less contiguous than PURPLE or Battenberg (Figure 3C). In order to ascertain the ability of the callers to accurately recall CNVs, we performed manual review of raw alignment data corresponding to every breakpoint called by AscatNGS, Battenberg, PURPLE, and Ploidetect in COLO829 (Supplemental Table 2). CNVkit and Sequenza were not considered as each tool alone yielded multiple times more CNV breakpoints than the four selected tools combined. We found that overall, all CNV callers except PURPLE returned more false positive calls than true positive calls, and PURPLE yielded few calls overall (Figure 3D). However, Ploidetect’s results yielded nearly twice as many true positive calls compared to the second highest, ascatNGS. Furthermore, homozygous deletion of the tumor suppressors PTEN and CNOT1 were detected only by Ploidetect (Supplemental Table 2).

### Detecting CNVs in advanced cancer biopsies

While cell line data are valuable for benchmarking due to the high degree of control that may be exerted over experimental parameters such as tumor purity, there is no replacement for real-world data collected under realistic conditions. We sought to leverage the hundreds of sequenced pretreated metastatic cancer genomes collected as part of the POG project to assess the performance of the CNV callers. We used a conservative set of genes known to be recurrently affected by CNV in cancer (Supplemental Table 3) to measure recall in POG, and continued using the segment N50 to compare contiguity of segmentation. In POG, we found that Ploidetect typically identified a higher proportion of cancer-related CNVs per case than other methods while maintaining comparable genome-wide contiguity (Figure 4A). We recognized that simply assuming that all cancer-related gene CNVs are true events was likely to cause error in our assessment of results, and as a result we conducted a blinded manual review of the CNV data to estimate the accuracy of each tool for identifying CNV calls within metastatic tumors. We randomly selected 100 (out of 547) unique homozygous deletions and 100 (out of 768) amplifications of the cancer genes and manually reviewed the data to judge whether each mutation was a false positive or true positive call according to a predetermined manual review algorithm (Supplemental Figure 3) (Supplemental Table 4). We recruited a second expert bioinformatician to perform a blind audit of 50 of these calls, and the two reviewers were found to be in exact agreement (Supplemental Table 5). The results of the review were mapped to the CNV callset to estimate the false positive rate of each tool. In addition to being one of the most sensitive tools evaluated for identification of cancer-related CNVs, Ploidetect also had the highest accuracy for both amplifications and homozygous deletions (Figure 4B).

**Figure 4.**
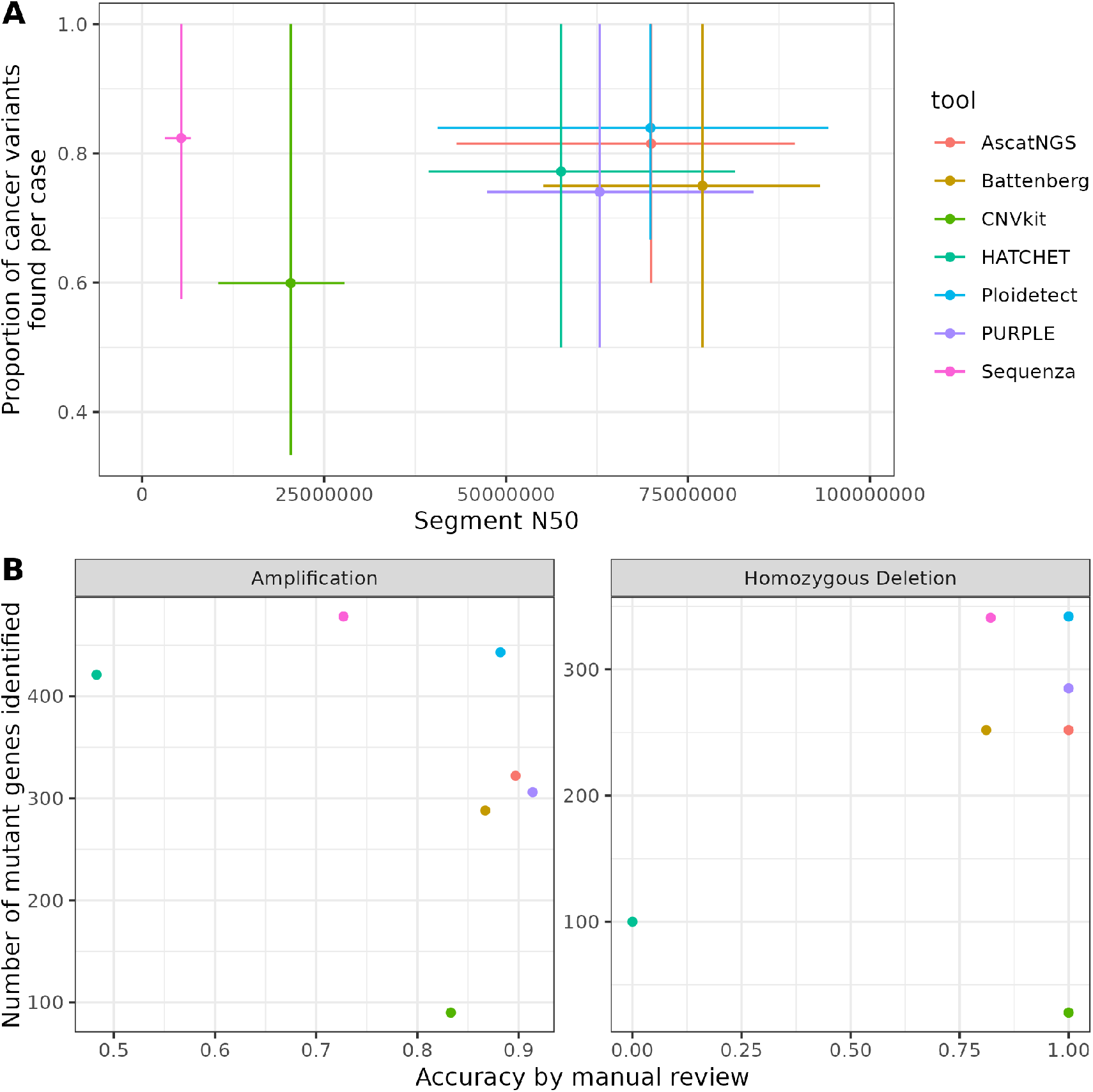
Ploidetect accurately identifies CNVs in metastatic cancer biopsies **A**. For each tumor, the gene-level homozygous deletions and amplifications in cancer-related genes were compiled from all tools. The proportion of these mutations which were detected by each individual tool are depicted on the Y-axis. Segment N50 of each tool’s CNV calls for each tumor are shown on the X-axis. Bars indicate interquartile ranges. **B**. Total number of cancer-related CNVs in POG by each tool are plotted against the assessed accuracy of cancer gene CNV calls by manual review.

### Pan-cancer analysis of homozygous deletions

Ploidetect was used to perform a pan-cancer analysis of homozygous deletions in POG. Of the 687 cases assessed, 501 harbored at least one homozygous deletion with a median length of 9628 base pairs, and an average of 2.05Mb of homozygously deleted sequence per case. Using a similar approach to Cheng *et al*. (***Cheng et al., 2017***), We identified 60 recurrently deleted regions (RDRs) which were recurrently homozygously deleted in POG, encompassing 282 genes (Figure 5A) (Supplemental Table 6) (FDR < 0.01, # cases ≥ 5). Overall, about 27.1Mb of the genome was identified to be recurrently deleted, with a median size per RDR of 64.4kb, minimum size of 7.6Kb and a maximum size of 9.7Mb. While known tumor suppressor genes such as CDKN2A and PTEN were typically entirely covered by RDRs, genes described as fragile site genes or fragile tumor suppressors in the literature such as WWOX (***Zanesi et al., 2011***) typically only had a fraction of their exons recurrently deleted in POG (Figure 5B).

**Figure 5.**
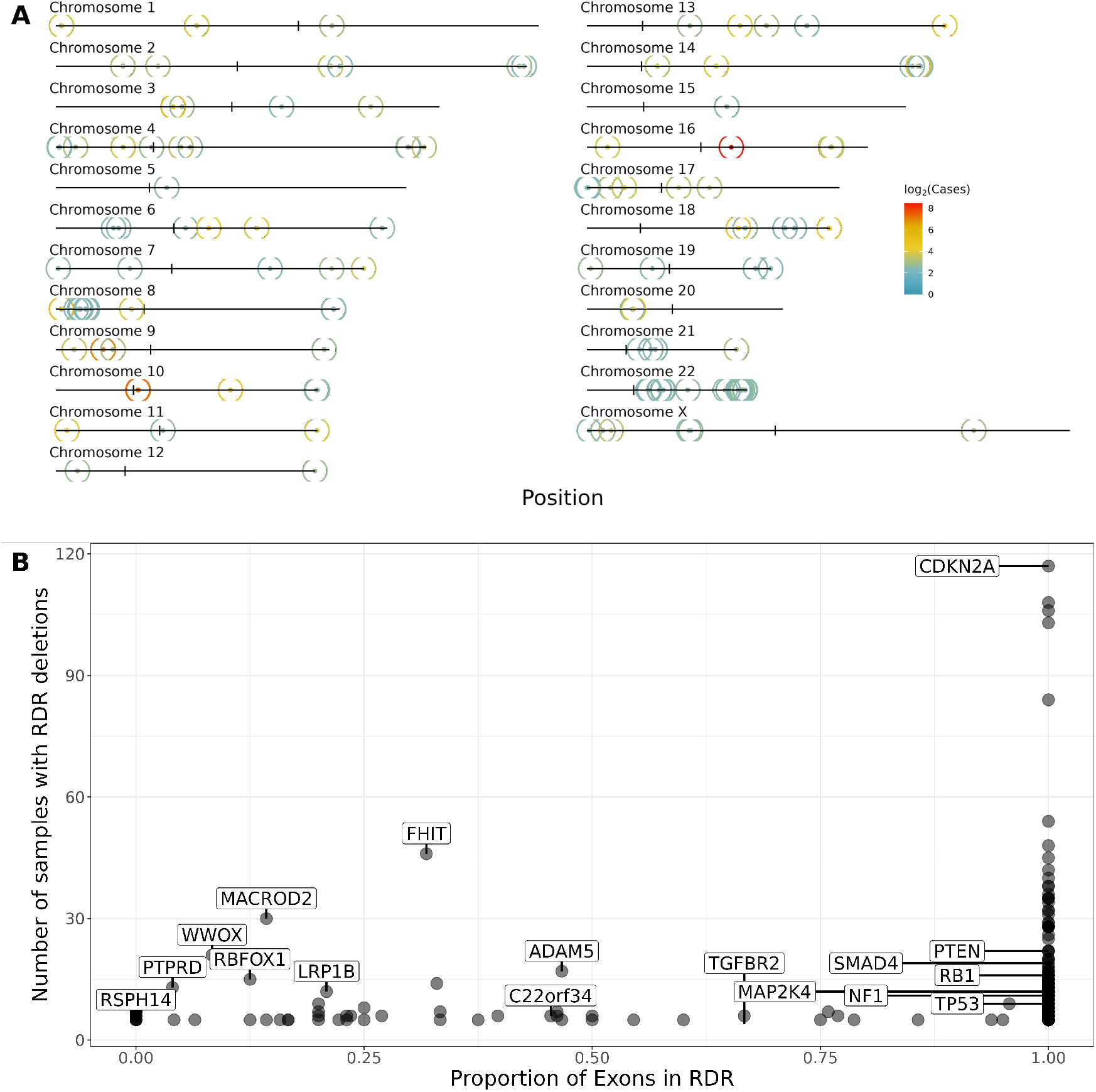
Recurrent homozygous deletions in the metastatic cancer genome. **A**. Positions of RDRs are indicated on each chromosome with a circle. Colors indicate the number of cases with deletions contributing to that RDR. **B**. For each gene within an RDR, the number of cases with a deletion in the gene were calculated. These values are plotted against the proportion of exons for that gene which were affected by any RDR. Selected genes with annotated oncological properties were manually labeled.

### The N50 of CNV segment lengths is an effective estimator of CIN

We next sought to characterize methods by which CIN could be measured in tumors. Previously, metrics such as the percent genome altered (PGA) and HRDscore have been used to describe CIN (***Sipos et al., 2021***; ***Przybytkowski et al., 2014***; ***Do Canto et al., 2019***; ***Hieronymus et al., 2014, 2018***; ***Bakhoum et al., 2018***). We noted that the PGA could characterize CIN acceptably well in some cases although some patterns of CNVs in tumors resulted in CNV landscapes with a very low PGA despite harboring multiple CNVs (Supplemental Figure 4). To address this, we investigated applying the N50 of CNV segment lengths. While PGA describes the whole-genome aggregate of all regions which are not copy neutral, the segment N50 measures the length of the longest segment where 50% of loci in the genome are contained within segments of that length or smaller. We assessed methods of measuring CIN by comparing their predictive ability for known CIN-causing mutations, namely in the TP53 and BRCA genes (***Dalton et al., 2010***; ***Ban et al., 2001***; ***Foijer et al., 2014***; ***Tutt et al., 1999***). The total breakpoint count, PGA, HRDscore, and segment N50 were compared between mutant and wild-type cases for TP53 and the two BRCA genes. For the TP53 case, all metrics were significantly associated with mutation status, and the segment N50 had the highest effect size of all metrics tested (p < 0.05) (Figure 6A) (Supplemental Figure 5). Expectedly, the HRDscore effect size in the BRCA mutant case was the highest among all metrics, although the segment N50 was comparable, and all metrics but PGA were significant (p < 0.05) (Figure 6B). Since the segment N50 appeared to be a strong predictor of CIN by this experiment, we next attempted to identify other genes whose mutations were associated with CIN.

**Figure 6.**
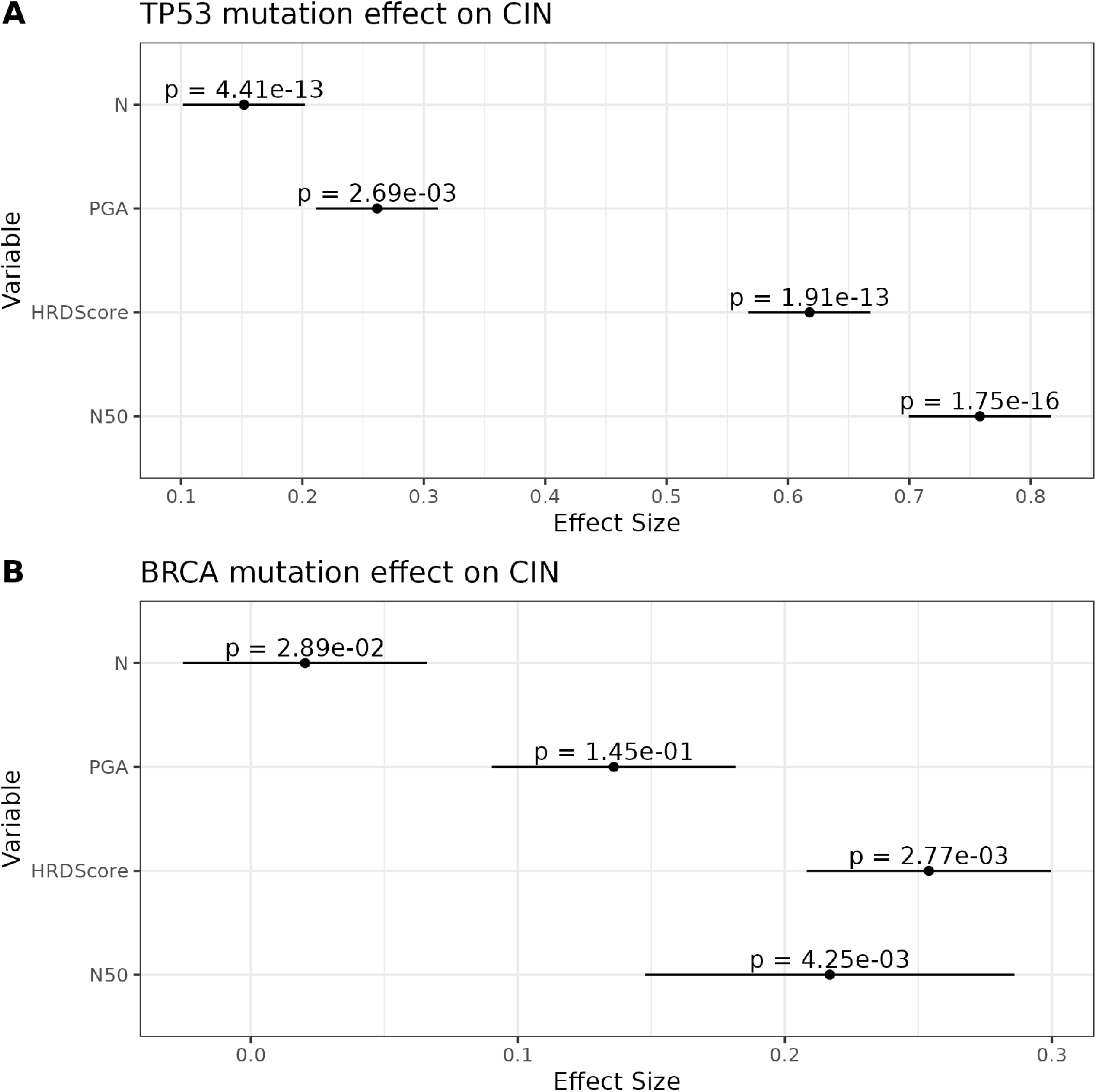
Segment N50 predicts CIN driver mutation status. Total breakpoint count (N), PGA, HRDScore and the segment N50 were compared for different CIN driver genes’ mutational status. Wilcoxon rank-sum test was used for significance. **A**. Effect sizes and p-values for TP53 mutant and wild-type states. **B**. Effect sizes and p-values for BRCA mutant and wild-type states.

### CIN Gene Discovery

We next attempted to identify genes whose mutations are associated with altered levels of CIN. We used support vector machines to predict segment N50 from mutation status for all genes with at least 10 cases with mutations in POG, and applied recursive feature elimination with 5-fold cross validation to select for a set of genes which provided the best predictive ability. We tested the resulting set of gene mutations for putative association with segment N50. We obtained 28 candidate genes whose mutations were associated with the segment N50 (Wilcoxon rank-sum test, Benjamini-Hochberg (***Benjamini and Hochberg, 1995***) false discovery rate < 0.2) (Table 1). Predictably, TP53 was observed to be significantly associated with reduced segment N50. The large genes encoding Titin and Giantin were identified as well, likely due to the underlying negative correlation between tumor mutational burden and segment N50 (Supplemental Figure 6). ROCK2 has previously been implicated in tumorigenesis and chromosomal integrity (***Lingy et al., 2015***; ***Oku et al., 2014***; ***Libanje et al., 2019***), and we identified ROCK2 mutations as significantly associated with a reduction in the segment N50 (Figure 7A). CHD4 mutations were associated with an increase in chromosomal stability (Figure 7B), and CHD4 has been reported to DNA-binding helicase activity as well as oncogenic activity (***Xia et al., 2017***; ***Oyama et al., 2021***; ***Arends et al., 2019***).

**Table 1.** List of genes with CIN-associated mutation status. Mutation statuses of all genes with 10 or more instances of somatic mutation in POG were used to predict segment N50 using support vector machines. Recursive feature elimination selected for a set of genes whose mutation status was most consistently predictive of the segment N50. Candidate genes were tested for association with segment N50.

| Gene ID | Coefficient | Nominal p | FDR |
| --- | --- | --- | --- |
| TP53 | -0.116891589 | 1.99575E-14 | 5.28873E-12 |
| AC074391.1 | -0.128563905 | 1.27325E-10 | 1.68706E-08 |
| TTN | -0.090456545 | 3.75897E-06 | 0.000332042 |
| ROCK2 | -0.266465237 | 0.00024803 | 0.016431955 |
| ZNF804B | -0.227724308 | 0.000743941 | 0.039428861 |
| IFT172 | -0.172811964 | 0.001206398 | 0.053282586 |
| ATP2B2 | -0.097852839 | 0.003013079 | 0.11312294 |
| DMBT1 | 0.345505448 | 0.003858758 | 0.11312294 |
| KAT6B | 0.105515244 | 0.003921761 | 0.11312294 |
| AKD1 | -0.148617727 | 0.00426879 | 0.11312294 |
| PCDH15 | -0.118837568 | 0.005302926 | 0.11884362 |
| PLCB1 | -0.27359427 | 0.005990587 | 0.11884362 |
| ODZ2 | -0.165090589 | 0.006350721 | 0.11884362 |
| MGAM | 0.153987884 | 0.006678628 | 0.11884362 |
| ANK2 | -0.131762504 | 0.006726997 | 0.11884362 |
| LRP1B | -0.078111484 | 0.008081839 | 0.132861023 |
| ZNF451 | -0.318875833 | 0.00852316 | 0.132861023 |
| COL12A1 | 0.146496448 | 0.009264726 | 0.136397353 |
| EPPK1 | -0.103690607 | 0.011135097 | 0.155305306 |
| CENPF | -0.147681442 | 0.012614446 | 0.162090186 |
| CNTN6 | -0.087261152 | 0.013611958 | 0.162090186 |
| MYO15A | 0.09770158 | 0.013843886 | 0.162090186 |
| CHD4 | 0.197767058 | 0.014303565 | 0.162090186 |
| PXDN | -0.192456299 | 0.015471793 | 0.162090186 |
| GOLGB1 | -0.179000736 | 0.015863945 | 0.162090186 |
| OBSCN | -0.086120287 | 0.016262484 | 0.162090186 |
| PTEN | 0.113716991 | 0.016514849 | 0.162090186 |
| ANO4 | -0.090204324 | 0.017417399 | 0.164843238 |

**Figure 7.**
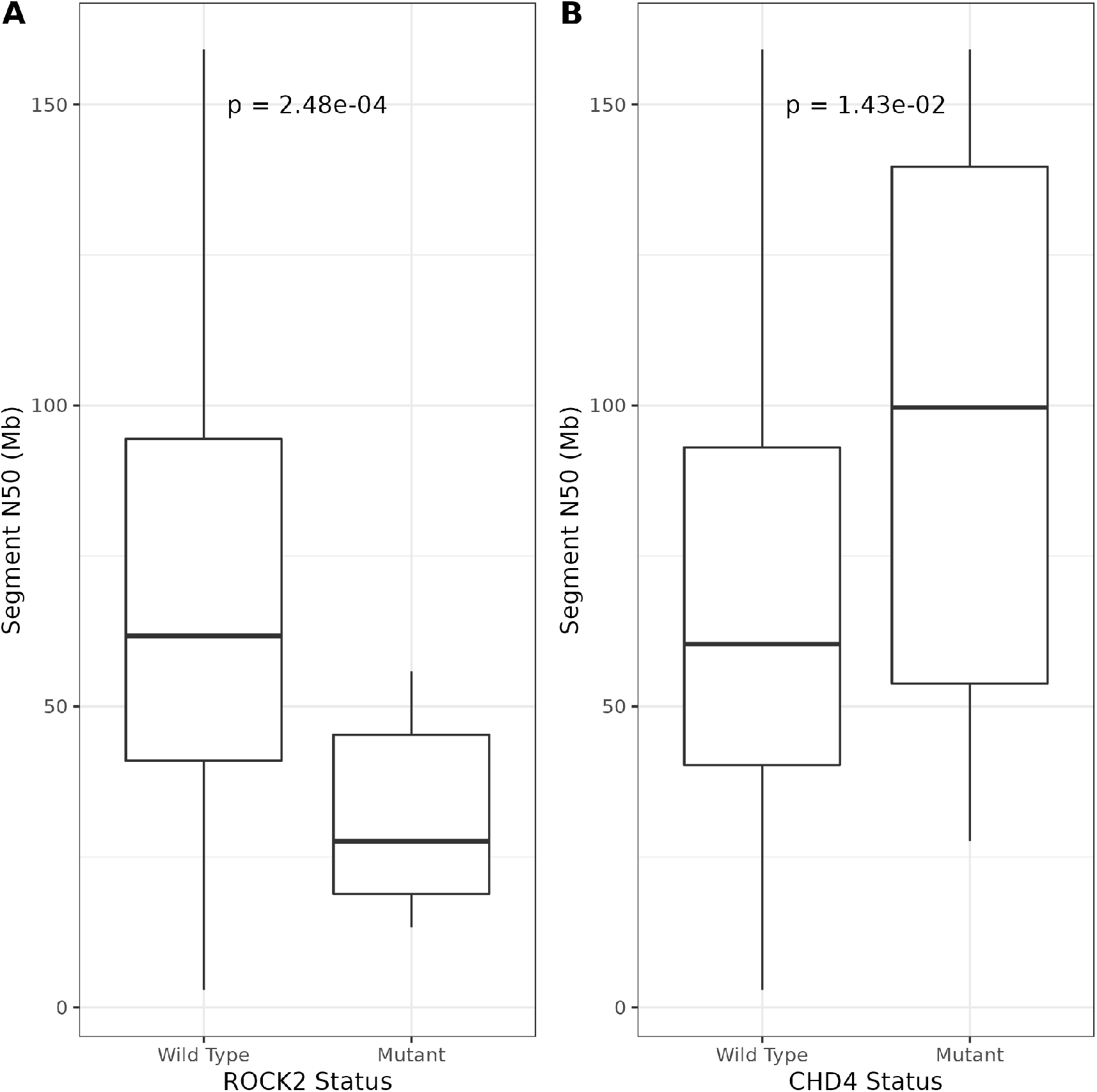
Gene mutations associated with altered CIN. Candidate genes were identified by using RFE to select for genes whose mutation status could predict CIN. Significance testing was performed on segment N50 values between mutant and wild-type groups using the Wilcoxon rank-sum test. **A**. Comparison of segment N50 in ROCK2 mutant and wild-type tumors in POG. **B**. Comparison of segment N50 in CHD4 mutant and wild-type tumors in POG.

## Discussion

The modern application of CNV analysis is principally focused on two different problems; identification of specific, focal CNV events within individual tumors, such as clinical biopsies (***Campbell et al., 2020***; ***Pleasance et al., 2020***), and pan-cancer patterns of CNV such as CIN, inferred from bulk CNV profiles and CNV size distributions (***Macintyre et al., 2018***; ***Steele et al., 2022***). For each of these cases software is required which faithfully models the underlying CNV landscape of a tumor while also identifying focal, impactful mutations that arise within a patient’s specific disease. We have introduced Ploidetect, a software package which enables CNV analysis in a manner more sensitive and precise than previously possible. We illustrated this by leveraging Ploidetect’s strengths in two key areas - firstly by identifying candidate genes involved in the maintenance and abrogation of chromosomal stability, and secondly in the pan-cancer assessment of homozygous deletion within metastatic cancers.

Ploidetect maximizes sensitivity for identifying CNVs while maintaining excellent contiguity. Analysis of the prevalence of CNVs within biological samples is highly dependant on the chosen software’s ability to identify the events themselves. Within tumor cell lines at the full spectrum of possible tumor purities and within real metastatic tumor biopsies, Ploidetect identified a greater proportion of cancer-related CNVs than any other tool assessed. The reliance on gaussian mixture models (GMMs) by Ploidetect allows for the modeling of the CNV landscape to be flexible to a variety of different conditions without the need for manual parameter tuning. Parameters commonly specified by the user in other methods, such as segmentation thresholds (***Nilsen et al., 2012***) are automatically determined from the data, rather than being set by the user. This allows for a naive application of Ploidetect to perform competitively or better than other software in large, diverse cohorts such as POG, since the algorithm adapts to each individual patient, rather than needing to be adapted ad-hoc. Ploidetect’s bottom-up, coarse-to-fine approach to CNV estimation conceptually allows for the compartmentalization of different tasks in the CNV calling process. Low-resolution data cannot identify focal CNVs, however detection of focal CNVs is not expected nor necessary in the initial segmentation passes. Rather, the initial segmentation iterations focus on identifying high confidence larger CNV segments where the only potential remaining CNVs are extremely small. Further iterations with finer resolution are then only concerned with the identification of focal CNVs which were too small for previous iterations’ resolution.

In our study, we introduced a metric for benchmarking of CNV results - the segment length N50. Oversegmentation is an overall difficult phenomenon to quantify using hard metrics such as precision, as an individual tumor with an oversegmented CNV profile may contain thousands of false CNV calls. Consequently “softer” metrics such as the N50 provide a useful method for assessing broad trends in this phenomenon. For this purpose, the N50 provides a useful metric that ideally should be as high as possible so long as the high contiguity does not come at the cost of missing real CNVs. Within COLO829, for instance, we found that Ploidetect did not provide the highest segment N50 among the tools tested, however it successfully identified the largest number of manually curated CNV breakpoints in COLO829 among the assessed tools by far. Similarly, in POG, we found that Ploidetect had a competitive segment N50 compared to other tools while maintaining a greater recall of common cancer CNVs. While Ploidetect’s performance was overall quite promising, we noted were some circumstances where we found that Ploidetect struggled to provide optimal results.

It is useful in the presentation of a novel method to also discuss its weaknesses and shortcomings. We found that one key area where Ploidetect would struggle was in the segmentation of longer stretches of amplified DNA, likely due to read depth observations “drifting” from their expected values as their copy number increases further beyond the values near ploidy. Furthermore, poisson-distributed counts data exhibit greater variance as the mean of counts increases, resulting in amplified segments having a larger variance than the model would suggest. We would therefore advise potential users to use care in the interpretation of Ploidetect’s CNV calls in these circumstances, for instance in the relatively common amplifications of 8q (***IBRAGIMOVA et al., 2021***), as these types of events commonly resulted in oversegmentation in our experience. Finally, we note that the performance of CNV calling is highly sensitive on accurate estimation of the tumor purity and ploidy. While we have demonstrated that Ploidetect estimates these parameters with a high degree of correctness, it, like all software, is not infallible and errors in purity and ploidy estimation can drastically impact CNV estimation.

We identified numerous recurrently homozygously deleted regions within metastatic tumors using Ploidetect. Although our cohort was a fraction of the size of Cheng *et al*.’s analysis cohort, we uncovered many of the same patterns of mutation. While it is not a particular challenge to identify CDKN2A, we also successfully identified, in a pan-cancer context, recurrent homozygous deletion of tumor suppressor genes with lower frequencies of homozygous deletion such as MAP2K4 and PTPRD, which were each only detected in about 0.2% of primary tumors previously (***Cheng et al., 2017***), but detected in POG at a nearly 10-fold higher frequency. Without further investigation using Ploidetect in the primary cancer context, we cannot conclude whether these differences are the result of increased mutational burden in metastatic tumors, or because of a difference in sensitivity between Ploidetect in WGS and ASCAT in SNP array data. Nevertheless, we have demonstrated that Ploidetect is capable of providing sensitive, biologically meaningful CNV calls in the metastatic cancer context.

Studies of CIN and patterns of CNV, including mutational signatures rely less on sensitive CNV calling, but moreso on accurate and contiguous CNV calls. Both published works to date that examined signatures of CNV within cancer utilized features of CNV segments, such as the size distributions of segments at different copy number states, as well as other variables such as the length of oscillating CNV states (***Macintyre et al., 2018***; ***Steele et al., 2022***). Fragmented CNVs from oversegmentation cause an overestimation of CIN, and a mischaracterization of distributions such as segment sizes commonly used in CNV signature analysis. Other specific features of CNV profiles, such as oscillating CNV length as used by Macintyre *et al*. are particularly susceptible to oversegmentation as those errors commonly manifest as focal CNV changes by a single copy number. We noticed that examination of oversegmentation is rarely performed in the introduction of CNV software (***Nik-Zainal et al., 2012***; ***Priestley et al., 2019***; ***Favero et al., 2015***), which is of particular concern when these types of errors have the potential to distort downstream analyses by a large degree.

Novel metrics for the measurement of biological features enable leaps in our understanding of those features. HRDScore and HRDetect allowed for the degree of a genome’s homologous recombination deficiency to be summarized into a single score (***Zhao et al., 2017***; ***Melinda et al., 2016***), and PGA has been used to describe the overall degree of CIN within tumors for years (***Hieronymus et al., 2014, 2018***; ***Bakhoum et al., 2018***). Comparing methods for characterization of CIN is difficult; how does one measure a phenotype that is largely defined using the metric intended to measure it? It is well understood both mechanistically and empirically that mutations in TP53 and the BRCA genes cause CIN (***Dalton et al., 2010***; ***Ban et al., 2001***; ***Foijer et al., 2014***; ***Tutt et al., 1999***), providing a means to label tumors which are likely to be chromosomally unstable. The segment N50, measuring the typical length of DNA unaffected by CNV is a rational metric for measuring the degree of CIN within a genome, and in POG was shown to best predict the mutation status of TP53. HRDscore is narrowly defined for the quantification of a specific etiology of CIN, and consequently it is of no surprise that BRCA gene mutation status had a strong and consistent effect on HRDscore, although the segment N50 still performed well. The PGA evidently did not estimate CIN to the same degree as either the HRDscore or the segment N50, likely because the bulk quantity of CNV-affected DNA does not reflect the actual structural landscape of the tumor. For example, HRD-induced chromosomal instability has been characterized in part by the accumulation of small CNVs that do not impact a large overall proportion of the genome (***Macintyre et al., 2018***). PGA necessarily struggles to characterize CIN induced by BRCA1/BRCA2 mutation, likely because HRD often causes numerous focal CNVs, which does not largely impact the singular feature that PGA measures - total altered DNA (***Macintyre et al., 2018***). One advantage of the PGA is that by essentially disregarding focal CNVs, the metric is theoretically robust to oversegmentation events. Modern CNV callers, including Ploidetect, Battenberg, ascatNGS, and PURPLE are all demonstrably capable of minimizing oversegmentation to a high enough degree that the use of contiguity to directly measure CIN is now possible. Overall the segment N50 is a rational, easily understood, easily computable, and potentially useful metric for the summarizing of chromosomal instability within a tumor.

The utility of novel approaches for performing CNV research will allow for advances in this field. Already, Ploidetect has been used in other studies tumor purity and ploidy estimation (***Gagliardi et al., 2020***), as well as germline CNV calling using oxford nanopore data (***Boerkoel et al., 2022***). Further application and development of improved CNV calling methods are likely to yield novel insights into the underlying biology of CNVs and cancer. Advancement in our understanding of how CIN manifests and can be measured will enable even further insights in this regard.

## Methods and Materials

### Methods

#### Sample collection, sequencing and POG analysis

Samples were obtained from the Personalized OncoGenomics (POG) project, with inclusion criteria, enrolment details, clinical background, and specific details on sequencing methodology, analysis, and software previously described (***Pleasance et al., 2020***). Briefly, POG tumor biopsies are obtained from advanced, treatment-resistant metastatic cancer patients. The biopsies are sequenced using short-read paired-end sequencing technology to approximately 80-fold genomic coverage, with a matched germline genome from blood sequenced to approximately 40-fold coverage. Expression data are also obtained using RNA-seq with a target depth of 200 million paired-end reads. Data from 687 total patients were used. Binary alignment map files were procesed using samtools 1.11 (***Danecek et al., 2021***), BCFtools 1.11(***Danecek et al., 2021***), and BEDtools 2.30 (***Quinlan and Hall, 2010***). SNVs, isoform-specific expression, and clinical data were processed as described previously and accessed from our internal database, vardb (Lewis et al., in preparation.). COLO829 and COLO829BL cell lines were prepared according to the “GSC” methods subsection of Craig *et al*., (***Craig et al., 2016***). COLO829 dilutions were prepared by mixing COLO829 tumor read pairs with COLO829BL read pairs at ploidy-adjusted ratios to produce a final tumor cell purity of 10-100% at 80-fold coverage. COLO829BL sequence was used at 40-fold coverage.

#### Ploidetect

The Ploidetect method is described in detail in the Appendix.

#### Statistical analysis

Statistical tests were the two-sided Wilcoxon rank-sum test unless otherwise specified. Analyses were performed in R version 3.6.3.

#### Cancer genome simulation

We simulated five genomes with somatic copy number alteration. Using the hg19 version of the human reference genome, we generated two haplotypes for the matched normal. We introduced the SNPs of NA12878 into hg19, with heterozygous variants being introduced into haplotype A and homozygous variants introduced into both haplotypes. We next introduced a custom set of segmental CNVs (CNVs affecting less than a whole chromosome) into each genome using simuG (***Yue and Liti, 2019***), and introduced chromosomal aneuploidies into the mutated genome by duplicating or deleting chromosomes with biopython 1.79 (***Cock et al., 2009***). The final CNV landscapes of each genome are shown in the supplemental data. Once the haplotypes were processed, they were concatenated into singular fasta files, ART 2.3.7 (***Huang et al., 2012***) was used to generate illumina HiSeq 2500-like 150bp paired-end sequence reads from the genomes (parameters: -ss HS25, -p, -m 500, -s 30), and the resulting fastq files were aligned to the hg19 reference genome using min-imap2 2.18-r1015 (***Li, 2018***). Because cherry-picking and parameter tuning are a common concern in studies which use synthetic data, we abstained from parameter tuning of software or modifying the synthetic genomes after the sequence data were generated for the first time without error. One exception was in simulation “sim4”; data entry of a CNV with an “end” position less than the “start” position resulted in the erroneous homozygous deletion of much of a chromosome arm, which is unrealistic in addition to being the result of a technical error (***Priestley et al., 2019***). This error was corrected by only altering the erroneously entered “end” position of the CNV. For each genome, we computed the tumor/normal sequence read proportions which would result in tumor purities of 10%, 20%, 30%, 40%, 50%, 60%, 70%, 80%, and 90% and mixed tumor and normal reads in those proportions to generate 50 total binary alignment map files corresponding to ten levels of tumor purity for each of five synthetic tumors.

#### Benchmarking

We compared Ploidetect with 3 other CNV tools, namely Sequenza (***Favero et al., 2015***), Battenberg (***Nik-Zainal et al., 2012***) and PURPLE (***Cameron et al., 2019***). Sequenza version 3.0.0, Battenberg version 1.1, ascatNGS version 4.2.1, HATCHet version 1.1.1, CNVkit version 0.9.9, and AMBER/COBALT/PURPLE versions 3.5/1.11/2.51, respectively. We tested using the 50 synthetic tumor/normal pairs described above, COLO829, also as described above, and 687 cases of the Personalized Oncogenomics (POG) cohort (***Pleasance et al., 2020***), including recent cases sequenced after the publication of Pleasance et al. COLO829 was sequenced as described in the “GSC” section of Craig *et al*. (***Craig et al., 2016***) Purity and ploidy results were obtained from each tool. Each tool was run using its respective default settings using a Snakemake (***Köster and Rahmann, 2012***) workflow. Homozygous deletions were defined as a copy number below 0.25, and amplifications were defined as a copy number ≥ ploidy + 3.

#### Manual Review

We performed a blinded manual review of 100 each randomly selected amplifications and homozygous deletions in POG according to our manual review algorithm (Supplemental Figure 3). One reviewer reviewed all 200 mutations, and a second reviewer reviewed the first 50 of 100 randomly selected mutations. After the first 50 mutations were reviewed by our second reviewer, we noticed that the inter-reviewer agreement was perfect and decided that additional review was extremely unlikely to discover systematic errors in the first reviewer’s results. Review of COLO829 breakpoints was conducted using IGV and breakpoint calls were concluded based on a qualitative balance of probabilities, taking into consideration the observed consistency of read mappings in the vicinity of the putative breakpoint, changes in sequencing coverage in the vicinity of each called breakpoint as well as discordant read pair mapping information.

#### Detection of significantly deleted regions

We performed a permutation analysis by shuffling the midpoint of all homozygous deletions in POG on each chromosome 10^7^ times. For each interval, we recorded the number of permutations where shuffled deletions were more numerous than actual deletions. P-values were obtained by dividing this count by the number of permutations overall. The p-values were adjusted for multiple testing using the Benjamini-Hochberg method (***Benjamini and Hochberg, 1995***). For analysis we considered regions with a false discovery rate ≤ 0.01 and which contained deletions in at least five separate samples.

#### Analysis of chromosomal instability

Segment N50 values were obtained from Ploidetect CNV segments by sorting segment lengths, computing the cumulative sum, and selecting the largest segment length whose index had a cumulative sum less than 50% of the sum of all segment lengths. HRDScores were obtained directly from POG analyses (***Pleasance et al., 2020***). PGA was computed by dividing the sum of the lengths of non-copy-neutral CNV segments by the sum of all CNV segment lengths. For CIN gene analysis, we obtained the set of all single nucleotide variants (SNVs) in POG (***Pleasance et al., 2020***) and removed all intergenic, synonymous, upstream, downstream, intronic, and untranslated region SNVs. We selected genes which had SNVs in at least 10 cases and assigned a binary variable for each case and gene combination, where 1 indicated an SNV was present in the gene, and a 0 indicated the absence. Segment N50 values were scaled by dividing each value by the maximum N50 observed. We performed recursive feature elimination (RFE) by training a support vector regression (SVR) model using scikit-learn (***Pedregosa et al., 2011***). We used a linear kernel, a C value of 1 and a gamma of 0.001. We eliminated the 10% least informative positively signed coefficients and the 10% least informative negatively signed coefficients, measured by the absolute value of the coefficients at each iteration. At each iteration, 5-fold cross validation was performed to assess the training and test error, measured as the root-mean-squared concatenated difference between predicted scaled N50 values and the real scaled N50 values for all folds. Using the set of genes from RFE with the lowest test error, we performed a one-sided Wilcoxon rank-sum test for each gene using their SVM coefficients as the prior to inform the directionality of each test. We tested the difference in segment N50 between variant and non-variant cases for each gene selected by RFE, and corrected the resulting P-values for multiple testing using the FDR method.

#### Ploidetect software availability

Ploidetect is implemented in R and made available freely at github.com/lculibrk/Ploidetect. Analyses in this paper were conducted using Ploidetect version 1.4.0.

## Acknowledgments

We would like to gratefully acknowledge the patients of the POG program and their families, without whom we would have never been able to conduct this study. We also gratefully thank the oncologists, nurses, analysts, and volunteers involved with POG. We thank the BC Cancer Foundation and Genome British Columbia for their support (project B20POG). We also acknowledge contributions towards equipment and infrastructure from Genome Canada and Genome British Columbia (projects 202SEQ, 212SEQ, 12002), Canada Foundation for Innovation (projects 20070, 30981, 30198, 33408 and 35444) and the BC Knowledge Development Fund. LC is supported by a Canadian Institutes for Health Research (CIHR) Frederick Banting and Charles Best Canada Graduate Scholarship, number GSD-164207 and a University of British Columbia 4-year fellowship. JG was supported by a University of British Columbia 4-year fellowship. SJMJ and MAM acknowledge funding from the Canadian Research Chairs Program.

## Availability of data and materials

Ploidetect is free and open source software, available at https://github.com/lculibrk/Ploidetect. POG data are available at the European genome-phenome archive (EGA), accession EGAS00001001159.

## Supplementary Methods

### Data preparation

Ploidetect utilizes sequence read information from paired tumor-normal samples. Read depth information is affected by biases in mappability and GC-content. Since many of these biases are closely linked to genomic position, we first generate a mappability-adjusted set of bins using the paired normal genome. Per-base read depth for high quality mapped reads in the normal genome is binned using cumulative sum threshold binning, excluding regions covering known repetitive elements identified by RepeatMasker (**?**), with a threshold of 100000 single-base coverage by default for paired-sequencing experiments which have roughly a 2:1 ratio of tumor genome coverage to normal genome coverage. For a 40x paired normal genome, this routinely results in bins of roughly 1-2 Kbp in size, depending on the read mappability. Total read coverage of the tumor and normal genomes is then counted over the resulting bins. To obtain *β*-allele frequencies (BAFs) we perform variant calling of the germline at loci used in the affymetrix 500K array spread throughout the genome and identify heterozygous SNPs occurring at these positions. The alleles at these positions within the matched tumor sample are then counted to yield the BAFs.

### Data prepossessing

Input data are first filtered to contain only non-centromeric regions from chromosomes 1 through 22 and the X chromosome. To reduce data complexity and improve the signal-to-noise ratio we merge adjacent bins into larger meta-bins of roughly 100kbp size by default, taking the sum of read depths from those bins. Read depths are corrected for residual GC content and mappability biases using the loess method, with a span of 0.75 for robustness to outlier values.

### Purity and Ploidy Estimation

To estimate the expected read depths and BAFs of CNV gains and losses, accurate estimates of tumor purity and ploidy must be obtained. Since the relationship between sequence read depth and copy number is linear, we model the read depth landscape using Gaussian mixture models (GMMs) whose component means are separated by the expected difference in read depth per copy number. Since different purities affect the differential read depth parameter, we fit a number of GMMs covering a range of differential read depths to the data.

To identify the most likely copy number represented by each GMM component, we overlay a second set of GMMs, each representing a different ploidy assumption. We observed empirically that tumors without notable regions of copy number two or lower are exceedingly rare, and accordingly we assume that the lowest well-represented (1% of genome) GMM component represents a copy number of 0, 1, or 2. For each possible genomic copy number and each ploidy assumption, we fit a GMM to the BAFs using predicted BAFs for the possible allele combinations as component means. Since we are interested in modeling copy number and not allele states, we take the top allele probability for each copy number as the copy number’s probability. The result is two models with identical dimensions. We take the inner product of the two models’ outputs and compute the maximum likelihood estimate of the copy number for each genomic bin. Since we computed one model for each possible ploidy state, we choose the ploidy state with the highest complete likelihood. Finally the tumor purity is calculated based on the estimated copy numbers and their respective read depths. We repeat this entire process for each possible value of differential read depth.

To provide additional information to assist in the scoring of purity and ploidy models, we segment the genome using a parametric segmentation method of our own design, described in detail later in the methods section. Models which under-estimate the read depth difference between integer copy numbers result in heavy oversegmentation (the introduction of too many breakpoints) of the genome, while models which over-estimate this difference will undersegment (the introduction of too few breakpoints) the genome. The model with the lowest read depth difference, corresponding to a tumor purity of 5% and a pentaploid tumor results in an unreasonably low estimate of read depth difference per copy number, and therefore provides a useful lower bound of segment contiguity. To measure undersegmentation, we use the median absolute deviation of read depth fitted to the mixture model, since this metric should be large in cases where read depth values are not represented by any components of the GMM. These metrics are factored into Equation 1, the scoring equation:

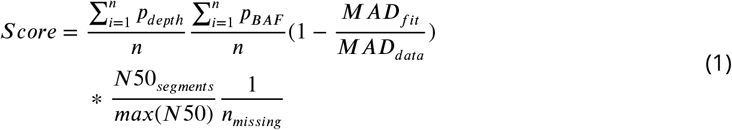

Where *p*_*depth*_ and *p*_*BAF*_ refer to the posterior probabilities for the read depth and the BAF GMMs, respectively, *N*50_*segments*_ refers to the segment N50 of the segmentation solution, defined as the size of the smallest segment for which all segments of that size and below account for half the length of the genome, *max*(*N*50) refers to the largest N50 value of all the models tested, and *n*_*m*_*issing* is the number of GMM components that explained less than 1% of the data, but whose two closest neighbor components did. *MAD*_*fit*_, described in Equation 2, refers to a weighted GMM median absolute deviation obtained by scaling the median absolute difference between the read depth values (*d*) and the *kth* GMM component means (*x*_*k*_) for each of the 11 components by the posteriors:

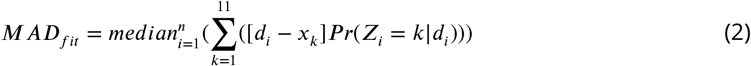

### Segmentation

Let *d* = *d*_1_, *d*_2_, …, *d*_*n*_ be the read depth values for the *n* genomic bins. Let *s* be a series of *k* = *n* − 1 similarity values describing the similarity of each bin to the *n* + 1st bin. We take a threshold *t* as the convergence threshold and iterate Algorithm 1 until convergence.

If the genome was pre-segmented, a vector of pre-existing segment lengths *z* exists. *f* is a mapping function that takes the depth values and *z* to return *v*, a vector of depth values which have been merged and averaged according to *z. g* takes the bins corresponding to the flagged transition values in *f lagged* to return *z*, a vector of the number of bins comprising each element in *m*. Eventually the algorithm converges on a set of segments whose length does not change, when all *s* exceed the specified threshold. This method is universal to any type of similarity metrics provided in *s*, on condition that a reasonable threshold is provided. For Ploidetect the threshold is determined through Monte Carlo simulation of the expected transition between two copy number values using the predicted means and the estimated variance of the data. This eliminates the need for parameter tuning by the user. The above segmentation method is known as segmentation by compression (SBC) as it iteratively “compresses” similar adjacent observations to achieve segmentation.

To compute the transition scores, let *d* = {*d*_0_, *d*_1_, *d*_2_, …, *d*_*n*_} be the read depths. Given an input vector *d* of read depth values, fit a GMM with *K* components. The posterior probability distribution for each *i, D*_*i*_ is given by equation 3:

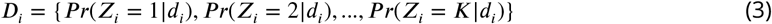

*D*_*i*_ describes, in other words, the likelihood of each *i*th point belonging to each GMM component. The transition score between an *i*th and *i* + 1st observation is therefore simply |*D*_*i*_ − *D*_*i*+1_|, which varies from 0, implying little evidence for transition, and 2, implying strong evidence for a transition. These transition scores are used for segmentation by Ploidetect.

#### Algorithm 1: Single iteration of segmentation by compression

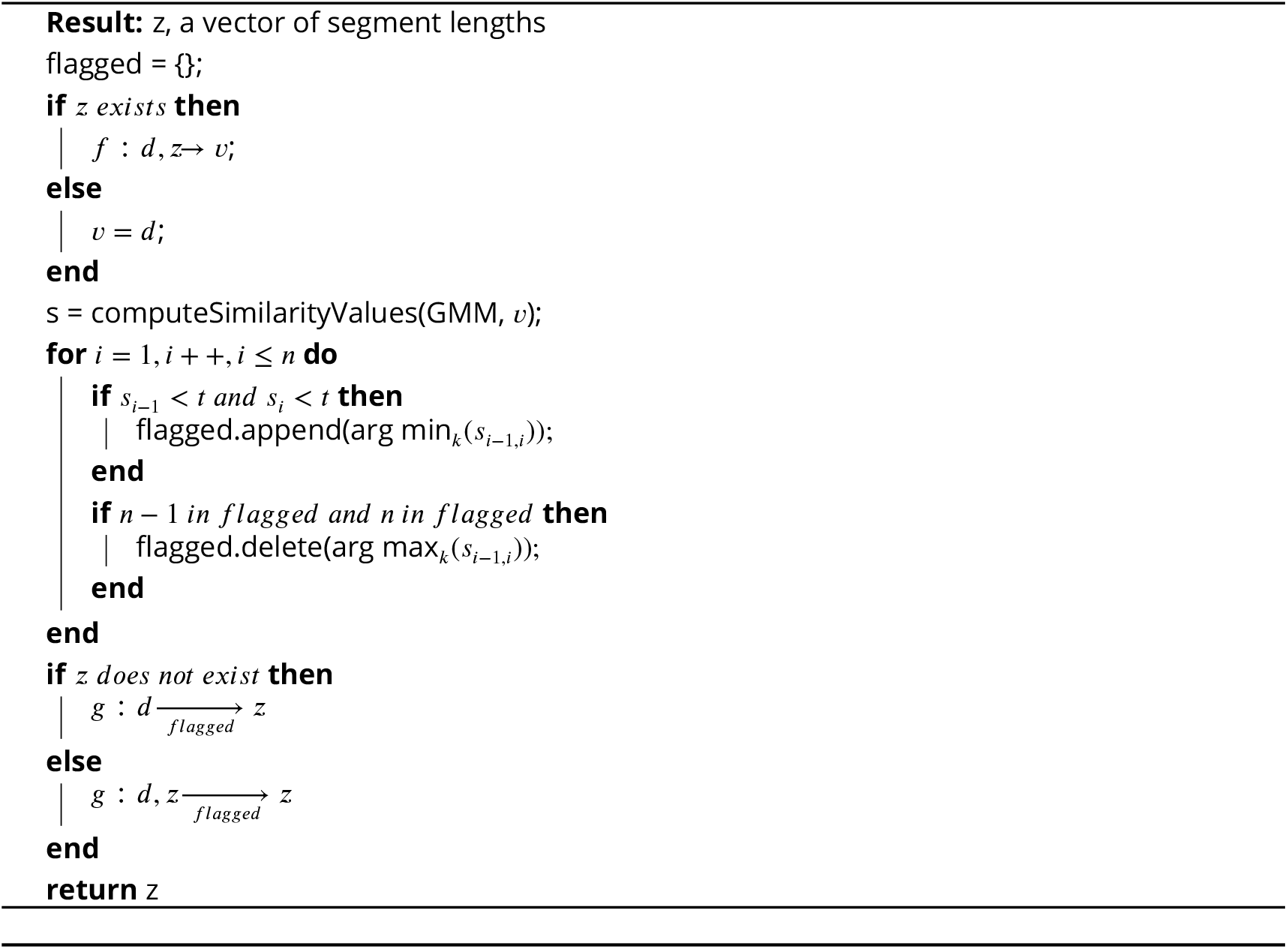

### CNV segmentation and detection

The overall strategy for segmentation is first to determine the posterior probability distribution of absolute copy number over the input data, and subsequently segmentation is performed to varying levels of stringency to segment the genome.

### Subclonal population detection

First a GMM corresponding to clonal copy number read depths is fit to the data. We perform SBC using the absolute differential GMM probabilities as the similarity values, and the 95^th^ percentile of the similarity values as the threshold. This threshold is purposefully low to enable segmentation of some of the copy number profile without risking undersegmentation of subclonal transitions, as they have not been explicitly defined in the GMM. To detect subclonal CNVs we fit another GMM which includes components for subclones with a clonal fraction of 0.5 or 0.25 between zero and ploidy + 3. We then use the resulting GMM in SBC to segment the genome with the subclone-aware transition values. Absolute copy number is assigned to the resulting segments, and we apply filters to retain high-quality subclone calls. We retain only those subclones which were the result of a transition from a clonal copy number segment, and the subclones whose copy number totals at least 5Mb of genomic content. Effectively, subclones mistakenly arising from noisy segmentation and those which are not well represented in the genome are excluded. This analysis identifies the catalogue of expected copy number states which occur within the whole genome, including subclonal CNVs with reasonable representation. These states are used to re-segment the genome using SBC to obtain a low-resolution copy number profile of the genome, which in turn is used to segment at high resolution in a coarse-to-fine process.

### Segmenting with resolution: a Coarse-to-fine approach

Thus far all of the genome segmentation has been performed on a set of meta-bins which are comprised of numerous smaller genomic bins in order to increase the signal-to-noise ratio. We now apply a coarse-to-fine approach to increase resolution while limiting the risk of oversegmentation from the noisier data. The subclone discovery step generates a map of CNV segments across the genome at the lower resolution provided. At this stage it can be assumed that the only CNVs which evaded detection were ones smaller than the size of the meta-bins. We divide the meta-bins in half each and map the previous segments to the data. We then fit a GMM to each of the segment-level median read depth and the meta-bin-level read depths and take the sums of the absolute differences in the posterior probability distributions for each bin as a metric describing how well each meta-bin belongs to its respective segment. For example, a meta-bin which is classified perfectly to a single GMM component but whose segment is classified perfectly to a different component would have a score of 2, and a meta-bin with no evidence of changing its classification would have a score of zero. To find a suitable threshold, we perform monte carlo simulation. We generate 10000 normally distributed read depths of invariant copy number, 10 times, for a range of copy numbers from zero to ploidy + 3. To estimate the expected variance for each copy number state, we use a linear model trained on the variance of the read depths for each copy number in the data. For each of the ten simulations, we obtain the second-highest transition score and use the median of the ten simulations as the differential posterior threshold for that copy number state. Segments whose transitions are flagged by this procedure are broken at those positions and SBC is then used on the broken segments to re-segment the candidate regions. We repeat this entire process, halving the bin size iteratively until either the genome’s segment N50 becomes halved since the previous iteration, indicating oversegmentation, or until the meta-bins are completely divided into their constituent bins.

One notable complication in estimating copy number likelihoods using GMMs is that occasionally local variation or extremely low abundance subclonal CNV may result in segments with an apparent fractional copy number slightly above or below their integer value. Particularly during Ploidetect’s coarse-to-fine segmentation, the data becomes progressively noisier and more prone to these offsets. These offsets artificially inflate the transition values used to identify segments to break, which result in a drastic increase in spurious breakpoint candidates. When computing GMMs during the coarse-to-fine segmentation stage, we slightly modify the GMM fitting on a per-segment basis. For each segment we record the median read depth from the previous iteration of the fit and re-center the single GMM component mean corresponding to the clonal CNV state. This ensures that when identifying bins which drastically shift from their parent segments, the expected transition scores are calculated from a correct starting position.

### Zygosity detection

The final step of the CNV calling procedure is the use of BAFs to estimate the zygosity of the CNVs. Based on the tumor purity we generate a GMM of BAF values for each observed copy number state in the genome, where each component corresponds to a specific allele state. For each segment, the maximum likelihood estimate of the allele state is used. For segments without BAF measurements, the segment is called as heterozygous. Plots depicting the CNV profile are automatically generated.

