## Supplementary Figure 1 for "Robust somatic copy number estimation using coarse-to-fine segmentation"

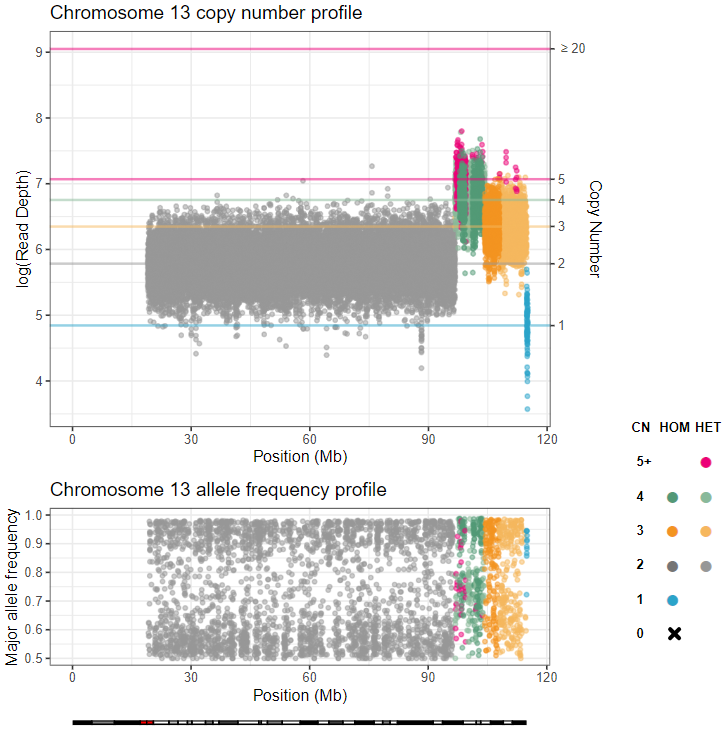

**Supplementary Figure 1.** Identification of CNVs in single-sample Oxford Nanopore data. Ploidetect was used to identify CNVs in a pediatric patient with anophthalmia. CNVs were identified by Ploidetect on Chromosome 13q. Detailed case report described in Boerkoel *et al.*, 2022.
