## Supplementary Figure 2 for "Robust somatic copy number estimation using coarse-to-fine segmentation"

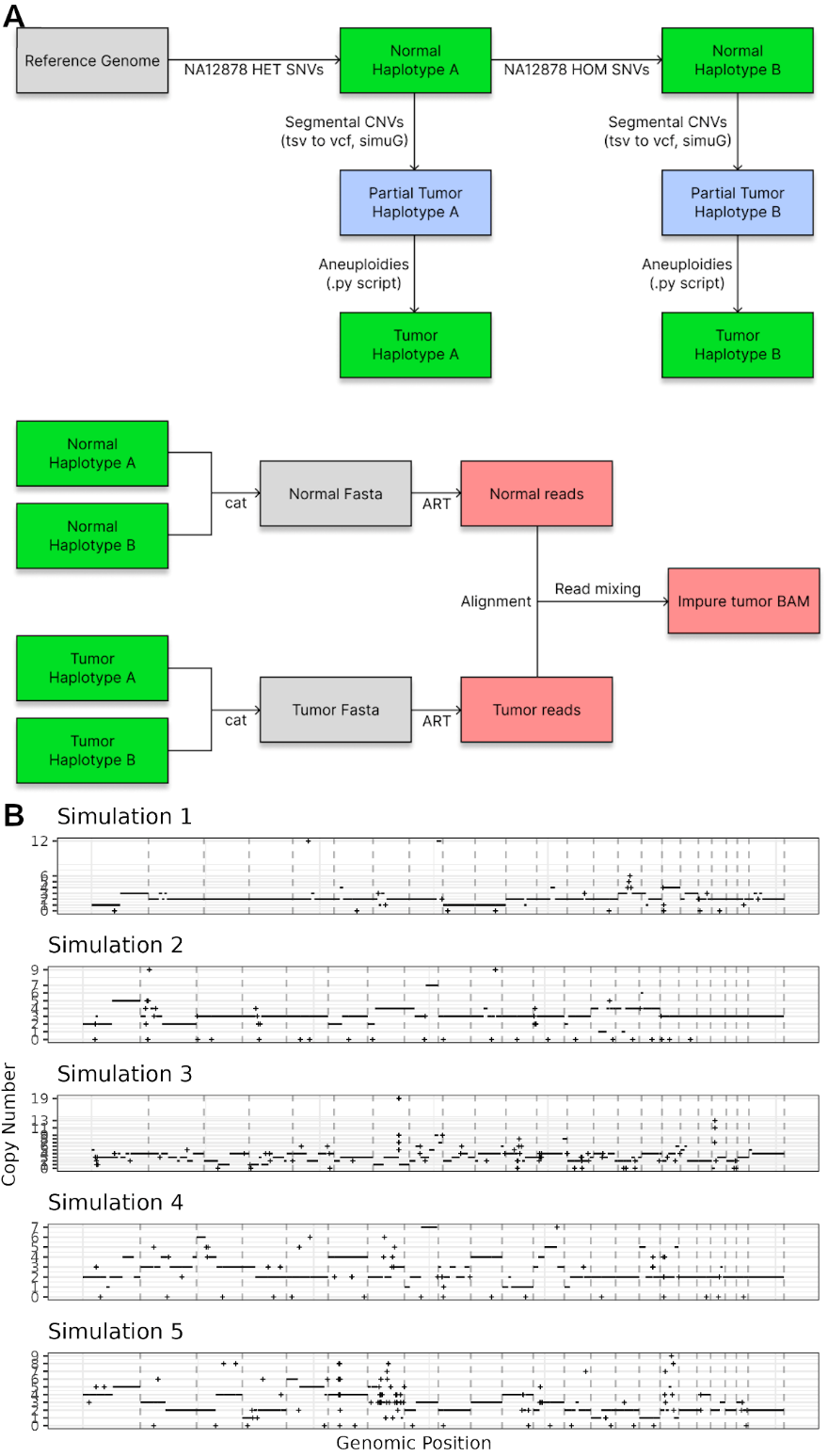

**Supplementary Figure 2.** Simulation of somatic CNVs. **A.** Workflow used for generation of a synthetic CNV profile. Normal haplotypes are generated by introducing variants in random and semi-random positions across the genome. Each “germline” haplotype is subjected to CNV using simuG as well as a custom python script to introduce aneuploidies. Resulting genomes are used with ART, aligned, and mixed at known proportions to generate synthetic impure sequence data. **B.** Synthetic CNV profiles used in this study. Points marked as “X” indicate a segment smaller than 10Mbp in size.
