## Supplementary Figure 3 for "Robust somatic copy number estimation using coarse-to-fine segmentation"

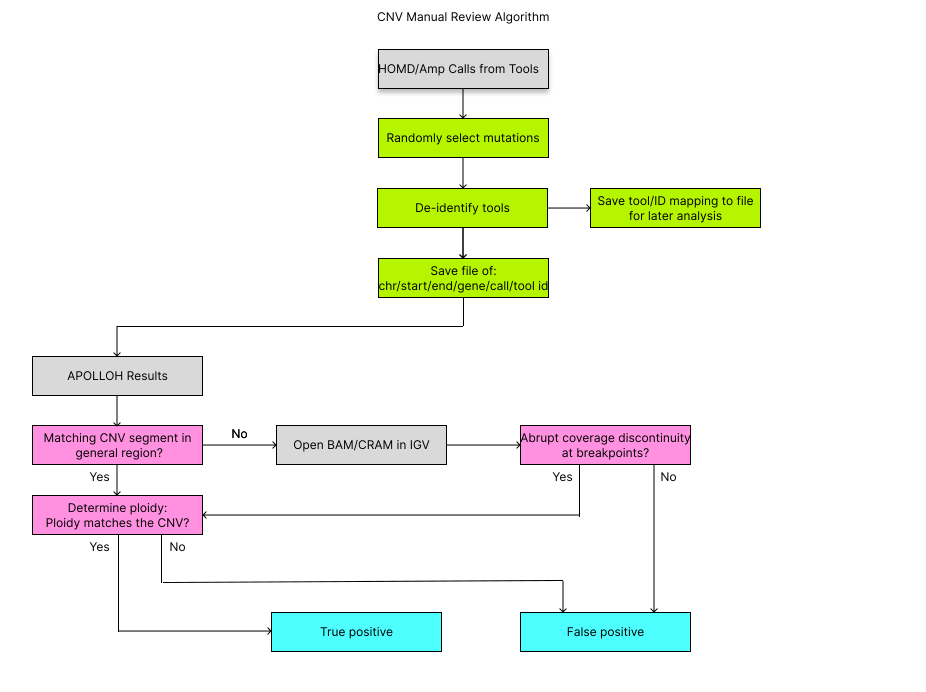

**Supplementary Figure 3.** Workflow for manual review of called CNVs. De-identified CNV calls from each CNV tool were saved to a text file without human observation of the intermediates. Resulting calls were manually reviewed first by using existing POG data as an unbiased first-pass. CNVs which were easily confirmed using this approach were immediately marked as true positives. Otherwise, integrated genome viewer was used to investigate sequence read alignment in the region of interest, and if a CNV was observed, it was marked as a true positive. In all other cases, including in cases where a CNV was identified only because of an error in ploidy estimation, it was marked as a false positive.
