## Supplementary Figure 4 for "Robust somatic copy number estimation using coarse-to-fine segmentation"

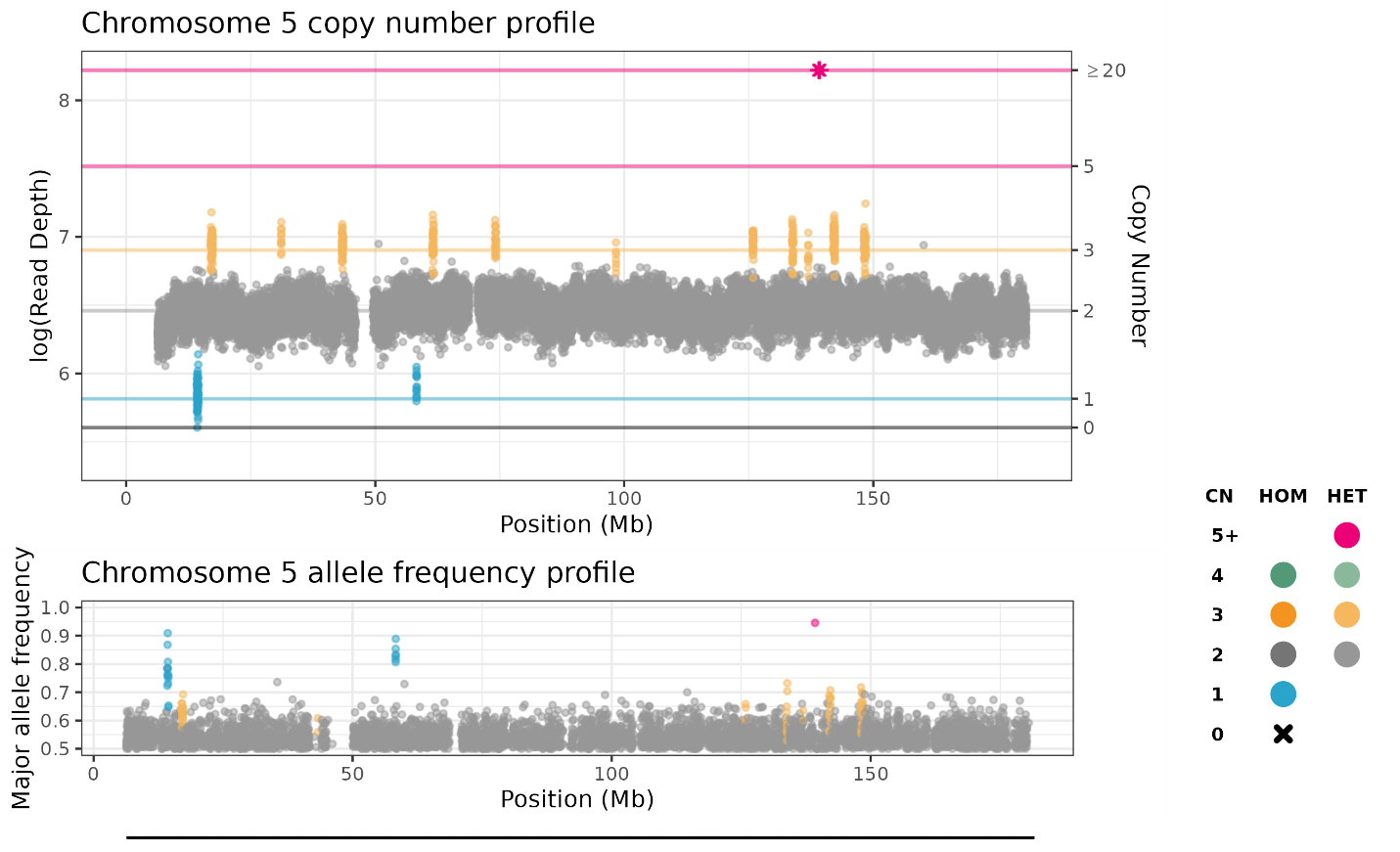
**Supplementary Figure 4.** PGA may not account for all patterns of CIN. Ploidetect CNV calls on a representative chromosome, Chromosome 5, from an example POG tumor is shown. Interspersed, focal CNVs result in a computed PGA of 7%, while the N50 was reduced from maximum by 87%.
