## Supplementary Figure 5 for "Robust somatic copy number estimation using coarse-to-fine segmentation"

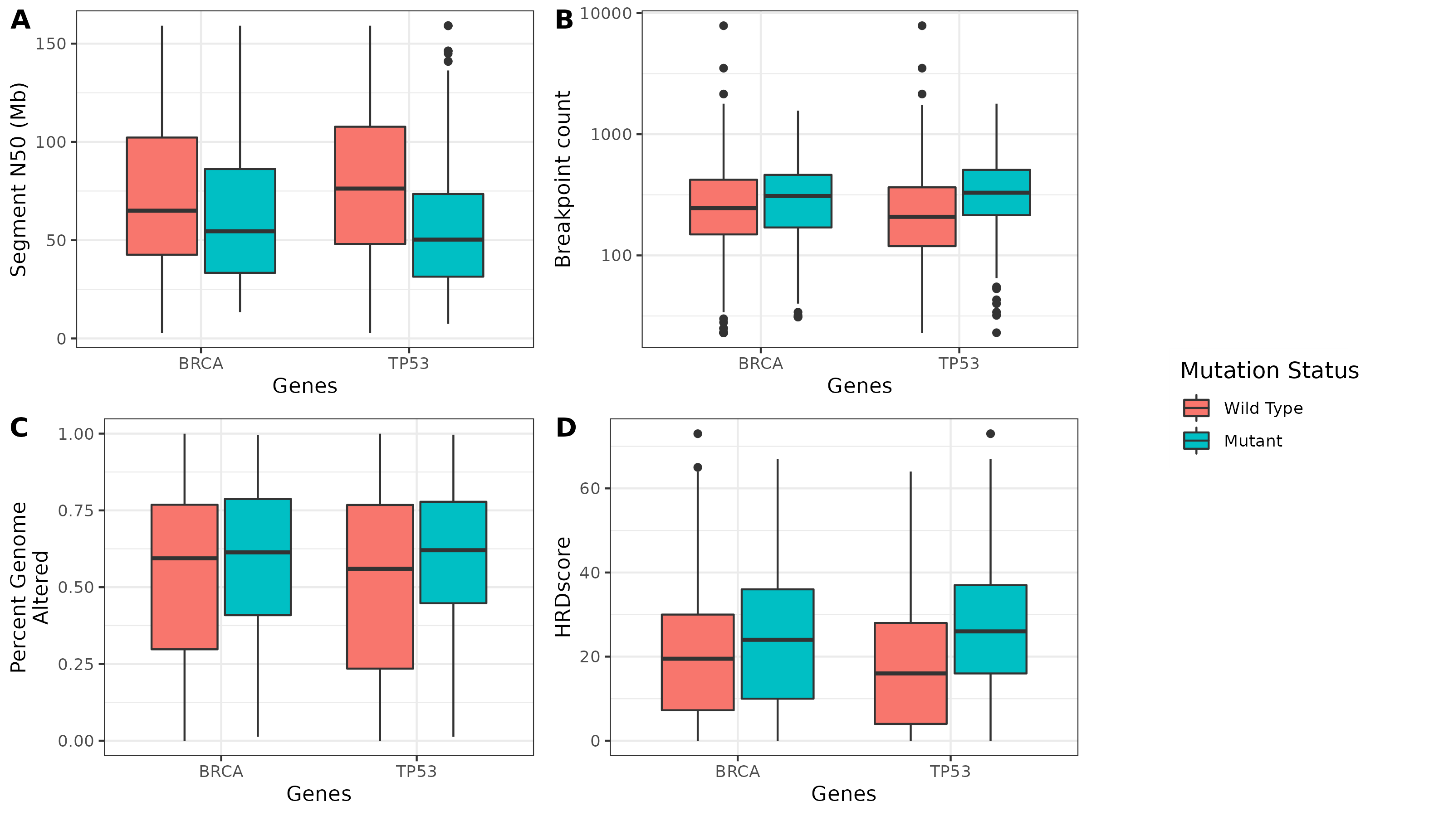
**Supplementary Figure 5.** Alteration of various CIN metrics according to mutation status. **A.** Difference in segment N50 in TP53 and BRCA wild-type versus mutant tumors. **B.**  Difference in segment count in TP53 and BRCA wild-type versus mutant tumors. **C.**  Difference in PGA in TP53 and BRCA wild-type versus mutant tumors. **D.**  Difference in HRDscore in TP53 and BRCA wild-type versus mutant tumors.
