## Supplementary Figure 6 for "Robust somatic copy number estimation using coarse-to-fine segmentation"

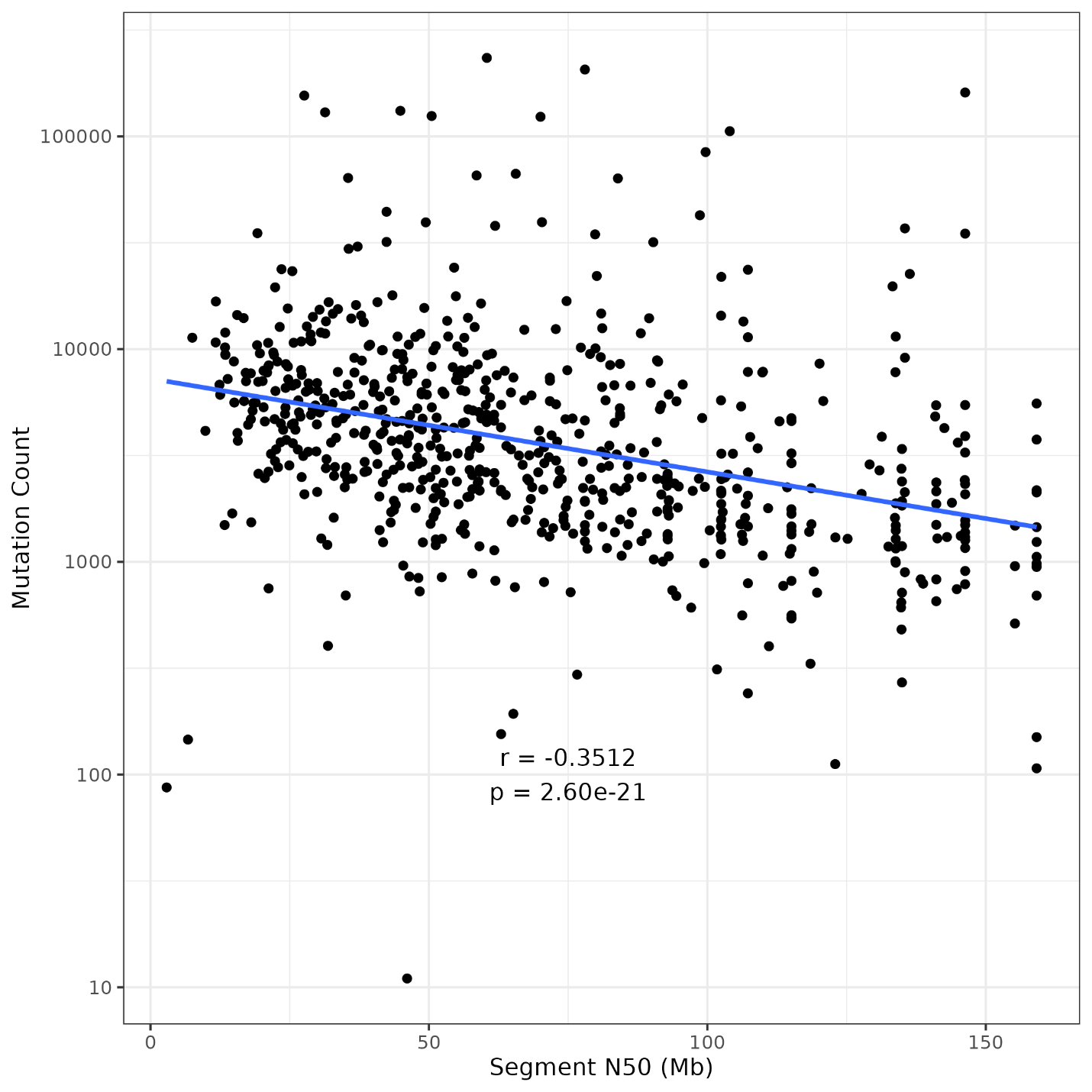

**Supplementary Figure 6.** Tumor mutational burden is associated with elevated CIN measured using the segment N50. Line of best fit, Pearson’s rho and linear model p-value are shown for the segment N50 versus log10 mutation count values.
