## Supplementary Table 1 for "Robust somatic copy number estimation using coarse-to-fine segmentation"

| **seqnames** | **start** | **end** | **width** | **CN** | **sim** |
| --- | --- | --- | --- | --- | --- |
| 1 | 1 | 100000000 | 1E+08 | 1 | SIM1 |
| 1 | 1E+08 | 100010000 | 10000 | 0 | SIM1 |
| 1 | 1E+08 | 125250000 | 25240000 | 1 | SIM1 |
| 1 | 1.25E+08 | 249250620 | 1.24E+08 | 3 | SIM1 |
| 2 | 1 | 45000000 | 45000000 | 2 | SIM1 |
| 2 | 45000001 | 60000000 | 15000000 | 3 | SIM1 |
| 2 | 60000001 | 70000000 | 10000000 | 2 | SIM1 |
| 2 | 70000001 | 80000000 | 10000000 | 3 | SIM1 |
| 2 | 80000001 | 90000000 | 10000000 | 2 | SIM1 |
| 2 | 90000001 | 243199373 | 1.53E+08 | 2 | SIM1 |
| 3 | 1 | 198022430 | 1.98E+08 | 2 | SIM1 |
| 4 | 1 | 191154276 | 1.91E+08 | 2 | SIM1 |
| 5 | 1 | 68000000 | 68000000 | 2 | SIM1 |
| 5 | 68000001 | 68100000 | 100000 | 12 | SIM1 |
| 5 | 68100001 | 80000000 | 11900000 | 2 | SIM1 |
| 5 | 80000001 | 95000000 | 15000000 | 3 | SIM1 |
| 5 | 95000001 | 180915259 | 85915259 | 2 | SIM1 |
| 6 | 1 | 25000000 | 25000000 | 2 | SIM1 |
| 6 | 25000001 | 40000000 | 15000000 | 4 | SIM1 |
| 6 | 40000001 | 80000000 | 40000000 | 2 | SIM1 |
| 6 | 80000001 | 90000000 | 10000000 | 2 | SIM1 |
| 6 | 90000001 | 100000000 | 10000000 | 2 | SIM1 |
| 6 | 1E+08 | 100010000 | 10000 | 0 | SIM1 |
| 6 | 1E+08 | 171115067 | 71105067 | 2 | SIM1 |
| 7 | 1 | 20000000 | 20000000 | 2 | SIM1 |
| 7 | 20000001 | 25000000 | 5000000 | 3 | SIM1 |
| 7 | 25000001 | 35000000 | 10000000 | 1 | SIM1 |
| 7 | 35000001 | 50000000 | 15000000 | 3 | SIM1 |
| 7 | 50000001 | 59000000 | 9000000 | 1 | SIM1 |
| 7 | 59000001 | 159138663 | 1E+08 | 2 | SIM1 |
| 8 | 1 | 120000000 | 1.2E+08 | 2 | SIM1 |
| 8 | 1.2E+08 | 140000000 | 20000000 | 12 | SIM1 |
| 8 | 1.4E+08 | 146364022 | 6364022 | 2 | SIM1 |
| 9 | 1 | 21950000 | 21950000 | 1 | SIM1 |
| 9 | 21950001 | 22000000 | 50000 | 0 | SIM1 |
| 9 | 22000001 | 141213431 | 1.19E+08 | 1 | SIM1 |
| 10 | 1 | 89600000 | 89600000 | 1 | SIM1 |
| 10 | 89600001 | 89750000 | 150000 | 0 | SIM1 |
| 10 | 89750001 | 135534747 | 45784747 | 1 | SIM1 |
| 11 | 1 | 80000000 | 80000000 | 2 | SIM1 |
| 11 | 80000001 | 90000000 | 10000000 | 1 | SIM1 |
| 11 | 90000001 | 135006516 | 45006516 | 2 | SIM1 |
| 12 | 1 | 60000000 | 60000000 | 2 | SIM1 |
| 12 | 60000001 | 80000000 | 20000000 | 4 | SIM1 |
| 12 | 80000001 | 133851895 | 53851895 | 2 | SIM1 |
| 13 | 1 | 73000000 | 73000000 | 2 | SIM1 |
| 13 | 73000001 | 79000000 | 6000000 | 3 | SIM1 |
| 13 | 79000001 | 115169878 | 36169878 | 2 | SIM1 |
| 14 | 1 | 67500000 | 67500000 | 2 | SIM1 |
| 14 | 67500001 | 67505000 | 5000 | 0 | SIM1 |
| 14 | 67505001 | 107349540 | 39844540 | 2 | SIM1 |
| 15 | 1 | 40000000 | 40000000 | 3 | SIM1 |
| 15 | 40000001 | 45000000 | 5000000 | 4 | SIM1 |
| 15 | 45000001 | 50000000 | 5000000 | 5 | SIM1 |
| 15 | 50000001 | 55000000 | 5000000 | 6 | SIM1 |
| 15 | 55000001 | 60000000 | 5000000 | 4 | SIM1 |
| 15 | 60000001 | 102531392 | 42531392 | 3 | SIM1 |
| 16 | 1 | 20000000 | 20000000 | 3 | SIM1 |
| 16 | 20000001 | 30000000 | 10000000 | 1 | SIM1 |
| 16 | 30000001 | 90354753 | 60354753 | 2 | SIM1 |
| 17 | 1 | 7500000 | 7500000 | 4 | SIM1 |
| 17 | 7500001 | 7550000 | 50000 | 1 | SIM1 |
| 17 | 7550001 | 7600000 | 50000 | 0 | SIM1 |
| 17 | 7600001 | 81195210 | 73595210 | 4 | SIM1 |
| 18 | 1 | 18000000 | 18000000 | 2 | SIM1 |
| 18 | 18000001 | 78000000 | 60000000 | 3 | SIM1 |
| 18 | 78000001 | 78077248 | 77248 | 2 | SIM1 |
| 19 | 1 | 7267000 | 7267000 | 2 | SIM1 |
| 19 | 7267001 | 7268000 | 1000 | 0 | SIM1 |
| 19 | 7268001 | 35000000 | 27732000 | 2 | SIM1 |
| 19 | 35000001 | 40000000 | 5000000 | 3 | SIM1 |
| 19 | 40000001 | 59128983 | 19128983 | 2 | SIM1 |
| 20 | 1 | 33865000 | 33865000 | 2 | SIM1 |
| 20 | 33865001 | 33875000 | 10000 | 0 | SIM1 |
| 20 | 33875001 | 63025520 | 29150520 | 2 | SIM1 |
| 21 | 1 | 48129895 | 48129895 | 2 | SIM1 |
| 22 | 1 | 26000000 | 26000000 | 2 | SIM1 |
| 22 | 26000001 | 31000000 | 5000000 | 1 | SIM1 |
| 22 | 31000001 | 51304566 | 20304566 | 2 | SIM1 |
| X | 1 | 20000000 | 20000000 | 2 | SIM1 |
| X | 20000001 | 36000000 | 16000000 | 3 | SIM1 |
| X | 36000001 | 45000000 | 9000000 | 2 | SIM1 |
| X | 45000001 | 58000000 | 13000000 | 3 | SIM1 |
| X | 58000001 | 155270560 | 97270560 | 2 | SIM1 |
| 1 | 1 | 52825341 | 52825341 | 2 | SIM2 |
| 1 | 52825342 | 52875341 | 50000 | 0 | SIM2 |
| 1 | 52875342 | 55324498 | 2449157 | 2 | SIM2 |
| 1 | 55324499 | 65324498 | 10000000 | 3 | SIM2 |
| 1 | 65324499 | 126325154 | 61000656 | 2 | SIM2 |
| 1 | 1.26E+08 | 249250620 | 1.23E+08 | 5 | SIM2 |
| 2 | 1 | 21597536 | 21597536 | 3 | SIM2 |
| 2 | 21597537 | 22604413 | 1006877 | 4 | SIM2 |
| 2 | 22604414 | 25604413 | 3000000 | 2 | SIM2 |
| 2 | 25604414 | 31953968 | 6349555 | 5 | SIM2 |
| 2 | 31953969 | 32004215 | 50247 | 0 | SIM2 |
| 2 | 32004216 | 35741965 | 3737750 | 5 | SIM2 |
| 2 | 35741966 | 42425819 | 6683854 | 9 | SIM2 |
| 2 | 42425820 | 56695481 | 14269662 | 4 | SIM2 |
| 2 | 56695482 | 62542196 | 5846715 | 3 | SIM2 |
| 2 | 62542197 | 69420126 | 6877930 | 4 | SIM2 |
| 2 | 69420127 | 94198174 | 24778048 | 3 | SIM2 |
| 2 | 94198175 | 243199372 | 1.49E+08 | 2 | SIM2 |
| 3 | 1 | 5123460 | 5123460 | 3 | SIM2 |
| 3 | 5123461 | 5159684 | 36224 | 0 | SIM2 |
| 3 | 5159685 | 65248148 | 60088464 | 3 | SIM2 |
| 3 | 65248149 | 65272615 | 24467 | 0 | SIM2 |
| 3 | 65272616 | 198022430 | 1.33E+08 | 3 | SIM2 |
| 4 | 1 | 55465165 | 55465165 | 3 | SIM2 |
| 4 | 55465166 | 62214563 | 6749398 | 4 | SIM2 |
| 4 | 62214564 | 67187568 | 4973005 | 3 | SIM2 |
| 4 | 67187569 | 72841584 | 5654016 | 2 | SIM2 |
| 4 | 72841585 | 72888456 | 46872 | 0 | SIM2 |
| 4 | 72888457 | 82147558 | 9259102 | 2 | SIM2 |
| 4 | 82147559 | 191154276 | 1.09E+08 | 3 | SIM2 |
| 5 | 1 | 60086907 | 60086907 | 3 | SIM2 |
| 5 | 60086908 | 60111465 | 24558 | 0 | SIM2 |
| 5 | 60111466 | 101061202 | 40949737 | 3 | SIM2 |
| 5 | 1.01E+08 | 101114165 | 52963 | 0 | SIM2 |
| 5 | 1.01E+08 | 180915260 | 79801095 | 3 | SIM2 |
| 6 | 1 | 59890273 | 59890272 | 2 | SIM2 |
| 6 | 59890274 | 89346510 | 29456237 | 3 | SIM2 |
| 6 | 89346511 | 89383512 | 37002 | 0 | SIM2 |
| 6 | 89383513 | 171115067 | 81731555 | 3 | SIM2 |
| 7 | 1 | 21125684 | 21125683 | 2 | SIM2 |
| 7 | 21125685 | 21161542 | 35858 | 0 | SIM2 |
| 7 | 21161543 | 31506935 | 10345393 | 2 | SIM2 |
| 7 | 31506936 | 159138663 | 1.28E+08 | 4 | SIM2 |
| 8 | 1 | 43624697 | 43624696 | 4 | SIM2 |
| 8 | 43624698 | 86511780 | 42887083 | 3 | SIM2 |
| 8 | 86511781 | 86534618 | 22838 | 0 | SIM2 |
| 8 | 86534619 | 92514846 | 5980228 | 3 | SIM2 |
| 8 | 92514847 | 146364021 | 53849175 | 7 | SIM2 |
| 9 | 1 | 21947751 | 21947751 | 3 | SIM2 |
| 9 | 21947752 | 21995323 | 47572 | 0 | SIM2 |
| 9 | 21995324 | 141213431 | 1.19E+08 | 3 | SIM2 |
| 10 | 1 | 55468513 | 55468513 | 3 | SIM2 |
| 10 | 55468514 | 72145247 | 16676734 | 4 | SIM2 |
| 10 | 72145248 | 89623383 | 17478136 | 3 | SIM2 |
| 10 | 89623384 | 89731687 | 108304 | 0 | SIM2 |
| 10 | 89731688 | 103154278 | 13422591 | 3 | SIM2 |
| 10 | 1.03E+08 | 109145274 | 5990996 | 9 | SIM2 |
| 10 | 1.09E+08 | 135534747 | 26389473 | 3 | SIM2 |
| 11 | 1 | 64795481 | 64795481 | 3 | SIM2 |
| 11 | 64795482 | 64831468 | 35987 | 0 | SIM2 |
| 11 | 64831469 | 135006516 | 70175048 | 3 | SIM2 |
| 12 | 1 | 5265481 | 5265481 | 3 | SIM2 |
| 12 | 5265482 | 7044838 | 1779357 | 2 | SIM2 |
| 12 | 7044839 | 8341568 | 1296730 | 4 | SIM2 |
| 12 | 8341569 | 14872433 | 6530865 | 2 | SIM2 |
| 12 | 14872434 | 63794910 | 48922477 | 3 | SIM2 |
| 12 | 63794911 | 63845416 | 50506 | 0 | SIM2 |
| 12 | 63845417 | 133851895 | 70006479 | 3 | SIM2 |
| 13 | 1 | 16980174 | 16980173 | 2 | SIM2 |
| 13 | 16980175 | 48877887 | 31897713 | 3 | SIM2 |
| 13 | 48877888 | 49056026 | 178139 | 0 | SIM2 |
| 13 | 49056027 | 115169878 | 66113852 | 3 | SIM2 |
| 14 | 1 | 31368373 | 31368373 | 4 | SIM2 |
| 14 | 31368374 | 67229004 | 35860631 | 1 | SIM2 |
| 14 | 67229005 | 79389090 | 12160086 | 4 | SIM2 |
| 14 | 79389091 | 83416633 | 4027543 | 5 | SIM2 |
| 14 | 83416634 | 95421813 | 12005180 | 4 | SIM2 |
| 14 | 95421814 | 95464153 | 42340 | 0 | SIM2 |
| 14 | 95464154 | 107349540 | 11885387 | 4 | SIM2 |
| 15 | 1 | 25025610 | 25025610 | 4 | SIM2 |
| 15 | 25025611 | 42175501 | 17149891 | 1 | SIM2 |
| 15 | 42175502 | 74287057 | 32111556 | 4 | SIM2 |
| 15 | 74287058 | 74340168 | 53111 | 0 | SIM2 |
| 15 | 74340169 | 102531392 | 28191224 | 4 | SIM2 |
| 16 | 1 | 6344927 | 6344927 | 4 | SIM2 |
| 16 | 6344928 | 16867964 | 10523037 | 6 | SIM2 |
| 16 | 16867965 | 55456128 | 38588164 | 4 | SIM2 |
| 16 | 55456129 | 55495486 | 39358 | 0 | SIM2 |
| 16 | 55495487 | 90354753 | 34859267 | 4 | SIM2 |
| 17 | 1 | 7541739 | 7541739 | 3 | SIM2 |
| 17 | 7541740 | 7590808 | 49069 | 0 | SIM2 |
| 17 | 7590809 | 41196312 | 33605504 | 3 | SIM2 |
| 17 | 41196313 | 41277381 | 81069 | 0 | SIM2 |
| 17 | 41277382 | 81195210 | 39917829 | 3 | SIM2 |
| 18 | 1 | 50164852 | 50164852 | 3 | SIM2 |
| 18 | 50164853 | 50204851 | 39999 | 0 | SIM2 |
| 18 | 50204852 | 78077248 | 27872397 | 3 | SIM2 |
| 19 | 1 | 59128983 | 59128983 | 3 | SIM2 |
| 20 | 1 | 63025520 | 63025520 | 3 | SIM2 |
| 21 | 1 | 48129895 | 48129895 | 3 | SIM2 |
| 22 | 1 | 51304566 | 51304566 | 3 | SIM2 |
| X | 1 | 155270560 | 1.55E+08 | 3 | SIM2 |
| 1 | 1 | 12614889 | 12614888 | 5 | SIM3 |
| 1 | 12614890 | 20416887 | 7801998 | 3 | SIM3 |
| 1 | 20416888 | 21068714 | 651827 | 2 | SIM3 |
| 1 | 21068715 | 22168714 | 1100000 | 1 | SIM3 |
| 1 | 22168715 | 26976814 | 4808100 | 3 | SIM3 |
| 1 | 26976815 | 27243562 | 266748 | 1 | SIM3 |
| 1 | 27243563 | 99614857 | 72371295 | 3 | SIM3 |
| 1 | 99614858 | 100481067 | 866210 | 4 | SIM3 |
| 1 | 1E+08 | 114691487 | 14210420 | 3 | SIM3 |
| 1 | 1.15E+08 | 126156043 | 11464556 | 4 | SIM3 |
| 1 | 1.26E+08 | 148791867 | 22635824 | 3 | SIM3 |
| 1 | 1.49E+08 | 165487197 | 16695330 | 4 | SIM3 |
| 1 | 1.65E+08 | 170941879 | 5454682 | 2 | SIM3 |
| 1 | 1.71E+08 | 249250621 | 78308742 | 4 | SIM3 |
| 2 | 1 | 45698179 | 45698178 | 4 | SIM3 |
| 2 | 45698180 | 55147845 | 9449666 | 3 | SIM3 |
| 2 | 55147846 | 85681487 | 30533642 | 4 | SIM3 |
| 2 | 85681488 | 124718987 | 39037500 | 3 | SIM3 |
| 2 | 1.25E+08 | 137178998 | 12460011 | 2 | SIM3 |
| 2 | 1.37E+08 | 168871467 | 31692469 | 3 | SIM3 |
| 2 | 1.69E+08 | 169016084 | 144617 | 5 | SIM3 |
| 2 | 1.69E+08 | 194618778 | 25602694 | 2 | SIM3 |
| 2 | 1.95E+08 | 243199373 | 48580595 | 4 | SIM3 |
| 3 | 1 | 60876874 | 60876873 | 2 | SIM3 |
| 3 | 60876875 | 60914861 | 37987 | 0 | SIM3 |
| 3 | 60914862 | 112781649 | 51866788 | 1 | SIM3 |
| 3 | 1.13E+08 | 142781649 | 30000000 | 4 | SIM3 |
| 3 | 1.43E+08 | 175614877 | 32833228 | 4 | SIM3 |
| 3 | 1.76E+08 | 176136487 | 521610 | 4 | SIM3 |
| 3 | 1.76E+08 | 198022430 | 21885943 | 4 | SIM3 |
| 4 | 1 | 8146387 | 8146386 | 1 | SIM3 |
| 4 | 8146388 | 8148418 | 2031 | 0 | SIM3 |
| 4 | 8148419 | 42194807 | 34046389 | 1 | SIM3 |
| 4 | 42194808 | 53186478 | 10991671 | 3 | SIM3 |
| 4 | 53186479 | 72617058 | 19430580 | 1 | SIM3 |
| 4 | 72617059 | 72659787 | 42729 | 4 | SIM3 |
| 4 | 72659788 | 85168477 | 12508690 | 1 | SIM3 |
| 4 | 85168478 | 92178691 | 7010214 | 4 | SIM3 |
| 4 | 92178692 | 92196145 | 17454 | 2 | SIM3 |
| 4 | 92196146 | 103198677 | 11002532 | 4 | SIM3 |
| 4 | 1.03E+08 | 153198677 | 50000000 | 3 | SIM3 |
| 4 | 1.53E+08 | 164618782 | 11420105 | 2 | SIM3 |
| 4 | 1.65E+08 | 189507748 | 24888966 | 4 | SIM3 |
| 4 | 1.9E+08 | 191154276 | 1646528 | 2 | SIM3 |
| 5 | 1 | 37465878 | 37465878 | 4 | SIM3 |
| 5 | 37465879 | 37481481 | 15603 | 6 | SIM3 |
| 5 | 37481482 | 61614587 | 24133106 | 4 | SIM3 |
| 5 | 61614588 | 110867982 | 49253395 | 2 | SIM3 |
| 5 | 1.11E+08 | 146961872 | 36093890 | 4 | SIM3 |
| 5 | 1.47E+08 | 156916788 | 9954916 | 6 | SIM3 |
| 5 | 1.57E+08 | 163148678 | 6231890 | 4 | SIM3 |
| 5 | 1.63E+08 | 163199678 | 51000 | 0 | SIM3 |
| 5 | 1.63E+08 | 180915260 | 17715582 | 4 | SIM3 |
| 6 | 1 | 25618787 | 25618786 | 2 | SIM3 |
| 6 | 25618788 | 81681787 | 56063000 | 3 | SIM3 |
| 6 | 81681788 | 81695867 | 14080 | 2 | SIM3 |
| 6 | 81695868 | 171115067 | 89419200 | 3 | SIM3 |
| 7 | 1 | 45615664 | 45615663 | 1 | SIM3 |
| 7 | 45615665 | 112654894 | 67039230 | 3 | SIM3 |
| 7 | 1.13E+08 | 112854894 | 200000 | 5 | SIM3 |
| 7 | 1.13E+08 | 112966418 | 111524 | 7 | SIM3 |
| 7 | 1.13E+08 | 113019687 | 53269 | 9 | SIM3 |
| 7 | 1.13E+08 | 113298674 | 278987 | 19 | SIM3 |
| 7 | 1.13E+08 | 159138663 | 45839989 | 1 | SIM3 |
| 8 | 1 | 26986147 | 26986146 | 2 | SIM3 |
| 8 | 26986148 | 44618789 | 17632642 | 6 | SIM3 |
| 8 | 44618790 | 46965841 | 2347052 | 4 | SIM3 |
| 8 | 46965842 | 48617854 | 1652013 | 6 | SIM3 |
| 8 | 48617855 | 68816404 | 20198550 | 3 | SIM3 |
| 8 | 68816405 | 68968477 | 152073 | 2 | SIM3 |
| 8 | 68968478 | 86864187 | 17895710 | 3 | SIM3 |
| 8 | 86864188 | 109684987 | 22820800 | 5 | SIM3 |
| 8 | 1.1E+08 | 126685417 | 17000430 | 9 | SIM3 |
| 8 | 1.27E+08 | 143364022 | 16678605 | 7 | SIM3 |
| 8 | 1.43E+08 | 146364022 | 3000000 | 9 | SIM3 |
| 9 | 1 | 38816478 | 38816477 | 2 | SIM3 |
| 9 | 38816479 | 74687978 | 35871500 | 3 | SIM3 |
| 9 | 74687979 | 82687897 | 7999919 | 4 | SIM3 |
| 9 | 82687898 | 82728791 | 40894 | 6 | SIM3 |
| 9 | 82728792 | 110698174 | 27969383 | 4 | SIM3 |
| 9 | 1.11E+08 | 141213431 | 30515257 | 3 | SIM3 |
| 10 | 1 | 13681489 | 13681488 | 2 | SIM3 |
| 10 | 13681490 | 50618779 | 36937290 | 4 | SIM3 |
| 10 | 50618780 | 50635846 | 17067 | 2 | SIM3 |
| 10 | 50635847 | 112984795 | 62348949 | 4 | SIM3 |
| 10 | 1.13E+08 | 126641830 | 13657035 | 6 | SIM3 |
| 10 | 1.27E+08 | 135534747 | 8892917 | 4 | SIM3 |
| 11 | 1 | 37190877 | 37190876 | 4 | SIM3 |
| 11 | 37190878 | 37278914 | 88037 | 5 | SIM3 |
| 11 | 37278915 | 50261884 | 12982970 | 4 | SIM3 |
| 11 | 50261885 | 50618779 | 356895 | 2 | SIM3 |
| 11 | 50618780 | 50635846 | 17067 | 0 | SIM3 |
| 11 | 50635847 | 54925747 | 4289901 | 2 | SIM3 |
| 11 | 54925748 | 57961879 | 3036132 | 6 | SIM3 |
| 11 | 57961880 | 58596871 | 634992 | 8 | SIM3 |
| 11 | 58596872 | 64175864 | 5578993 | 4 | SIM3 |
| 11 | 64175865 | 71618475 | 7442611 | 6 | SIM3 |
| 11 | 71618476 | 83618047 | 11999572 | 2 | SIM3 |
| 11 | 83618048 | 135006516 | 51388469 | 3 | SIM3 |
| 12 | 1 | 10368741 | 10368740 | 3 | SIM3 |
| 12 | 10368742 | 10394152 | 25411 | 2 | SIM3 |
| 12 | 10394153 | 12681741 | 2287589 | 3 | SIM3 |
| 12 | 12681742 | 12751416 | 69675 | 4 | SIM3 |
| 12 | 12751417 | 20487898 | 7736482 | 3 | SIM3 |
| 12 | 20487899 | 20512356 | 24458 | 4 | SIM3 |
| 12 | 20512357 | 20616145 | 103789 | 3 | SIM3 |
| 12 | 20616146 | 112684178 | 92068033 | 4 | SIM3 |
| 12 | 1.13E+08 | 132851895 | 20167717 | 8 | SIM3 |
| 12 | 1.33E+08 | 133851895 | 1000000 | 4 | SIM3 |
| 13 | 1 | 31941885 | 31941884 | 2 | SIM3 |
| 13 | 31941886 | 32090746 | 148861 | 0 | SIM3 |
| 13 | 32090747 | 57489167 | 25398421 | 2 | SIM3 |
| 13 | 57489168 | 63416879 | 5927712 | 3 | SIM3 |
| 13 | 63416880 | 63458647 | 41768 | 2 | SIM3 |
| 13 | 63458648 | 65416887 | 1958240 | 3 | SIM3 |
| 13 | 65416888 | 65444681 | 27794 | 0 | SIM3 |
| 13 | 65444682 | 90618487 | 25173806 | 2 | SIM3 |
| 13 | 90618488 | 90861384 | 242897 | 4 | SIM3 |
| 13 | 90861385 | 98418143 | 7556759 | 2 | SIM3 |
| 13 | 98418144 | 115169878 | 16751735 | 3 | SIM3 |
| 14 | 1 | 35419887 | 35419886 | 4 | SIM3 |
| 14 | 35419888 | 40968741 | 5548854 | 3 | SIM3 |
| 14 | 40968742 | 41086179 | 117438 | 4 | SIM3 |
| 14 | 41086180 | 62615438 | 21529259 | 3 | SIM3 |
| 14 | 62615439 | 92417858 | 29802420 | 4 | SIM3 |
| 14 | 92417859 | 107349540 | 14931682 | 2 | SIM3 |
| 15 | 1 | 24689778 | 24689777 | 2 | SIM3 |
| 15 | 24689779 | 24711878 | 22100 | 0 | SIM3 |
| 15 | 24711879 | 34986142 | 10274264 | 2 | SIM3 |
| 15 | 34986143 | 35019351 | 33209 | 0 | SIM3 |
| 15 | 35019352 | 57968741 | 22949390 | 2 | SIM3 |
| 15 | 57968742 | 73687198 | 15718457 | 3 | SIM3 |
| 15 | 73687199 | 74638178 | 950980 | 2 | SIM3 |
| 15 | 74638179 | 74671468 | 33290 | 0 | SIM3 |
| 15 | 74671469 | 87631459 | 12959991 | 2 | SIM3 |
| 15 | 87631460 | 102531392 | 14899933 | 3 | SIM3 |
| 16 | 1 | 77687110 | 77687110 | 4 | SIM3 |
| 16 | 77687111 | 79168147 | 1481037 | 2 | SIM3 |
| 16 | 79168148 | 90354753 | 11186606 | 4 | SIM3 |
| 17 | 1 | 20168792 | 20168791 | 2 | SIM3 |
| 17 | 20168793 | 26681489 | 6512697 | 3 | SIM3 |
| 17 | 26681490 | 58197867 | 31516378 | 4 | SIM3 |
| 17 | 58197868 | 58214691 | 16824 | 2 | SIM3 |
| 17 | 58214692 | 76164878 | 17950187 | 6 | SIM3 |
| 17 | 76164879 | 81195210 | 5030332 | 4 | SIM3 |
| 18 | 1 | 17618497 | 17618496 | 5 | SIM3 |
| 18 | 17618498 | 28179178 | 10560681 | 6 | SIM3 |
| 18 | 28179179 | 78077248 | 49898070 | 2 | SIM3 |
| 19 | 1 | 59128983 | 59128983 | 2 | SIM3 |
| 20 | 1 | 12412856 | 12412855 | 0 | SIM3 |
| 20 | 12412857 | 12798147 | 385291 | 7 | SIM3 |
| 20 | 12798148 | 13491847 | 693700 | 13 | SIM3 |
| 20 | 13491848 | 13518914 | 27067 | 11 | SIM3 |
| 20 | 13518915 | 22861497 | 9342583 | 2 | SIM3 |
| 20 | 22861498 | 63025520 | 40164023 | 3 | SIM3 |
| 21 | 1 | 33946887 | 33946887 | 2 | SIM3 |
| 21 | 33946888 | 35041897 | 1095010 | 0 | SIM3 |
| 21 | 35041898 | 42598146 | 7556249 | 2 | SIM3 |
| 21 | 42598147 | 42947987 | 349841 | 0 | SIM3 |
| 21 | 42947988 | 48129895 | 5181908 | 2 | SIM3 |
| 22 | 1 | 51304566 | 51304566 | 3 | SIM3 |
| X | 1 | 13259847 | 13259846 | 5 | SIM3 |
| X | 13259848 | 155270560 | 1.42E+08 | 4 | SIM3 |
| 1 | 1 | 57684115 | 57684115 | 2 | SIM4 |
| 1 | 57684116 | 57734618 | 50503 | 3 | SIM4 |
| 1 | 57734619 | 76418224 | 18683606 | 2 | SIM4 |
| 1 | 76418225 | 76434816 | 16592 | 0 | SIM4 |
| 1 | 76434817 | 100643148 | 24208332 | 2 | SIM4 |
| 1 | 1.01E+08 | 115316842 | 14673694 | 1 | SIM4 |
| 1 | 1.15E+08 | 171423868 | 56107026 | 2 | SIM4 |
| 1 | 1.71E+08 | 221368483 | 49944615 | 4 | SIM4 |
| 1 | 2.21E+08 | 249250621 | 27882138 | 2 | SIM4 |
| 2 | 1 | 56415681 | 56415681 | 3 | SIM4 |
| 2 | 56415682 | 59114683 | 2699002 | 5 | SIM4 |
| 2 | 59114684 | 87126484 | 28011801 | 3 | SIM4 |
| 2 | 87126485 | 116478056 | 29351572 | 4 | SIM4 |
| 2 | 1.16E+08 | 116504156 | 26100 | 0 | SIM4 |
| 2 | 1.17E+08 | 126416388 | 9912232 | 4 | SIM4 |
| 2 | 1.26E+08 | 179364180 | 52947792 | 3 | SIM4 |
| 2 | 1.79E+08 | 179461475 | 97295 | 2 | SIM4 |
| 2 | 1.79E+08 | 225147582 | 45686107 | 3 | SIM4 |
| 2 | 2.25E+08 | 243199373 | 18051791 | 4 | SIM4 |
| 3 | 1 | 39417582 | 39417581 | 6 | SIM4 |
| 3 | 39417583 | 45147558 | 5729976 | 5 | SIM4 |
| 3 | 45147559 | 47416856 | 2269298 | 4 | SIM4 |
| 3 | 47416857 | 52135688 | 4718832 | 5 | SIM4 |
| 3 | 52135689 | 86416386 | 34280698 | 4 | SIM4 |
| 3 | 86416387 | 112416750 | 26000364 | 3 | SIM4 |
| 3 | 1.12E+08 | 112454168 | 37418 | 0 | SIM4 |
| 3 | 1.12E+08 | 198022430 | 85568262 | 3 | SIM4 |
| 4 | 1 | 56416854 | 56416853 | 3 | SIM4 |
| 4 | 56416855 | 112415638 | 55998784 | 2 | SIM4 |
| 4 | 1.12E+08 | 112641684 | 226046 | 3 | SIM4 |
| 4 | 1.13E+08 | 135851464 | 23209780 | 2 | SIM4 |
| 4 | 1.36E+08 | 135894152 | 42688 | 0 | SIM4 |
| 4 | 1.36E+08 | 164417528 | 28523376 | 2 | SIM4 |
| 4 | 1.64E+08 | 164814525 | 396997 | 3 | SIM4 |
| 4 | 1.65E+08 | 191154276 | 26339751 | 2 | SIM4 |
| 5 | 1 | 23451962 | 23451962 | 2 | SIM4 |
| 5 | 23451963 | 23481508 | 29546 | 0 | SIM4 |
| 5 | 23481509 | 56146852 | 32665344 | 2 | SIM4 |
| 5 | 56146853 | 56241865 | 95013 | 5 | SIM4 |
| 5 | 56241866 | 102416850 | 46174985 | 2 | SIM4 |
| 5 | 1.02E+08 | 102814582 | 397732 | 6 | SIM4 |
| 5 | 1.03E+08 | 180915260 | 78100678 | 2 | SIM4 |
| 6 | 1 | 27168578 | 27168578 | 4 | SIM4 |
| 6 | 27168579 | 27241862 | 73284 | 2 | SIM4 |
| 6 | 27241863 | 102416689 | 75174827 | 4 | SIM4 |
| 6 | 1.02E+08 | 102464187 | 47498 | 2 | SIM4 |
| 6 | 1.02E+08 | 156413661 | 53949474 | 4 | SIM4 |
| 6 | 1.56E+08 | 156416486 | 2825 | 0 | SIM4 |
| 6 | 1.56E+08 | 171115067 | 14698581 | 4 | SIM4 |
| 7 | 1 | 65106578 | 65106578 | 2 | SIM4 |
| 7 | 65106579 | 72416384 | 7309806 | 4 | SIM4 |
| 7 | 72416385 | 73106586 | 690202 | 6 | SIM4 |
| 7 | 73106587 | 103416031 | 30309445 | 2 | SIM4 |
| 7 | 1.03E+08 | 106498166 | 3082135 | 3 | SIM4 |
| 7 | 1.06E+08 | 109461843 | 2963677 | 4 | SIM4 |
| 7 | 1.09E+08 | 116142568 | 6680725 | 3 | SIM4 |
| 7 | 1.16E+08 | 117103684 | 961116 | 5 | SIM4 |
| 7 | 1.17E+08 | 159138663 | 42034979 | 3 | SIM4 |
| 8 | 1 | 23107656 | 23107655 | 1 | SIM4 |
| 8 | 23107657 | 72418763 | 49311107 | 2 | SIM4 |
| 8 | 72418764 | 72441824 | 23061 | 0 | SIM4 |
| 8 | 72441825 | 141416418 | 68974594 | 7 | SIM4 |
| 8 | 1.41E+08 | 146364022 | 4947604 | 2 | SIM4 |
| 9 | 1 | 12418637 | 12418636 | 3 | SIM4 |
| 9 | 12418638 | 21016381 | 8597744 | 2 | SIM4 |
| 9 | 21016382 | 22438613 | 1422232 | 0 | SIM4 |
| 9 | 22438614 | 24438613 | 2000000 | 1 | SIM4 |
| 9 | 24438614 | 50000000 | 25561387 | 2 | SIM4 |
| 9 | 50000001 | 76143684 | 26143684 | 3 | SIM4 |
| 9 | 76143685 | 114168375 | 38024691 | 2 | SIM4 |
| 9 | 1.14E+08 | 135146386 | 20978011 | 3 | SIM4 |
| 9 | 1.35E+08 | 141213431 | 6067045 | 2 | SIM4 |
| 10 | 1 | 89623382 | 89623382 | 4 | SIM4 |
| 10 | 89623383 | 89731687 | 108305 | 0 | SIM4 |
| 10 | 89731688 | 135534747 | 45803060 | 4 | SIM4 |
| 11 | 1 | 26431684 | 26431684 | 1 | SIM4 |
| 11 | 26431685 | 38418630 | 11986946 | 2 | SIM4 |
| 11 | 38418631 | 101416803 | 62998173 | 1 | SIM4 |
| 11 | 1.01E+08 | 101421418 | 4615 | 0 | SIM4 |
| 11 | 1.01E+08 | 135006516 | 33585098 | 1 | SIM4 |
| 12 | 1 | 21416384 | 21416384 | 3 | SIM4 |
| 12 | 21416385 | 48418752 | 27002368 | 4 | SIM4 |
| 12 | 48418753 | 101416389 | 52997637 | 5 | SIM4 |
| 12 | 1.01E+08 | 103410831 | 1994442 | 7 | SIM4 |
| 12 | 1.03E+08 | 133851895 | 30441064 | 3 | SIM4 |
| 13 | 1 | 24412314 | 24412313 | 1 | SIM4 |
| 13 | 24412315 | 81416834 | 57004520 | 2 | SIM4 |
| 13 | 81416835 | 86182331 | 4765497 | 3 | SIM4 |
| 13 | 86182332 | 115169878 | 28987547 | 2 | SIM4 |
| 14 | 1 | 46415633 | 46415633 | 2 | SIM4 |
| 14 | 46415634 | 47041862 | 626229 | 0 | SIM4 |
| 14 | 47041863 | 107349540 | 60307678 | 2 | SIM4 |
| 15 | 1 | 102531392 | 1.03E+08 | 2 | SIM4 |
| 16 | 1 | 27418631 | 27418630 | 5 | SIM4 |
| 16 | 27418632 | 57143684 | 29725053 | 2 | SIM4 |
| 16 | 57143685 | 58418364 | 1274680 | 3 | SIM4 |
| 16 | 58418365 | 59641825 | 1223461 | 4 | SIM4 |
| 16 | 59641826 | 60141863 | 500038 | 3 | SIM4 |
| 16 | 60141864 | 90354753 | 30212890 | 2 | SIM4 |
| 17 | 1 | 12614358 | 12614358 | 2 | SIM4 |
| 17 | 12614359 | 14314683 | 1700325 | 4 | SIM4 |
| 17 | 14314684 | 16418648 | 2103965 | 2 | SIM4 |
| 17 | 16418649 | 16444816 | 26168 | 0 | SIM4 |
| 17 | 16444817 | 18413504 | 1968688 | 2 | SIM4 |
| 17 | 18413505 | 21483160 | 3069656 | 4 | SIM4 |
| 17 | 21483161 | 62418614 | 40935454 | 2 | SIM4 |
| 17 | 62418615 | 79413684 | 16995070 | 5 | SIM4 |
| 17 | 79413685 | 81195210 | 1781526 | 2 | SIM4 |
| 18 | 1 | 78077248 | 78077248 | 2 | SIM4 |
| 19 | 1 | 41148675 | 41148675 | 2 | SIM4 |
| 19 | 41148676 | 41183465 | 34790 | 0 | SIM4 |
| 19 | 41183466 | 59128983 | 17945518 | 2 | SIM4 |
| 20 | 1 | 34418864 | 34418864 | 2 | SIM4 |
| 20 | 34418865 | 34464864 | 46000 | 0 | SIM4 |
| 20 | 34464865 | 41786031 | 7321167 | 2 | SIM4 |
| 20 | 41786032 | 52418468 | 10632437 | 3 | SIM4 |
| 20 | 52418469 | 63025520 | 10607052 | 2 | SIM4 |
| 21 | 1 | 48129895 | 48129895 | 2 | SIM4 |
| 22 | 1 | 31416834 | 31416834 | 2 | SIM4 |
| 22 | 31416835 | 31504186 | 87352 | 0 | SIM4 |
| 22 | 31504187 | 51304566 | 19800380 | 2 | SIM4 |
| X | 1 | 155270560 | 1.55E+08 | 2 | SIM4 |
| 1 | 1 | 27146869 | 27146869 | 4 | SIM5 |
| 1 | 27146870 | 27434108 | 287239 | 3 | SIM5 |
| 1 | 27434109 | 57146870 | 29712762 | 4 | SIM5 |
| 1 | 57146871 | 57183574 | 36704 | 5 | SIM5 |
| 1 | 57183575 | 92367184 | 35183610 | 4 | SIM5 |
| 1 | 92367185 | 92756413 | 389229 | 5 | SIM5 |
| 1 | 92756414 | 130000000 | 37243587 | 4 | SIM5 |
| 1 | 1.3E+08 | 249250621 | 1.19E+08 | 5 | SIM5 |
| 2 | 1 | 57412073 | 57412073 | 3 | SIM5 |
| 2 | 57412074 | 57446589 | 34516 | 0 | SIM5 |
| 2 | 57446590 | 76347152 | 18900563 | 3 | SIM5 |
| 2 | 76347153 | 76594169 | 247017 | 6 | SIM5 |
| 2 | 76594170 | 108436801 | 31842632 | 3 | SIM5 |
| 2 | 1.08E+08 | 243199373 | 1.35E+08 | 2 | SIM5 |
| 3 | 1 | 82416834 | 82416833 | 2 | SIM5 |
| 3 | 82416835 | 82434163 | 17329 | 0 | SIM5 |
| 3 | 82434164 | 116934014 | 34499851 | 4 | SIM5 |
| 3 | 1.17E+08 | 117341874 | 407860 | 8 | SIM5 |
| 3 | 1.17E+08 | 169420478 | 52078604 | 4 | SIM5 |
| 3 | 1.69E+08 | 169694200 | 273722 | 8 | SIM5 |
| 3 | 1.7E+08 | 198022430 | 28328230 | 4 | SIM5 |
| 4 | 1 | 23108171 | 23108170 | 1 | SIM5 |
| 4 | 23108172 | 23176146 | 67975 | 0 | SIM5 |
| 4 | 23176147 | 42148865 | 18972719 | 1 | SIM5 |
| 4 | 42148866 | 42238614 | 89749 | 2 | SIM5 |
| 4 | 42238615 | 50000000 | 7761386 | 1 | SIM5 |
| 4 | 50000001 | 62416748 | 12416748 | 2 | SIM5 |
| 4 | 62416749 | 66714688 | 4297940 | 1 | SIM5 |
| 4 | 66714689 | 89417594 | 22702906 | 2 | SIM5 |
| 4 | 89417595 | 89641257 | 223663 | 6 | SIM5 |
| 4 | 89641258 | 167417658 | 77776401 | 2 | SIM5 |
| 4 | 1.67E+08 | 169107542 | 1689884 | 4 | SIM5 |
| 4 | 1.69E+08 | 191154276 | 22046734 | 2 | SIM5 |
| 5 | 1 | 49470178 | 49470177 | 6 | SIM5 |
| 5 | 49470179 | 56417810 | 6947632 | 2 | SIM5 |
| 5 | 56417811 | 164471350 | 1.08E+08 | 5 | SIM5 |
| 5 | 1.64E+08 | 164492468 | 21118 | 0 | SIM5 |
| 5 | 1.64E+08 | 175417097 | 10924629 | 4 | SIM5 |
| 5 | 1.75E+08 | 180915260 | 5498163 | 6 | SIM5 |
| 6 | 1 | 44648710 | 44648710 | 4 | SIM5 |
| 6 | 44648711 | 44943178 | 294468 | 6 | SIM5 |
| 6 | 44943179 | 46413687 | 1470509 | 4 | SIM5 |
| 6 | 46413688 | 46713605 | 299918 | 8 | SIM5 |
| 6 | 46713606 | 47401893 | 688288 | 4 | SIM5 |
| 6 | 47401894 | 47539810 | 137917 | 6 | SIM5 |
| 6 | 47539811 | 48366532 | 826722 | 4 | SIM5 |
| 6 | 48366533 | 48413697 | 47165 | 6 | SIM5 |
| 6 | 48413698 | 49638714 | 1225017 | 4 | SIM5 |
| 6 | 49638715 | 49684168 | 45454 | 8 | SIM5 |
| 6 | 49684169 | 49714168 | 30000 | 0 | SIM5 |
| 6 | 49714169 | 51867378 | 2153210 | 4 | SIM5 |
| 6 | 51867379 | 51894136 | 26758 | 2 | SIM5 |
| 6 | 51894137 | 52114896 | 220760 | 4 | SIM5 |
| 6 | 52114897 | 52144188 | 29292 | 6 | SIM5 |
| 6 | 52144189 | 115419641 | 63275453 | 4 | SIM5 |
| 6 | 1.15E+08 | 115483684 | 64043 | 0 | SIM5 |
| 6 | 1.15E+08 | 171115067 | 55631383 | 4 | SIM5 |
| 7 | 1 | 12618765 | 12618764 | 5 | SIM5 |
| 7 | 12618766 | 15368401 | 2749636 | 3 | SIM5 |
| 7 | 15368402 | 41681407 | 26313006 | 5 | SIM5 |
| 7 | 41681408 | 48614898 | 6933491 | 3 | SIM5 |
| 7 | 48614899 | 50698789 | 2083891 | 5 | SIM5 |
| 7 | 50698790 | 52618765 | 1919976 | 3 | SIM5 |
| 7 | 52618766 | 53769418 | 1150653 | 4 | SIM5 |
| 7 | 53769419 | 54641386 | 871968 | 3 | SIM5 |
| 7 | 54641387 | 55086487 | 445101 | 4 | SIM5 |
| 7 | 55086488 | 55846837 | 760350 | 3 | SIM5 |
| 7 | 55846838 | 56489674 | 642837 | 4 | SIM5 |
| 7 | 56489675 | 56986441 | 496767 | 3 | SIM5 |
| 7 | 56986442 | 57136877 | 150436 | 6 | SIM5 |
| 7 | 57136878 | 57396847 | 259970 | 4 | SIM5 |
| 7 | 57396848 | 59846641 | 2449794 | 3 | SIM5 |
| 7 | 59846642 | 65648179 | 5801538 | 4 | SIM5 |
| 7 | 65648180 | 76168407 | 10520228 | 3 | SIM5 |
| 7 | 76168408 | 79387521 | 3219114 | 1 | SIM5 |
| 7 | 79387522 | 83681474 | 4293953 | 3 | SIM5 |
| 7 | 83681475 | 84416897 | 735423 | 7 | SIM5 |
| 7 | 84416898 | 86484169 | 2067272 | 6 | SIM5 |
| 7 | 86484170 | 92463841 | 5979672 | 8 | SIM5 |
| 7 | 92463842 | 95348744 | 2884903 | 3 | SIM5 |
| 7 | 95348745 | 103964876 | 8616132 | 5 | SIM5 |
| 7 | 1.04E+08 | 113684479 | 9719603 | 1 | SIM5 |
| 7 | 1.14E+08 | 121863717 | 8179238 | 3 | SIM5 |
| 7 | 1.22E+08 | 122168494 | 304777 | 4 | SIM5 |
| 7 | 1.22E+08 | 125687694 | 3519200 | 3 | SIM5 |
| 7 | 1.26E+08 | 125874618 | 186924 | 4 | SIM5 |
| 7 | 1.26E+08 | 129468179 | 3593561 | 3 | SIM5 |
| 7 | 1.29E+08 | 131368798 | 1900619 | 2 | SIM5 |
| 7 | 1.31E+08 | 142368416 | 10999618 | 3 | SIM5 |
| 7 | 1.42E+08 | 142526841 | 158425 | 4 | SIM5 |
| 7 | 1.43E+08 | 143486787 | 959946 | 3 | SIM5 |
| 7 | 1.43E+08 | 153486787 | 10000000 | 1 | SIM5 |
| 7 | 1.53E+08 | 157648315 | 4161528 | 3 | SIM5 |
| 7 | 1.58E+08 | 159138663 | 1490348 | 5 | SIM5 |
| 8 | 1 | 34683777 | 34683777 | 2 | SIM5 |
| 8 | 34683778 | 34699415 | 15638 | 0 | SIM5 |
| 8 | 34699416 | 49418678 | 14719263 | 2 | SIM5 |
| 8 | 49418679 | 146364022 | 96945344 | 3 | SIM5 |
| 9 | 1 | 21941062 | 21941062 | 2 | SIM5 |
| 9 | 21941063 | 22086481 | 145419 | 0 | SIM5 |
| 9 | 22086482 | 141213431 | 1.19E+08 | 2 | SIM5 |
| 10 | 1 | 135534747 | 1.36E+08 | 3 | SIM5 |
| 11 | 1 | 82684178 | 82684177 | 4 | SIM5 |
| 11 | 82684179 | 89614874 | 6930696 | 2 | SIM5 |
| 11 | 89614875 | 91938617 | 2323743 | 4 | SIM5 |
| 11 | 91938618 | 102499878 | 10561261 | 2 | SIM5 |
| 11 | 1.02E+08 | 113619879 | 11120001 | 4 | SIM5 |
| 11 | 1.14E+08 | 113649089 | 29210 | 2 | SIM5 |
| 11 | 1.14E+08 | 113681618 | 32529 | 0 | SIM5 |
| 11 | 1.14E+08 | 135006516 | 21324898 | 4 | SIM5 |
| 12 | 1 | 26614087 | 26614086 | 2 | SIM5 |
| 12 | 26614088 | 29491064 | 2876977 | 3 | SIM5 |
| 12 | 29491065 | 31418657 | 1927593 | 5 | SIM5 |
| 12 | 31418658 | 34618077 | 3199420 | 3 | SIM5 |
| 12 | 34618078 | 43946189 | 9328112 | 3 | SIM5 |
| 12 | 43946190 | 73146830 | 29200641 | 3 | SIM5 |
| 12 | 73146831 | 73172489 | 25659 | 0 | SIM5 |
| 12 | 73172490 | 133851895 | 60679406 | 3 | SIM5 |
| 13 | 1 | 31547945 | 31547945 | 2 | SIM5 |
| 13 | 31547946 | 31574698 | 26753 | 0 | SIM5 |
| 13 | 31574699 | 48631789 | 17057091 | 2 | SIM5 |
| 13 | 48631790 | 62614807 | 13983018 | 3 | SIM5 |
| 13 | 62614808 | 92648107 | 30033300 | 2 | SIM5 |
| 13 | 92648108 | 92758741 | 110634 | 7 | SIM5 |
| 13 | 92758742 | 115169878 | 22411137 | 2 | SIM5 |
| 14 | 1 | 44687179 | 44687178 | 1 | SIM5 |
| 14 | 44687180 | 64568864 | 19881685 | 2 | SIM5 |
| 14 | 64568865 | 64591687 | 22823 | 1 | SIM5 |
| 14 | 64591688 | 86671872 | 22080185 | 2 | SIM5 |
| 14 | 86671873 | 86691546 | 19674 | 0 | SIM5 |
| 14 | 86691547 | 107349540 | 20657994 | 2 | SIM5 |
| 15 | 1 | 37166878 | 37166878 | 2 | SIM5 |
| 15 | 37166879 | 37681439 | 514561 | 3 | SIM5 |
| 15 | 37681440 | 78968417 | 41286978 | 2 | SIM5 |
| 15 | 78968418 | 78980174 | 11757 | 0 | SIM5 |
| 15 | 78980175 | 102531392 | 23551218 | 2 | SIM5 |
| 16 | 1 | 62861478 | 62861478 | 1 | SIM5 |
| 16 | 62861479 | 62946871 | 85393 | 0 | SIM5 |
| 16 | 62946872 | 90354753 | 27407882 | 1 | SIM5 |
| 17 | 1 | 18681479 | 18681478 | 2 | SIM5 |
| 17 | 18681480 | 21986498 | 3305019 | 4 | SIM5 |
| 17 | 21986499 | 22169887 | 183389 | 7 | SIM5 |
| 17 | 22169888 | 24169887 | 2000000 | 3 | SIM5 |
| 17 | 24169888 | 37486197 | 13316310 | 2 | SIM5 |
| 17 | 37486198 | 48861486 | 11375289 | 5 | SIM5 |
| 17 | 48861487 | 49197864 | 336378 | 9 | SIM5 |
| 17 | 49197865 | 56684987 | 7487123 | 4 | SIM5 |
| 17 | 56684988 | 57361527 | 676540 | 8 | SIM5 |
| 17 | 57361528 | 59418698 | 2057171 | 2 | SIM5 |
| 17 | 59418699 | 81195210 | 21776512 | 1 | SIM5 |
| 18 | 1 | 78077248 | 78077248 | 2 | SIM5 |
| 19 | 1 | 16678987 | 16678986 | 2 | SIM5 |
| 19 | 16678988 | 16694681 | 15694 | 4 | SIM5 |
| 19 | 16694682 | 17648156 | 953475 | 2 | SIM5 |
| 19 | 17648157 | 59128983 | 41480827 | 4 | SIM5 |
| 20 | 1 | 39000000 | 38999999 | 2 | SIM5 |
| 20 | 39000001 | 63025520 | 24025520 | 3 | SIM5 |
| 21 | 1 | 48129895 | 48129895 | 2 | SIM5 |
| 22 | 1 | 20976481 | 20976480 | 3 | SIM5 |
| 22 | 20976482 | 24683787 | 3707306 | 2 | SIM5 |
| 22 | 24683788 | 28468198 | 3784411 | 3 | SIM5 |
| 22 | 28468199 | 42946817 | 14478619 | 2 | SIM5 |
| 22 | 42946818 | 42971648 | 24831 | 0 | SIM5 |
| 22 | 42971649 | 51304566 | 8332918 | 2 | SIM5 |
| X | 1 | 155270560 | 1.55E+08 | 2 | SIM5 |

**Supplementary Table 1.** Ground truth copy number profiles for the five synthetic genomes used in the study.
