## Supplementary Table 2 for "Robust somatic copy number estimation using coarse-to-fine segmentation"

| **chr** | **pos** | **end** | **CN** | **tool** | **call** |
| --- | --- | --- | --- | --- | --- |
| 1 | 532372 | 709195 | 1 | ploidetect | F |
| 1 | 709195 | 9319439 | 2.5 | ploidetect | T |
| 1 | 762601 | 9340454 | 2.224793 | battenberg | T |
| 1 | 9340695 | 9399461 | 2.680401 | battenberg | T |
| 1 | 9319439 | 9402773 | 3 | ploidetect | T |
| 1 | 9402773 | 17008262 | 2.5 | ploidetect | F |
| 1 | 17008262 | 17295816 | 3 | ploidetect | F |
| 1 | 13116 | 25590806 | 3 | ascatngs | T |
| 1 | 17295816 | 25606269 | 2.5 | ploidetect | F |
| 1 | 25592695 | 25660621 | 1 | ascatngs | F |
| 1 | 25606269 | 25664908 | 1 | ploidetect | F |
| 1 | 9402518 | 87304095 | 2.266711 | battenberg | F |
| 1 | 25663049 | 87333107 | 3 | ascatngs | F |
| 1 | 25664908 | 87337222 | 2.5 | ploidetect | F |
| 1 | 1 | 87371000 | 2.5352 | purple | F |
| 1 | 87337222 | 1.21E+08 | 2 | ploidetect | F |
| 1 | 1.21E+08 | 1.21E+08 | 2.5 | ploidetect | F |
| 1 | 1.21E+08 | 1.21E+08 | 3 | ploidetect | F |
| 1 | 1.21E+08 | 1.21E+08 | 2.5 | ploidetect | F |
| 1 | 87334960 | 1.21E+08 | 2 | ascatngs | F |
| 1 | 1.21E+08 | 1.21E+08 | 1 | ascatngs | F |
| 1 | 87430864 | 1.21E+08 | 1.783752 | battenberg | F |
| 1 | 1.21E+08 | 1.21E+08 | 2 | ascatngs | F |
| 1 | 1.21E+08 | 1.21E+08 | 1 | ascatngs | F |
| 1 | 1.21E+08 | 1.43E+08 | 2 | ploidetect | F |
| 1 | 1.43E+08 | 1.43E+08 | 2.5 | ploidetect | F |
| 1 | 1.43E+08 | 1.43E+08 | 3 | ploidetect | F |
| 1 | 1.43E+08 | 1.45E+08 | 3.5 | ploidetect | F |
| 1 | 1.45E+08 | 1.49E+08 | 4 | ploidetect | F |
| 1 | 1.49E+08 | 1.49E+08 | 3 | ploidetect | T |
| 1 | 1.43E+08 | 1.49E+08 | 4 | ascatngs | F |
| 1 | 1.49E+08 | 1.49E+08 | 0 | ploidetect | T |
| 1 | 1.49E+08 | 1.49E+08 | 1 | ploidetect | F |
| 1 | 1.49E+08 | 1.49E+08 | 2.5 | ploidetect | T |
| 1 | 1.49E+08 | 1.49E+08 | 1 | ploidetect | T |
| 1 | 1.49E+08 | 1.49E+08 | 3 | ploidetect | F |
| 1 | 1.49E+08 | 1.49E+08 | 2.5 | ploidetect | F |
| 1 | 1.49E+08 | 1.49E+08 | 1 | ascatngs | T |
| 1 | 1.49E+08 | 1.5E+08 | 2 | ascatngs | F |
| 1 | 1.49E+08 | 1.5E+08 | 1 | ploidetect | F |
| 1 | 1.5E+08 | 1.5E+08 | 2.5 | ploidetect | F |
| 1 | 1.23E+08 | 2.25E+08 | 3.9528 | purple | T |
| 1 | 2.25E+08 | 2.25E+08 | 6.1201 | purple | T |
| 1 | 1.5E+08 | 2.49E+08 | 4 | ploidetect | T |
| 1 | 2.49E+08 | 2.49E+08 | 0 | ploidetect | T |
| 1 | 1.45E+08 | 2.49E+08 | 3.525646 | battenberg | NA |
| 1 | 2.49E+08 | 2.49E+08 | 3.75 | ploidetect | NA |
| 1 | 1.5E+08 | 2.49E+08 | 4 | ascatngs | NA |
| 1 | 2.25E+08 | 2.49E+08 | 3.9241 | purple | NA |
| 10 | 93528 | 7048329 | 1.753823 | battenberg | T |
| 10 | 0 | 7059579 | 2 | ploidetect | T |
| 10 | 7086117 | 7112848 | 2.549018 | battenberg | T |
| 10 | 7059579 | 7132041 | 3 | ploidetect | T |
| 10 | 1 | 7627000 | 1.9922 | purple | F |
| 10 | 7132041 | 7634291 | 2 | ploidetect | F |
| 10 | 7139607 | 7639031 | 1.708478 | battenberg | F |
| 10 | 60969 | 7908897 | 2 | ascatngs | F |
| 10 | 7913058 | 18281176 | 3 | ascatngs | F |
| 10 | 18282421 | 18354127 | 2 | ascatngs | F |
| 10 | 18355456 | 19272870 | 3 | ascatngs | F |
| 10 | 19274196 | 19354244 | 2 | ascatngs | F |
| 10 | 19355552 | 20390193 | 3 | ascatngs | F |
| 10 | 20391290 | 20498946 | 2 | ascatngs | F |
| 10 | 20500592 | 20628270 | 3 | ascatngs | F |
| 10 | 20629321 | 20656157 | 2 | ascatngs | F |
| 10 | 20657567 | 23801555 | 3 | ascatngs | F |
| 10 | 23802824 | 23868365 | 2 | ascatngs | F |
| 10 | 23869533 | 25701341 | 3 | ascatngs | F |
| 10 | 25702556 | 25832211 | 2 | ascatngs | F |
| 10 | 25833267 | 26187217 | 3 | ascatngs | F |
| 10 | 26188455 | 26399456 | 2 | ascatngs | F |
| 10 | 26400644 | 26435330 | 3 | ascatngs | F |
| 10 | 26437506 | 26628775 | 2 | ascatngs | F |
| 10 | 26630030 | 27544931 | 3 | ascatngs | F |
| 10 | 27545984 | 27581394 | 2 | ascatngs | F |
| 10 | 27582640 | 31053914 | 3 | ascatngs | F |
| 10 | 31055117 | 31080284 | 2 | ascatngs | F |
| 10 | 31081548 | 33275518 | 3 | ascatngs | F |
| 10 | 33277540 | 33370878 | 2 | ascatngs | F |
| 10 | 33371929 | 33385294 | 3 | ascatngs | T |
| 10 | 7654501 | 33385946 | 2.245987 | battenberg | T |
| 10 | 7634291 | 33388794 | 2.5 | ploidetect | T |
| 10 | 7627001 | 33389000 | 2.4929 | purple | T |
| 10 | 33386226 | 39087270 | 1.770978 | battenberg | F |
| 10 | 33386314 | 39129538 | 2 | ascatngs | F |
| 10 | 33388794 | 42383348 | 2 | ploidetect | F |
| 10 | 42383348 | 42384206 | 3 | ploidetect | F |
| 10 | 42384206 | 42389372 | 2 | ploidetect | F |
| 10 | 42389372 | 42389513 | 3 | ploidetect | F |
| 10 | 42389513 | 42391573 | 2 | ploidetect | F |
| 10 | 42391573 | 42391873 | 3 | ploidetect | F |
| 10 | 42391873 | 42393673 | 2 | ploidetect | F |
| 10 | 42393673 | 42394550 | 3 | ploidetect | F |
| 10 | 39130629 | 42446847 | 3 | ascatngs | F |
| 10 | 42394550 | 42694510 | 2 | ploidetect | F |
| 10 | 42694510 | 42718824 | 3 | ploidetect | F |
| 10 | 42718824 | 89699626 | 2 | ploidetect | T |
| 10 | 89699626 | 89713578 | 0 | ploidetect | T |
| 10 | 89713578 | 1.28E+08 | 2 | ploidetect | F |
| 10 | 1.28E+08 | 1.28E+08 | 3 | ploidetect | F |
| 10 | 42397373 | 1.35E+08 | 1.774563 | battenberg | NA |
| 10 | 1.28E+08 | 1.35E+08 | 2 | ploidetect | F |
| 10 | 1.35E+08 | 1.35E+08 | 2.5 | ploidetect | NA |
| 10 | 42448304 | 1.36E+08 | 2 | ascatngs | NA |
| 10 | 40754935 | 1.36E+08 | 1.9907 | purple | NA |
| 11 | 196944 | 48337732 | 2.76674 | battenberg | F |
| 11 | 48346681 | 48355838 | 2.490138 | battenberg | F |
| 11 | 48424597 | 50056195 | 2.604486 | battenberg | F |
| 11 | 50105688 | 50136994 | 2.528976 | battenberg | F |
| 11 | 50137201 | 51565191 | 2.514855 | battenberg | F |
| 11 | 1 | 53144204 | 3.0924 | purple | F |
| 11 | 87268 | 55362513 | 3 | ascatngs | F |
| 11 | 55364529 | 55439799 | 6 | ascatngs | F |
| 11 | 0 | 80777395 | 3 | ploidetect | T |
| 11 | 55440868 | 80781809 | 3 | ascatngs | T |
| 11 | 53144205 | 80785000 | 2.9519 | purple | T |
| 11 | 80777395 | 80788723 | 2 | ploidetect | F |
| 11 | 80782982 | 80810788 | 2 | ascatngs | F |
| 11 | 80788723 | 81093030 | 1.5 | ploidetect | T |
| 11 | 80785001 | 81094000 | 1.5889 | purple | T |
| 11 | 81093030 | 1.17E+08 | 2.75 | ploidetect | F |
| 11 | 1.17E+08 | 1.17E+08 | 3 | ploidetect | F |
| 11 | 1.17E+08 | 1.35E+08 | 2.75 | ploidetect | NA |
| 11 | 80811894 | 1.35E+08 | 3 | ascatngs | NA |
| 11 | 54794237 | 1.35E+08 | 2.506544 | battenberg | NA |
| 11 | 81094001 | 1.35E+08 | 2.7869 | purple | NA |
| 12 | 0 | 9623918 | 3 | ploidetect | F |
| 12 | 9623918 | 9735474 | 2 | ploidetect | F |
| 12 | 9735474 | 11226224 | 3 | ploidetect | F |
| 12 | 11226224 | 11247269 | 2 | ploidetect | F |
| 12 | 1 | 36356693 | 2.9841 | purple | F |
| 12 | 191525 | 38974970 | 2.668416 | battenberg | T |
| 12 | 38975282 | 39968287 | 2.741969 | battenberg | F |
| 12 | 39970412 | 1.34E+08 | 2.650656 | battenberg | NA |
| 12 | 11247269 | 1.34E+08 | 3 | ploidetect | F |
| 12 | 1.34E+08 | 1.34E+08 | 2 | ploidetect | NA |
| 12 | 68657 | 1.34E+08 | 3 | ascatngs | NA |
| 12 | 36356694 | 1.34E+08 | 2.9747 | purple | NA |
| 13 | 1 | 17499999 | 3.1035 | purple | F |
| 13 | 0 | 41509558 | 3.25 | ploidetect | F |
| 13 | 41509558 | 41632789 | 4 | ploidetect | F |
| 13 | 19448984 | 1.15E+08 | 2.828689 | battenberg | NA |
| 13 | 41632789 | 1.15E+08 | 3.25 | ploidetect | F |
| 13 | 1.15E+08 | 1.15E+08 | 3 | ploidetect | NA |
| 13 | 19020095 | 1.15E+08 | 3 | ascatngs | NA |
| 13 | 17500000 | 1.15E+08 | 3.1035 | purple | NA |
| 14 | 1 | 17499999 | 2.9813 | purple | NA |
| 14 | 0 | 19121312 | 4 | ploidetect | F |
| 14 | 19121312 | 19410334 | 3 | ploidetect | F |
| 14 | 19410334 | 19438699 | 1 | ploidetect | T |
| 14 | 19438699 | 19480862 | 0 | ploidetect | T |
| 14 | 19480862 | 19786138 | 1 | ploidetect | F |
| 14 | 19786138 | 19796251 | 3 | ploidetect | F |
| 14 | 19796251 | 20105693 | 1 | ploidetect | F |
| 14 | 19000060 | 20424926 | 2 | ascatngs | F |
| 14 | 20105693 | 20425034 | 1.75 | ploidetect | F |
| 14 | 20426398 | 43819536 | 3 | ascatngs | T |
| 14 | 20425034 | 43823750 | 3 | ploidetect | T |
| 14 | 43820732 | 44247493 | 4 | ascatngs | T |
| 14 | 43823750 | 44253539 | 4 | ploidetect | T |
| 14 | 44253539 | 1.06E+08 | 3 | ploidetect | F |
| 14 | 1.06E+08 | 1.06E+08 | 4 | ploidetect | F |
| 14 | 44248654 | 1.06E+08 | 3 | ascatngs | F |
| 14 | 1.06E+08 | 1.06E+08 | 11 | ploidetect | F |
| 14 | 1.06E+08 | 1.06E+08 | 11 | ascatngs | F |
| 14 | 1.06E+08 | 1.06E+08 | 111 | ascatngs | F |
| 14 | 1.06E+08 | 1.06E+08 | 19 | ploidetect | F |
| 14 | 1.06E+08 | 1.06E+08 | 16 | ploidetect | F |
| 14 | 1.06E+08 | 1.06E+08 | 21 | ascatngs | F |
| 14 | 1.06E+08 | 1.07E+08 | 6 | ascatngs | F |
| 14 | 1.06E+08 | 1.07E+08 | 5 | ploidetect | F |
| 14 | 1.07E+08 | 1.07E+08 | 5 | ascatngs | F |
| 14 | 1.07E+08 | 1.07E+08 | 4 | ploidetect | F |
| 14 | 1.07E+08 | 1.07E+08 | 6 | ascatngs | F |
| 14 | 1.07E+08 | 1.07E+08 | 5 | ploidetect | F |
| 14 | 20475482 | 1.07E+08 | 2.679259 | battenberg | NA |
| 14 | 1.07E+08 | 1.07E+08 | 3 | ploidetect | F |
| 14 | 1.07E+08 | 1.07E+08 | 2 | ploidetect | NA |
| 14 | 1.07E+08 | 1.07E+08 | 3 | ascatngs | NA |
| 14 | 17500000 | 1.07E+08 | 2.9813 | purple | NA |
| 15 | 1 | 18499999 | 3.8063 | purple | F |
| 15 | 20001861 | 20202442 | 2.477869 | battenberg | F |
| 15 | 0 | 20396243 | 3 | ploidetect | F |
| 15 | 20000538 | 22563905 | 3 | ascatngs | F |
| 15 | 22565974 | 23662895 | 4 | ascatngs | F |
| 15 | 20212798 | 23688394 | 3.201466 | battenberg | F |
| 15 | 23664545 | 23718659 | 5 | ascatngs | F |
| 15 | 23719829 | 23803295 | 6 | ascatngs | F |
| 15 | 23804370 | 23816119 | 7 | ascatngs | T |
| 15 | 23817238 | 23940401 | 5 | ascatngs | T |
| 15 | 23688554 | 23985800 | 4.024087 | battenberg | T |
| 15 | 23941678 | 23991669 | 4 | ascatngs | T |
| 15 | 18500000 | 23992000 | 3.8063 | purple | T |
| 15 | 20396243 | 24000834 | 3.75 | ploidetect | T |
| 15 | 24000834 | 28701193 | 2 | ploidetect | F |
| 15 | 28701193 | 28708291 | 3 | ploidetect | F |
| 15 | 23992001 | 84807000 | 2.004 | purple | T |
| 15 | 23994573 | 84808065 | 1.798263 | battenberg | T |
| 15 | 23993038 | 84810480 | 2 | ascatngs | T |
| 15 | 28708291 | 84811154 | 2 | ploidetect | T |
| 15 | 84811774 | 84858253 | 3 | ascatngs | F |
| 15 | 84859270 | 84931022 | 4 | ascatngs | F |
| 15 | 84932436 | 84956894 | 5 | ascatngs | F |
| 15 | 84958034 | 89413243 | 4 | ascatngs | F |
| 15 | 89414295 | 89643573 | 5 | ascatngs | F |
| 15 | 89644938 | 90960620 | 4 | ascatngs | F |
| 15 | 90961813 | 90979121 | 5 | ascatngs | F |
| 15 | 90980980 | 91031048 | 4 | ascatngs | F |
| 15 | 91034456 | 91077809 | 5 | ascatngs | F |
| 15 | 91079465 | 93396686 | 4 | ascatngs | F |
| 15 | 93397763 | 93575153 | 5 | ascatngs | F |
| 15 | 93576190 | 94848183 | 4 | ascatngs | F |
| 15 | 94849637 | 94866787 | 5 | ascatngs | F |
| 15 | 84811774 | 1.02E+08 | 3.542511 | battenberg | NA |
| 15 | 84811154 | 1.02E+08 | 4 | ploidetect | NA |
| 15 | 94867962 | 1.03E+08 | 4 | ascatngs | NA |
| 15 | 84807001 | 1.03E+08 | 3.9939 | purple | NA |
| 16 | 60291 | 32041587 | 4 | ascatngs | F |
| 16 | 0 | 33968278 | 3.25 | ploidetect | F |
| 16 | 83802 | 34173796 | 2.977351 | battenberg | F |
| 16 | 33968278 | 34182022 | 2 | ploidetect | F |
| 16 | 34182022 | 34184604 | 3 | ploidetect | F |
| 16 | 32046359 | 34193941 | 3 | ascatngs | F |
| 16 | 1 | 34197000 | 3.3599 | purple | F |
| 16 | 34176649 | 35282052 | 1.800723 | battenberg | F |
| 16 | 34197001 | 36835800 | 2.0047 | purple | F |
| 16 | 34184604 | 46436895 | 2 | ploidetect | F |
| 16 | 46436895 | 46438259 | 3 | ploidetect | F |
| 16 | 46438259 | 46449201 | 2 | ploidetect | F |
| 16 | 46449201 | 46454802 | 3 | ploidetect | F |
| 16 | 46454802 | 58624326 | 2 | ploidetect | T |
| 16 | 58624326 | 58662799 | 0 | ploidetect | T |
| 16 | 34196008 | 78928026 | 2 | ascatngs | T |
| 16 | 58662799 | 78928178 | 2 | ploidetect | T |
| 16 | 36835801 | 78929000 | 1.9872 | purple | T |
| 16 | 78929186 | 78947725 | 1 | ascatngs | F |
| 16 | 78928178 | 79093743 | 1 | ploidetect | T |
| 16 | 78929001 | 79095000 | 0.9329 | purple | T |
| 16 | 46461292 | 89998957 | 1.769281 | battenberg | NA |
| 16 | 79093743 | 90240935 | 2 | ploidetect | NA |
| 16 | 78948837 | 90292766 | 2 | ascatngs | NA |
| 16 | 79095001 | 90354753 | 2.0131 | purple | NA |
| 17 | 0 | 18350892 | 3 | ploidetect | F |
| 17 | 18350892 | 18447882 | 2 | ploidetect | F |
| 17 | 7293 | 22244184 | 2.722146 | battenberg | F |
| 17 | 1 | 23763005 | 3.0542 | purple | F |
| 17 | 18447882 | 76661368 | 3 | ploidetect | F |
| 17 | 76661368 | 76789525 | 3.5 | ploidetect | F |
| 17 | 76789525 | 79026163 | 3.25 | ploidetect | F |
| 17 | 25282979 | 79998834 | 2.751668 | battenberg | NA |
| 17 | 79026163 | 81133834 | 3 | ploidetect | F |
| 17 | 81133834 | 81154780 | 3.25 | ploidetect | NA |
| 17 | 828 | 81185372 | 3 | ascatngs | NA |
| 17 | 23763006 | 81195210 | 3.1008 | purple | NA |
| 18 | 16444 | 9867933 | 2 | ascatngs | T |
| 18 | 1 | 9868000 | 2.0002 | purple | T |
| 18 | 125371 | 9868204 | 1.797623 | battenberg | T |
| 18 | 0 | 9872201 | 2 | ploidetect | T |
| 18 | 9868700 | 14142120 | 2.243019 | battenberg | F |
| 18 | 9868001 | 16960897 | 2.5359 | purple | F |
| 18 | 9872201 | 78007624 | 2.5 | ploidetect | F |
| 18 | 78007624 | 78011492 | 3 | ploidetect | NA |
| 18 | 9869011 | 78017073 | 3 | ascatngs | NA |
| 18 | 18605053 | 78017073 | 2.228791 | battenberg | NA |
| 18 | 16960898 | 78077248 | 2.4849 | purple | NA |
| 19 | 226776 | 24512434 | 3 | ascatngs | F |
| 19 | 255741 | 24512434 | 2.628738 | battenberg | F |
| 19 | 24515088 | 24592510 | 2 | ascatngs | F |
| 19 | 1 | 26181781 | 2.9633 | purple | F |
| 19 | 0 | 54728629 | 3 | ploidetect | F |
| 19 | 54728629 | 54743667 | 2 | ploidetect | F |
| 19 | 27741144 | 59080607 | 2.660074 | battenberg | NA |
| 19 | 54743667 | 59095127 | 3 | ploidetect | NA |
| 19 | 24594382 | 59118779 | 3 | ascatngs | NA |
| 19 | 26181782 | 59128983 | 2.9648 | purple | NA |
| 2 | 10587 | 89130009 | 3 | ascatngs | F |
| 2 | 0 | 89151715 | 3 | ploidetect | F |
| 2 | 89133443 | 89161322 | 11 | ascatngs | F |
| 2 | 89162353 | 89245827 | 60 | ascatngs | F |
| 2 | 89151715 | 89248228 | 29 | ploidetect | F |
| 2 | 89246942 | 89387414 | 8 | ascatngs | F |
| 2 | 89388973 | 89516771 | 6 | ascatngs | F |
| 2 | 89517897 | 89533272 | 5 | ascatngs | F |
| 2 | 1 | 93826170 | 2.9839 | purple | NA |
| 2 | 55984 | 2.43E+08 | 2.654637 | battenberg | NA |
| 2 | 89248228 | 2.43E+08 | 3 | ploidetect | NA |
| 2 | 89538879 | 2.43E+08 | 3 | ascatngs | NA |
| 2 | 93826171 | 2.43E+08 | 2.977 | purple | NA |
| 20 | 0 | 14960127 | 4 | ploidetect | T |
| 20 | 14960127 | 15000849 | 3 | ploidetect | T |
| 20 | 15000849 | 15014179 | 2 | ploidetect | T |
| 20 | 20000786 | 25734937 | 3.446834 | battenberg | F |
| 20 | 25829761 | 26307077 | 3.459779 | battenberg | NA |
| 20 | 1 | 27869568 | 3.9672 | purple | NA |
| 20 | 29420083 | 29565395 | 3.491752 | battenberg | NA |
| 20 | 29573972 | 62891857 | 3.545471 | battenberg | NA |
| 20 | 15014179 | 62960973 | 4 | ploidetect | NA |
| 20 | 61098 | 62962869 | 4 | ascatngs | NA |
| 20 | 27869569 | 63025520 | 4.0241 | purple | NA |
| 21 | 9411410 | 9554245 | 4 | ascatngs | F |
| 21 | 9555825 | 10597683 | 4 | ascatngs | F |
| 21 | 0 | 10645981 | 3.5 | ploidetect | F |
| 21 | 1 | 12788128 | 2.9547 | purple | NA |
| 21 | 15273817 | 48082324 | 2.664614 | battenberg | NA |
| 21 | 10645981 | 48101658 | 3 | ploidetect | NA |
| 21 | 10598696 | 48119634 | 3 | ascatngs | NA |
| 21 | 12788129 | 48129895 | 2.9547 | purple | NA |
| 22 | 1 | 14499999 | 4.0354 | purple | NA |
| 22 | 0 | 16351953 | 2 | ploidetect | F |
| 22 | 16051249 | 16425814 | 3 | ascatngs | F |
| 22 | 16351953 | 16484899 | 3 | ploidetect | F |
| 22 | 16428299 | 17034805 | 4 | ascatngs | F |
| 22 | 16566779 | 17297558 | 3.434762 | battenberg | F |
| 22 | 17035811 | 17300230 | 4 | ascatngs | T |
| 22 | 17301534 | 23111003 | 4 | ascatngs | F |
| 22 | 23112021 | 23175968 | 8 | ascatngs | F |
| 22 | 23177175 | 23248256 | 7 | ascatngs | F |
| 22 | 16484899 | 33758987 | 4 | ploidetect | T |
| 22 | 33758987 | 33841448 | 3 | ploidetect | T |
| 22 | 17300320 | 49998118 | 3.56483 | battenberg | NA |
| 22 | 33841448 | 51220827 | 4 | ploidetect | NA |
| 22 | 23249354 | 51240820 | 4 | ascatngs | NA |
| 22 | 14500000 | 51304566 | 4.0354 | purple | NA |
| 3 | 0 | 1214009 | 2 | ploidetect | F |
| 3 | 60197 | 1252218 | 2 | ascatngs | F |
| 3 | 1253525 | 1315199 | 1 | ascatngs | F |
| 3 | 1214009 | 1318906 | 1 | ploidetect | F |
| 3 | 60596 | 24533278 | 1.741782 | battenberg | T |
| 3 | 1 | 24566000 | 1.9689 | purple | T |
| 3 | 1318906 | 24566582 | 2 | ploidetect | T |
| 3 | 1316388 | 24570343 | 2 | ascatngs | T |
| 3 | 24572140 | 25331908 | 10 | ascatngs | T |
| 3 | 24566001 | 25332000 | 9.7375 | purple | T |
| 3 | 24566582 | 25332907 | 10 | ploidetect | T |
| 3 | 25332907 | 25396148 | 12 | ploidetect | F |
| 3 | 25332992 | 25400903 | 13 | ascatngs | T |
| 3 | 25332001 | 25401000 | 11.965 | purple | T |
| 3 | 25396148 | 26423822 | 8 | ploidetect | T |
| 3 | 25401001 | 26426000 | 7.7681 | purple | T |
| 3 | 25401957 | 26431473 | 8 | ascatngs | T |
| 3 | 24566631 | 26651227 | 8.033271 | battenberg | T |
| 3 | 26432540 | 26663787 | 12 | ascatngs | T |
| 3 | 26426001 | 26664000 | 11.3967 | purple | T |
| 3 | 26423822 | 26665219 | 12 | ploidetect | T |
| 3 | 26665476 | 60131360 | 4 | ascatngs | T |
| 3 | 26665219 | 60132116 | 4 | ploidetect | T |
| 3 | 60132116 | 60206160 | 3 | ploidetect | T |
| 3 | 60132390 | 60210491 | 3 | ascatngs | T |
| 3 | 60206160 | 60872452 | 4 | ploidetect | T |
| 3 | 60211698 | 60872721 | 4 | ascatngs | T |
| 3 | 26669772 | 60878803 | 3.541077 | battenberg | T |
| 3 | 60891684 | 60977150 | 2.580212 | battenberg | F |
| 3 | 60872452 | 61011596 | 3 | ploidetect | T |
| 3 | 60873734 | 61012867 | 3 | ascatngs | T |
| 3 | 61011596 | 75271622 | 4 | ploidetect | T |
| 3 | 75271622 | 75275197 | 3 | ploidetect | T |
| 3 | 61061196 | 90482099 | 3.538856 | battenberg | F |
| 3 | 26664001 | 92004853 | 3.9519 | purple | NA |
| 3 | 75275197 | 98190081 | 4 | ploidetect | F |
| 3 | 61013985 | 98191073 | 4 | ascatngs | F |
| 3 | 98190081 | 98352142 | 5 | ploidetect | F |
| 3 | 98192785 | 98411452 | 5 | ascatngs | F |
| 3 | 93511746 | 1.98E+08 | 3.541974 | battenberg | NA |
| 3 | 98352142 | 1.98E+08 | 4 | ploidetect | NA |
| 3 | 98413111 | 1.98E+08 | 4 | ascatngs | NA |
| 3 | 92004854 | 1.98E+08 | 3.9635 | purple | NA |
| 4 | 0 | 72600 | 3.75 | ploidetect | F |
| 4 | 72600 | 3567412 | 2 | ploidetect | F |
| 4 | 11961 | 3571410 | 2 | ascatngs | F |
| 4 | 3573108 | 3634358 | 3 | ascatngs | F |
| 4 | 3567412 | 3635870 | 3 | ploidetect | F |
| 4 | 3635870 | 33843849 | 2 | ploidetect | F |
| 4 | 33843849 | 33852597 | 3 | ploidetect | F |
| 4 | 33852597 | 49214360 | 2 | ploidetect | F |
| 4 | 49214360 | 49220460 | 3 | ploidetect | F |
| 4 | 49220460 | 49324296 | 2 | ploidetect | F |
| 4 | 3635907 | 49493257 | 2 | ascatngs | NA |
| 4 | 49324296 | 49509417 | 3.75 | ploidetect | NA |
| 4 | 97884 | 49633309 | 1.821116 | battenberg | NA |
| 4 | 1 | 51160116 | 2.0303 | purple | NA |
| 4 | 49494265 | 1.91E+08 | 4 | ascatngs | F |
| 4 | 1.91E+08 | 1.91E+08 | 3 | ascatngs | F |
| 4 | 53639492 | 1.91E+08 | 3.457224 | battenberg | NA |
| 4 | 49509417 | 1.91E+08 | 4 | ploidetect | NA |
| 4 | 1.91E+08 | 1.91E+08 | 4 | ascatngs | NA |
| 4 | 51160117 | 1.91E+08 | 3.8326 | purple | NA |
| 5 | 0 | 17516577 | 2 | ploidetect | F |
| 5 | 17516577 | 17648553 | 0 | ploidetect | T |
| 5 | 17648553 | 21477579 | 2 | ploidetect | F |
| 5 | 21477579 | 21570863 | 2.75 | ploidetect | F |
| 5 | 1 | 28788000 | 1.9478 | purple | T |
| 5 | 21570863 | 28788579 | 2 | ploidetect | T |
| 5 | 28788001 | 28963000 | 0.9173 | purple | T |
| 5 | 28788579 | 28964166 | 1 | ploidetect | T |
| 5 | 850203 | 46405572 | 1.76449 | battenberg | NA |
| 5 | 28963001 | 47905640 | 1.9874 | purple | NA |
| 5 | 28964166 | 70241340 | 2 | ploidetect | F |
| 5 | 70241340 | 70325097 | 1 | ploidetect | F |
| 5 | 70325097 | 1.8E+08 | 2 | ploidetect | T |
| 5 | 1.8E+08 | 1.8E+08 | 0 | ploidetect | T |
| 5 | 49441623 | 1.81E+08 | 1.771757 | battenberg | NA |
| 5 | 11882 | 1.81E+08 | 2 | ascatngs | NA |
| 5 | 1.8E+08 | 1.81E+08 | 2 | ploidetect | NA |
| 5 | 47905641 | 1.81E+08 | 1.9867 | purple | NA |
| 6 | 0 | 25993604 | 4 | ploidetect | F |
| 6 | 202777 | 57189265 | 3.384993 | battenberg | F |
| 6 | 100116 | 57379829 | 4 | ascatngs | F |
| 6 | 57381003 | 57569567 | 4 | ascatngs | F |
| 6 | 57211138 | 58773345 | 3.188819 | battenberg | F |
| 6 | 1 | 60330165 | 3.786 | purple | NA |
| 6 | 25993604 | 65297995 | 3.75 | ploidetect | T |
| 6 | 60330166 | 65298000 | 3.6293 | purple | T |
| 6 | 57570848 | 65298045 | 4 | ascatngs | T |
| 6 | 61882658 | 65338640 | 3.234424 | battenberg | T |
| 6 | 65297995 | 1.09E+08 | 2.75 | ploidetect | F |
| 6 | 1.09E+08 | 1.09E+08 | 3 | ploidetect | F |
| 6 | 65342926 | 1.71E+08 | 2.459922 | battenberg | NA |
| 6 | 1.09E+08 | 1.71E+08 | 2.75 | ploidetect | NA |
| 6 | 65299056 | 1.71E+08 | 3 | ascatngs | NA |
| 6 | 65298001 | 1.71E+08 | 2.7537 | purple | NA |
| 7 | 0 | 56558896 | 4 | ploidetect | F |
| 7 | 56558896 | 56564501 | 3 | ploidetect | F |
| 7 | 81237 | 57470343 | 3.546665 | battenberg | T |
| 7 | 56564501 | 57470857 | 4 | ploidetect | T |
| 7 | 24220 | 57470970 | 4 | ascatngs | T |
| 7 | 57470857 | 57482989 | 2 | ploidetect | T |
| 7 | 57471815 | 57494706 | 1.632279 | battenberg | T |
| 7 | 57471972 | 57495828 | 2 | ascatngs | T |
| 7 | 57497027 | 57584913 | 3.464423 | battenberg | F |
| 7 | 57612544 | 58051318 | 3.3908 | battenberg | NA |
| 7 | 1 | 59554330 | 3.9705 | purple | NA |
| 7 | 57496874 | 77978486 | 4 | ascatngs | T |
| 7 | 57482989 | 77980103 | 4 | ploidetect | T |
| 7 | 61071378 | 77981388 | 3.499623 | battenberg | T |
| 7 | 77980103 | 78081980 | 3 | ploidetect | T |
| 7 | 78081980 | 78196608 | 4 | ploidetect | T |
| 7 | 77982085 | 78256643 | 3.020111 | battenberg | T |
| 7 | 78196608 | 78257353 | 3 | ploidetect | T |
| 7 | 77980510 | 78257877 | 3 | ascatngs | T |
| 7 | 78258968 | 1.04E+08 | 4 | ascatngs | T |
| 7 | 78257353 | 1.04E+08 | 4 | ploidetect | T |
| 7 | 1.04E+08 | 1.05E+08 | 6 | ascatngs | F |
| 7 | 1.05E+08 | 1.05E+08 | 7 | ascatngs | T |
| 7 | 1.04E+08 | 1.05E+08 | 6 | ploidetect | T |
| 7 | 78258968 | 1.26E+08 | 3.590317 | battenberg | T |
| 7 | 1.05E+08 | 1.26E+08 | 4 | ascatngs | T |
| 7 | 59554331 | 1.26E+08 | 3.9423 | purple | T |
| 7 | 1.05E+08 | 1.26E+08 | 4 | ploidetect | T |
| 7 | 1.26E+08 | 1.26E+08 | 1.75 | ploidetect | T |
| 7 | 1.26E+08 | 1.26E+08 | 2 | ascatngs | T |
| 7 | 1.26E+08 | 1.26E+08 | 1.677115 | battenberg | T |
| 7 | 1.26E+08 | 1.26E+08 | 1.8679 | purple | T |
| 7 | 1.26E+08 | 1.26E+08 | 3.422134 | battenberg | T |
| 7 | 1.26E+08 | 1.26E+08 | 3.8082 | purple | T |
| 7 | 1.26E+08 | 1.26E+08 | 4 | ascatngs | T |
| 7 | 1.26E+08 | 1.26E+08 | 4 | ploidetect | T |
| 7 | 1.26E+08 | 1.31E+08 | 6 | ploidetect | F |
| 7 | 1.31E+08 | 1.31E+08 | 7 | ploidetect | F |
| 7 | 1.31E+08 | 1.34E+08 | 6 | ploidetect | T |
| 7 | 1.34E+08 | 1.34E+08 | 5 | ploidetect | T |
| 7 | 1.34E+08 | 1.42E+08 | 6 | ploidetect | F |
| 7 | 1.42E+08 | 1.42E+08 | 8 | ploidetect | F |
| 7 | 1.42E+08 | 1.43E+08 | 6 | ploidetect | F |
| 7 | 1.43E+08 | 1.43E+08 | 4 | ploidetect | T |
| 7 | 1.26E+08 | 1.44E+08 | 5.362929 | battenberg | F |
| 7 | 1.43E+08 | 1.44E+08 | 6 | ploidetect | F |
| 7 | 1.26E+08 | 1.44E+08 | 6.0039 | purple | F |
| 7 | 1.26E+08 | 1.44E+08 | 6 | ascatngs | F |
| 7 | 1.44E+08 | 1.51E+08 | 3.457713 | battenberg | T |
| 7 | 1.44E+08 | 1.51E+08 | 3.8605 | purple | T |
| 7 | 1.44E+08 | 1.51E+08 | 4 | ploidetect | T |
| 7 | 1.44E+08 | 1.51E+08 | 4 | ascatngs | T |
| 7 | 1.51E+08 | 1.52E+08 | 2 | ploidetect | F |
| 7 | 1.52E+08 | 1.52E+08 | 3 | ploidetect | F |
| 7 | 1.51E+08 | 1.59E+08 | 1.734785 | battenberg | NA |
| 7 | 1.51E+08 | 1.59E+08 | 2 | ascatngs | NA |
| 7 | 1.52E+08 | 1.59E+08 | 2 | ploidetect | NA |
| 7 | 1.51E+08 | 1.59E+08 | 1.9687 | purple | NA |
| 8 | 146791 | 43819896 | 2.960762 | battenberg | F |
| 8 | 1 | 45338886 | 3.2886 | purple | NA |
| 8 | 11774 | 56116482 | 4 | ascatngs | F |
| 8 | 0 | 1.46E+08 | 3.25 | ploidetect | F |
| 8 | 1.46E+08 | 1.46E+08 | 3 | ploidetect | NA |
| 8 | 46858994 | 1.46E+08 | 2.928841 | battenberg | NA |
| 8 | 56118810 | 1.46E+08 | 3 | ascatngs | NA |
| 8 | 45338887 | 1.46E+08 | 3.2875 | purple | NA |
| 9 | 0 | 23352441 | 2 | ploidetect | F |
| 9 | 23352441 | 23380231 | 3 | ploidetect | F |
| 9 | 23380231 | 28060192 | 2 | ploidetect | T |
| 9 | 28060192 | 28160086 | 1 | ploidetect | T |
| 9 | 28160086 | 30336210 | 2 | ploidetect | T |
| 9 | 203937 | 30338681 | 1.76786 | battenberg | T |
| 9 | 1 | 30339000 | 1.9626 | purple | T |
| 9 | 10469 | 30389018 | 2 | ascatngs | T |
| 9 | 30340701 | 38772381 | 3.353704 | battenberg | F |
| 9 | 30390419 | 40777931 | 4 | ascatngs | F |
| 9 | 40779579 | 41244986 | 3 | ascatngs | F |
| 9 | 41247719 | 42542485 | 4 | ascatngs | F |
| 9 | 42549097 | 43398864 | 3 | ascatngs | F |
| 9 | 43400052 | 43736147 | 4 | ascatngs | F |
| 9 | 43737545 | 43871367 | 5 | ascatngs | F |
| 9 | 43873554 | 44083789 | 3 | ascatngs | F |
| 9 | 44084972 | 44158043 | 5 | ascatngs | F |
| 9 | 44159853 | 44875147 | 4 | ascatngs | F |
| 9 | 44867223 | 45098181 | 3.610092 | battenberg | F |
| 9 | 44877072 | 45354882 | 5 | ascatngs | F |
| 9 | 45356325 | 45730599 | 4 | ascatngs | F |
| 9 | 45733681 | 46119593 | 5 | ascatngs | F |
| 9 | 30339001 | 48867678 | 3.7833 | purple | NA |
| 9 | 46121628 | 68477816 | 4 | ascatngs | F |
| 9 | 68479207 | 70939686 | 3 | ascatngs | F |
| 9 | 71083668 | 1.41E+08 | 3.230152 | battenberg | NA |
| 9 | 30336210 | 1.41E+08 | 3.75 | ploidetect | F |
| 9 | 1.41E+08 | 1.41E+08 | 3 | ploidetect | NA |
| 9 | 70943506 | 1.41E+08 | 4 | ascatngs | NA |
| 9 | 48867679 | 1.41E+08 | 3.629 | purple | NA |
| X | 0 | 158616 | 1 | ploidetect | F |
| X | 158616 | 2698351 | 2 | ploidetect | F |
| X | 1455089 | 2699246 | 1.682355 | battenberg | F |
| X | 2698351 | 31196695 | 4 | ploidetect | T |
| X | 31196695 | 31217760 | 2 | ploidetect | T |
| X | 31217760 | 31298802 | 4 | ploidetect | T |
| X | 2699555 | 31300162 | 4 | ascatngs | T |
| X | 1 | 31301000 | 1.9688 | purple | T |
| X | 31298802 | 32034718 | 2 | ploidetect | T |
| X | 31302979 | 32039342 | 2 | ascatngs | T |
| X | 32034718 | 32078559 | 4 | ploidetect | T |
| X | 32078559 | 32092541 | 2 | ploidetect | T |
| X | 32041161 | 32097405 | 4 | ascatngs | T |
| X | 32092541 | 32199353 | 0 | ploidetect | T |
| X | 32100414 | 32200753 | 0 | ascatngs | T |
| X | 31301001 | 32293000 | 0.9862 | purple | T |
| X | 32201893 | 32293091 | 2 | ascatngs | T |
| X | 32199353 | 32293183 | 2 | ploidetect | T |
| X | 32293001 | 60132011 | 1.9501 | purple | NA |
| X | 32294253 | 1.55E+08 | 4 | ascatngs | NA |
| X | 32293183 | 1.55E+08 | 4 | ploidetect | F |
| X | 1.55E+08 | 1.55E+08 | 1.69882 | battenberg | NA |
| X | 1.55E+08 | 1.55E+08 | 2 | ploidetect | NA |
| X | 60132012 | 1.55E+08 | 1.9288 | purple | NA |

**Supplementary Table 2.** Manual review of COLO829 CNV Calls. Every CNV breakpoint identified in COLO829 by ascatNGS, Battenberg, Ploidetect, and PURPLE was subjected to manual review of raw alignment data. Judged truth of each breakpoint is indicated in the “call” column. T = True, breakpoint assessed as a true positive. F = False, breakpoint assessed as a false positive. NA = Not applicable, breakpoint either corresponds to chromosome end or within centromere. Breakpoints were assessed using the “end” column.
