## Supplementary Table 3 for "Robust somatic copy number estimation using coarse-to-fine segmentation"

| **gene_id** | **Mutation** |
| --- | --- |
| TP53 | HOMD |
| CDKN2A | HOMD |
| CDKN2B | HOMD |
| PTEN | HOMD |
| NF1 | HOMD |
| RB1 | HOMD |
| ARID1A | HOMD |
| RAD51B | HOMD |
| SMAD4 | HOMD |
| STK11 | HOMD |
| BRCA1 | HOMD |
| BRCA2 | HOMD |
| FANCD2 | HOMD |
| FAT1 | HOMD |
| SMARCB1 | HOMD |
| MYC | AMP |
| AURKA | AMP |
| ERBB2 | AMP |
| CCND1 | AMP |
| EGFR | AMP |
| FGFR1 | AMP |
| CDK6 | AMP |

**Supplementary Table 3.** Recurrently homozygously deleted (HOMD) and amplified (AMP) genes in cancer used for assessing sensitivity for variants in CNV software.
