## Supplementary Table 4 for "Robust somatic copy number estimation using coarse-to-fine segmentation"

| **type** | **case** | **hgnc_symbol** | **Call** |
| --- | --- | --- | --- |
| homd | POG003 | PTEN | T |
| homd | POG004 | CDKN2AB | T |
| homd | POG007 | TP53 | T |
| amp | POG007 | CCND1 | T |
| amp | POG012 | EGFR | F |
| amp | POG022 | CCND1 | T |
| amp | POG023 | MYC | T |
| amp | POG031 | AURKA | T |
| amp | POG033 | EGFR | T |
| homd | POG040 | PTEN | T |
| homd | POG046 | PTEN | T |
| homd | POG049 | RB1 | T |
| amp | POG050 | MYC | T |
| homd | POG056 | PTEN | T |
| amp | POG057 | AURKA | T |
| amp | POG063 | EGFR | F |
| amp | POG074 | CCND1 | F |
| amp | POG092 | MYC | F |
| amp | POG099 | MYC | T |
| homd | POG1008 | STK11 | T |
| homd | POG1020 | SMAD4 | F |
| amp | POG1023 | EGFR | F |
| homd | POG1034 | STK11 | T |
| homd | POG1034 | CDKN2AB | T |
| amp | POG104 | AURKA | F |
| amp | POG1062 | EGFR | F |
| homd | POG1068 | PTEN | F |
| homd | POG1071 | CDKN2AB | T |
| amp | POG108 | AURKA | F |
| amp | POG1083 | CDK6 | F |
| homd | POG1087 | ARID1A | F |
| homd | POG1087 | NF1 | F |
| amp | POG1093 | FGFR1 | T |
| homd | POG1098 | BRCA2 | T |
| amp | POG1098 | MYC | T |
| homd | POG1103 | BRCA2 | T |
| homd | POG1108 | ARID1A | F |
| homd | POG1108 | SMAD4 | F |
| homd | POG1129 | CDKN2AB | T |
| amp | POG114 | MYC | T |
| amp | POG139 | CDK6 | F |
| homd | POG140 | RB1 | T |
| amp | POG143 | MYC | T |
| amp | POG143 | AURKA | T |
| amp | POG143 | EGFR | F |
| homd | POG147 | CDKN2AB | T |
| homd | POG151 | CDKN2AB | T |
| homd | POG174 | SMAD4 | T |
| amp | POG181 | EGFR | F |
| amp | POG181 | CDK6 | F |
| amp | POG183 | ERBB2 | T |
| homd | POG185 | PTEN | T |
| homd | POG189 | CDKN2AB | T |
| amp | POG197 | EGFR | F |
| homd | POG198 | SMAD4 | T |
| amp | POG199 | AURKA | F |
| amp | POG216 | MYC | F |
| amp | POG221 | FGFR1 | T |
| homd | POG227 | CDKN2AB | T |
| amp | POG227 | CCND1 | F |
| amp | POG235 | AURKA | F |
| homd | POG243 | CDKN2AB | T |
| amp | POG248 | FGFR1 | F |
| homd | POG253 | FAT1 | F |
| amp | POG253 | FGFR1 | F |
| amp | POG253 | MYC | T |
| homd | POG258 | RB1 | T |
| homd | POG262 | STK11 | F |
| homd | POG262 | SMARCB1 | F |
| amp | POG266 | CDK6 | T |
| amp | POG280 | AURKA | F |
| amp | POG286 | EGFR | F |
| amp | POG295 | AURKA | T |
| homd | POG297 | FAT1 | F |
| homd | POG298 | CDKN2AB | T |
| amp | POG303 | FGFR1 | T |
| homd | POG326 | PTEN | T |
| amp | POG330 | CDK6 | F |
| amp | POG342 | AURKA | T |
| homd | POG346 | CDKN2AB | T |
| homd | POG359 | CDKN2AB | T |
| homd | POG365 | RAD51B | F |
| amp | POG365 | MYC | F |
| amp | POG371 | MYC | T |
| homd | POG373 | RB1 | F |
| homd | POG373 | BRCA2 | F |
| homd | POG382 | CDKN2AB | T |
| homd | POG382 | ARID1A | F |
| homd | POG384 | NF1 | T |
| amp | POG388 | CCND1 | T |
| homd | POG396 | CDKN2AB | T |
| homd | POG399 | CDKN2AB | T |
| homd | POG400 | SMAD4 | T |
| homd | POG405 | SMAD4 | T |
| homd | POG405 | FAT1 | T |
| amp | POG408 | EGFR | T |
| homd | POG416 | PTEN | T |
| amp | POG416 | MYC | T |
| homd | POG420 | CDKN2AB | T |
| amp | POG425 | MYC | F |
| amp | POG436 | CCND1 | T |
| homd | POG449 | CDKN2AB | T |
| homd | POG457 | CDKN2AB | T |
| amp | POG460 | MYC | T |
| homd | POG465 | CDKN2AB | T |
| amp | POG477 | ERBB2 | T |
| homd | POG478 | CDKN2AB | T |
| homd | POG486 | PTEN | T |
| homd | POG490 | RAD51B | F |
| amp | POG491 | AURKA | F |
| homd | POG495 | FAT1 | F |
| homd | POG497 | CDKN2AB | T |
| homd | POG509 | PTEN | T |
| amp | POG511 | ERBB2 | F |
| homd | POG513 | BRCA1 | F |
| homd | POG513 | NF1 | F |
| amp | POG549 | AURKA | T |
| amp | POG559 | MYC | T |
| homd | POG580 | PTEN | F |
| amp | POG588 | MYC | T |
| amp | POG594 | MYC | F |
| homd | POG598 | CDKN2AB | T |
| amp | POG604 | MYC | T |
| amp | POG605 | FGFR1 | T |
| homd | POG608 | STK11 | T |
| homd | POG613 | RB1 | T |
| amp | POG623 | AURKA | T |
| amp | POG623 | MYC | T |
| homd | POG643 | CDKN2AB | T |
| homd | POG662 | SMARCB1 | F |
| homd | POG662 | ARID1A | F |
| homd | POG663 | RB1 | T |
| homd | POG666 | CDKN2AB | T |
| homd | POG677 | CDKN2AB | F |
| homd | POG677 | SMARCB1 | F |
| homd | POG691 | CDKN2AB | T |
| homd | POG694 | PTEN | T |
| homd | POG713 | CDKN2AB | T |
| homd | POG714 | NF1 | F |
| amp | POG716 | CDK6 | F |
| amp | POG717 | MYC | T |
| amp | POG717 | FGFR1 | F |
| homd | POG719 | NF1 | T |
| amp | POG720 | CCND1 | F |
| amp | POG728 | ERBB2 | T |
| amp | POG731 | MYC | T |
| amp | POG745 | ERBB2 | T |
| homd | POG748 | RB1 | T |
| homd | POG754 | CDKN2AB | T |
| amp | POG754 | MYC | T |
| amp | POG763 | MYC | F |
| amp | POG766 | MYC | F |
| amp | POG768 | EGFR | F |
| amp | POG768 | CCND1 | T |
| homd | POG776 | SMARCB1 | F |
| homd | POG777 | BRCA2 | F |
| homd | POG777 | FAT1 | F |
| amp | POG777 | CCND1 | F |
| amp | POG778 | EGFR | T |
| amp | POG795 | AURKA | T |
| amp | POG803 | EGFR | F |
| amp | POG804 | AURKA | T |
| homd | POG811 | ARID1A | F |
| amp | POG814 | EGFR | F |
| amp | POG816 | FGFR1 | T |
| homd | POG818 | TP53 | F |
| homd | POG818 | BRCA1 | F |
| homd | POG818 | ARID1A | F |
| amp | POG821 | CCND1 | F |
| amp | POG827 | AURKA | F |
| homd | POG834 | STK11 | F |
| homd | POG834 | FANCD2 | F |
| amp | POG834 | FGFR1 | F |
| amp | POG835 | EGFR | T |
| amp | POG835 | MYC | T |
| homd | POG847 | STK11 | F |
| homd | POG875 | TP53 | F |
| homd | POG875 | RB1 | F |
| homd | POG875 | SMARCB1 | F |
| homd | POG875 | BRCA1 | F |
| amp | POG880 | EGFR | F |
| amp | POG885 | CDK6 | F |
| homd | POG890 | CDKN2AB | T |
| homd | POG901 | FAT1 | F |
| amp | POG910 | CDK6 | T |
| amp | POG918 | CCND1 | F |
| amp | POG925 | MYC | T |
| amp | POG926 | EGFR | T |
| homd | POG931 | CDKN2AB | T |
| amp | POG931 | MYC | T |
| amp | POG947 | EGFR | F |
| amp | POG947 | CDK6 | F |
| amp | POG958 | AURKA | F |
| amp | POG963 | MYC | F |
| homd | POG969 | SMAD4 | T |
| homd | POG970 | CDKN2AB | T |
| homd | POG992 | FANCD2 | F |
| amp | POG992 | EGFR | F |
| amp | POG996 | MYC | T |
| amp | POG998 | CDK6 | F |

**Supplementary Table 4.** Manually reviewed CNV calls in POG. Rows represent a randomly selected cancer gene CNV that was identified within the POG cohort. “Call” column represents best assessed truthiness of the call based on manual review. T = True, call assessed as a true positive. F = False, call assessed as a false positive.
