## Supplementary Table 5 for "Robust somatic copy number estimation using coarse-to-fine segmentation"

| **type** | **case** | **hgnc_symbol** | **Chr** | **Band** | **Start (hg19)** | **End (hg19)** | **Reviewer 1** | **Reviewer 2** |
| --- | --- | --- | --- | --- | --- | --- | --- | --- |
| amp | POG022 | CCND1 | chr11 | q13.3 | 69455855 | 69469242 | Yes | Yes |
| amp | POG023 | MYC | chr8 | q24.21 | 128747680 | 128753674 | Yes | Yes |
| amp | POG033 | EGFR | chr7 | p11.2 | 55086714 | 55324313 | Yes | Yes? |
| homd | POG046 | PTEN | chr10 | q23.31 | 89622870 | 89731687 | Yes | Yes |
| amp | POG050 | MYC | chr8 | q24.21 | 128747680 | 128753674 | Yes | Yes |
| homd | POG056 | PTEN | chr10 | q23.31 | 89622870 | 89731687 | Yes | Yes |
| amp | POG057 | AURKA | chr20 | q13.2 | 54944445 | 54967393 | Yes | Yes? |
| amp | POG063 | EGFR | chr7 | p11.2 | 55086714 | 55324313 | No | No |
| amp | POG092 | MYC | chr8 | q24.21 | 128747680 | 128753674 | No | No |
| amp | POG099 | MYC | chr8 | q24.21 | 128747680 | 128753674 | Yes | Yes? |
| homd | POG1020 | SMAD4 | chr18 | q21.2 | 48494410 | 48611415 | No | No |
| amp | POG1023 | EGFR | chr7 | p11.2 | 55086714 | 55324313 | No | No |
| homd | POG1034 | STK11 | chr19 | p13.3 | 1189406 | 1228428 | Yes | Yes |
| homd | POG1034 | CDKN2AB | chr9 | p21.3 | 21967751 | 22009362 | Yes | Yes |
| amp | POG104 | AURKA | chr20 | q13.2 | 54944445 | 54967393 | No | No |
| amp | POG1062 | EGFR | chr7 | p11.2 | 55086714 | 55324313 | No | No? |
| homd | POG1068 | PTEN | chr10 | q23.31 | 89622870 | 89731687 | No | No |
| amp | POG1093 | FGFR1 | chr8 | p11.23 | 38268656 | 38326352 | Yes | Yes? |
| homd | POG1098 | BRCA2 | chr13 | q13.1 | 32889611 | 32973805 | Yes | Yes |
| amp | POG1098 | MYC | chr8 | q24.21 | 128747680 | 128753674 | Yes | Yes |
| homd | POG1108 | ARID1A | chr1 | p36.11 | 27022524 | 27108595 | No | No |
| homd | POG140 | RB1 | chr13 | q14.2 | 48877887 | 49056122 | Yes | Yes |
| amp | POG143 | AURKA | chr20 | q13.2 | 54944445 | 54967393 | Yes | Yes? |
| amp | POG143 | MYC | chr8 | q24.21 | 128747680 | 128753674 | Yes | Yes |
| homd | POG151 | CDKN2AB | chr9 | p21.3 | 21967751 | 22009362 | Yes | Yes |
| amp | POG181 | EGFR | chr7 | p11.2 | 55086714 | 55324313 | No | No |
| homd | POG185 | PTEN | chr10 | q23.31 | 89622870 | 89731687 | Yes | Yes |
| homd | POG189 | CDKN2AB | chr9 | p21.3 | 21967751 | 22009362 | Yes | Yes |
| homd | POG198 | SMAD4 | chr18 | q21.2 | 48494410 | 48611415 | Yes | Yes |
| homd | POG227 | CDKN2AB | chr9 | p21.3 | 21967751 | 22009362 | Yes | Yes |
| amp | POG235 | AURKA | chr20 | q13.2 | 54944445 | 54967393 | No | No |
| homd | POG243 | CDKN2AB | chr9 | p21.3 | 21967751 | 22009362 | Yes | Yes |
| amp | POG248 | FGFR1 | chr8 | p11.23 | 38268656 | 38326352 | No | No |
| homd | POG258 | RB1 | chr13 | q14.2 | 48877887 | 49056122 | Yes | Yes |
| homd | POG262 | SMARCB1 | chr22 | q11.23 | 24129150 | 24176703 | No | No |
| amp | POG286 | EGFR | chr7 | p11.2 | 55086714 | 55324313 | No | No |
| homd | POG298 | CDKN2AB | chr9 | p21.3 | 21967751 | 22009362 | Yes | Yes |
| homd | POG326 | PTEN | chr10 | q23.31 | 89622870 | 89731687 | Yes | Yes |
| amp | POG342 | AURKA | chr20 | q13.2 | 54944445 | 54967393 | Yes | Yes |
| homd | POG346 | CDKN2AB | chr9 | p21.3 | 21967751 | 22009362 | Yes | Yes |
| homd | POG359 | CDKN2AB | chr9 | p21.3 | 21967751 | 22009362 | Yes | Yes |
| homd | POG365 | RAD51B | chr14 | q24.1 | 68286496 | 69196935 | No | No |
| amp | POG365 | MYC | chr8 | q24.21 | 128747680 | 128753674 | No | No |
| amp | POG371 | MYC | chr8 | q24.21 | 128747680 | 128753674 | Yes | Yes |
| homd | POG373 | BRCA2 | chr13 | q13.1 | 32889611 | 32973805 | No | No |
| homd | POG373 | RB1 | chr13 | q14.2 | 48877887 | 49056122 | No | No |
| homd | POG382 | CDKN2AB | chr9 | p21.3 | 21967751 | 22009362 | Yes | Yes |
| homd | POG384 | NF1 | chr17 | q11.2 | 29421945 | 29709134 | Yes | Yes |
| homd | POG396 | CDKN2AB | chr9 | p21.3 | 21967751 | 22009362 | Yes | Yes |
| homd | POG400 | SMAD4 | chr18 | q21.2 | 48494410 | 48611415 | Yes | Yes |

**Supplementary Table 5.** Audit of manually reviewed CNVs in POG. We recruited a second reviewer to perform manual review of cancer-related CNVs in POG. 50 total CNVs were reviewed. Reviewer 1 was the original reviewer whose full results are shown in Supplementary Table 4. Reviewer 2 was the new reviewer. Yes indicates that the CNV was assessed as a true positive, No indicates that the CNV was assessed as false positive.
