## Supplementary Table 6 for "Robust somatic copy number estimation using coarse-to-fine segmentation"

| **gene_id** | **n_deleted** | **n_exons** | **chr** | **start** | **end** | **p_adj** | **cases** |
| --- | --- | --- | --- | --- | --- | --- | --- |
| PLCH2 | 27 | 27 | 1 | 2363768 | 2438457 | 0 | 8 |
| PANK4 | 19 | 19 | 1 | 2438458 | 2517369 | 0 | 7 |
| TTC34 | 8 | 8 | 1 | 2539739 | 2984588 | 0 | 7 |
| MEGF6 | 37 | 37 | 1 | 3385196 | 3530070 | 0 | 5 |
| LRP1B | 19 | 91 | 2 | 1.42E+08 | 1.42E+08 | 0 | 12 |
| LINC01107 | 1 | 3 | 2 | 2.39E+08 | 2.39E+08 | 0 | 7 |
| FHIT | 7 | 22 | 3 | 59760906 | 61205399 | 0 | 46 |
| MIR548BB | 1 | 1 | 3 | 60594232 | 60610342 | 0 | 10 |
| NAALADL2 |  |  | 3 | 1.75E+08 | 1.75E+08 | 0.00352 | 6 |
| TACC3 | 6 | 16 | 4 | 1741397 | 1750554 | 0 | 5 |
| FGFR3 | 22 | 22 | 4 | 1751959 | 1810873 | 0 | 6 |
| NAT8L | 3 | 3 | 4 | 2059181 | 2076823 | 0 | 5 |
| POLN | 7 | 26 | 4 | 2069491 | 2175700 | 2E-05 | 6 |
| UGT2B17 | 6 | 13 | 4 | 69381669 | 69484501 | 0 | 6 |
| CCSER1 | 3 | 13 | 4 | 91584177 | 91984830 | 6.84E-06 | 6 |
| HLA-H | 8 | 64 | 6 | 29854383 | 29893472 | 0 | 5 |
| HCG4B | 1 | 7 | 6 | 29856426 | 29904817 | 0 | 5 |
| HLA-DRB5 | 6 | 11 | 6 | 32478885 | 32501473 | 0 | 5 |
| PRKN | 2 | 12 | 6 | 1.63E+08 | 1.63E+08 | 0.000193 | 5 |
| AFDN-DT | 2 | 2 | 6 | 1.68E+08 | 1.68E+08 | 0 | 5 |
| AFDN | 44 | 44 | 6 | 1.68E+08 | 1.68E+08 | 0 | 5 |
| KIF25 | 9 | 9 | 6 | 1.68E+08 | 1.68E+08 | 0 | 5 |
| FRMD1 | 20 | 20 | 6 | 1.68E+08 | 1.68E+08 | 0 | 5 |
| UNCX | 3 | 3 | 7 | 1269420 | 1278004 | 0 | 5 |
| TARP | 4 | 4 | 7 | 38295706 | 38362646 | 0 | 6 |
| IMMP2L | 3 | 13 | 7 | 1.1E+08 | 1.11E+08 | 0.000148 | 5 |
| MTRNR2L6 | 1 | 1 | 7 | 1.42E+08 | 1.42E+08 | 0 | 5 |
| PRSS1 | 5 | 5 | 7 | 1.42E+08 | 1.42E+08 | 0 | 6 |
| PRSS3P2 | 5 | 5 | 7 | 1.42E+08 | 1.42E+08 | 0 | 7 |
| OR4F21 | 1 | 1 | 8 | 0 | 236023 | 0 | 12 |
| RPL23AP53 | 4 | 4 | 8 | 0 | 236023 | 0 | 12 |
| ZNF596 | 13 | 13 | 8 | 0 | 236023 | 0 | 12 |
| FAM87A | 4 | 4 | 8 | 245367 | 335225 | 0 | 12 |
| FBXO25 | 13 | 13 | 8 | 344783 | 435697 | 0 | 13 |
| TDRP | 7 | 7 | 8 | 435698 | 606080 | 0 | 13 |
| ERICH1 | 8 | 8 | 8 | 435698 | 686869 | 0 | 14 |
| DLGAP2 | 17 | 17 | 8 | 686870 | 1705266 | 0 | 18 |
| LOC401442 | 1 | 1 | 8 | 686870 | 698495 | 0 | 12 |
| LOC105377777 | 2 | 2 | 8 | 698496 | 849692 | 0 | 13 |
| LOC286083 | 3 | 3 | 8 | 1089059 | 1253564 | 0 | 13 |
| DLGAP2-AS1 | 4 | 4 | 8 | 1398720 | 1705266 | 0 | 15 |
| LOC101927752 | 3 | 3 | 8 | 1705267 | 1730798 | 0 | 12 |
| CLN8 | 3 | 3 | 8 | 1705267 | 1747889 | 0 | 12 |
| MIR3674 | 1 | 1 | 8 | 1747890 | 1771455 | 0 | 12 |
| MIR596 | 1 | 1 | 8 | 1747890 | 1771455 | 0 | 12 |
| ARHGEF10 | 32 | 32 | 8 | 1771456 | 1908297 | 0 | 14 |
| LOC101928058 | 4 | 4 | 8 | 1908298 | 1922996 | 0 | 12 |
| KBTBD11-OT1 | 3 | 3 | 8 | 1908298 | 1972299 | 0 | 13 |
| KBTBD11 | 2 | 2 | 8 | 1908298 | 1972299 | 0 | 13 |
| MYOM2 | 37 | 37 | 8 | 1992741 | 2141111 | 0 | 14 |
| MIR7160 | 1 | 1 | 8 | 1992741 | 2040150 | 0 | 13 |
| LOC101927815 | 11 | 11 | 8 | 2324094 | 2743012 | 2.33E-06 | 11 |
| CSMD1 | 70 | 70 | 8 | 2743013 | 4853031 | 0 | 17 |
| C8orf74 | 4 | 4 | 8 | 10528423 | 10593620 | 0.007183 | 5 |
| SOX7 | 2 | 2 | 8 | 10528423 | 10593620 | 0.007183 | 5 |
| LOC102723313 | 1 | 3 | 8 | 10528423 | 10594941 | 0.000882 | 5 |
| FAM66A |  |  | 8 | 12231856 | 12245068 | 0.005453 | 5 |
| LOC649352 | 1 | 1 | 8 | 12231856 | 12245068 | 0.005453 | 5 |
| TUSC3 |  |  | 8 | 15453750 | 15462384 | 0.00537 | 5 |
| ADAM5 | 7 | 15 | 8 | 39225787 | 39346345 | 0 | 17 |
| ADAM3A | 23 | 23 | 8 | 39251052 | 39380543 | 0 | 17 |
| TSNARE1 | 3 | 19 | 8 | 1.43E+08 | 1.43E+08 | 0.004961 | 5 |
| ADGRB1 | 2 | 31 | 8 | 1.44E+08 | 1.44E+08 | 0.000764 | 5 |
| PTPRD | 2 | 50 | 9 | 9364129 | 9724338 | 0 | 13 |
| SLC24A2 | 3 | 12 | 9 | 19684508 | 19918020 | 0.00296 | 8 |
| MLLT3 | 12 | 12 | 9 | 20329973 | 20653068 | 0 | 15 |
| MIR4473 | 1 | 1 | 9 | 20329973 | 20491397 | 0 | 15 |
| MIR4474 | 1 | 1 | 9 | 20491398 | 20548455 | 2.33E-06 | 13 |
| FOCAD | 48 | 48 | 9 | 20653069 | 21036488 | 0 | 20 |
| FOCAD-AS1 | 2 | 2 | 9 | 20653069 | 20726657 | 0 | 16 |
| MIR491 | 1 | 1 | 9 | 20653069 | 20726657 | 0 | 16 |
| SNORA30B | 1 | 1 | 9 | 20726658 | 20792331 | 0 | 17 |
| HACD4 | 10 | 10 | 9 | 20847854 | 21036488 | 0 | 20 |
| IFNB1 | 1 | 1 | 9 | 21068773 | 21111353 | 0 | 25 |
| IFNW1 | 1 | 1 | 9 | 21111354 | 21145148 | 0 | 26 |
| IFNA21 | 1 | 1 | 9 | 21151511 | 21227899 | 0 | 28 |
| IFNA4 | 1 | 1 | 9 | 21151511 | 21227899 | 0 | 28 |
| IFNA7 | 1 | 1 | 9 | 21151511 | 21227899 | 0 | 28 |
| IFNA10 | 1 | 1 | 9 | 21151511 | 21227899 | 0 | 28 |
| IFNA16 | 1 | 1 | 9 | 21151511 | 21227899 | 0 | 28 |
| IFNA17 | 1 | 1 | 9 | 21151511 | 21263438 | 0 | 29 |
| IFNA14 | 1 | 1 | 9 | 21227900 | 21263438 | 0 | 29 |
| IFNA22P | 1 | 1 | 9 | 21276902 | 21292379 | 0 | 31 |
| IFNA5 | 1 | 1 | 9 | 21292380 | 21306980 | 0 | 32 |
| KLHL9 | 1 | 1 | 9 | 21326775 | 21369720 | 0 | 35 |
| IFNA6 | 1 | 1 | 9 | 21335295 | 21369720 | 0 | 35 |
| IFNA13 | 1 | 1 | 9 | 21335295 | 21369720 | 0 | 35 |
| IFNA2 | 1 | 1 | 9 | 21369721 | 21400901 | 0 | 36 |
| IFNA8 | 1 | 1 | 9 | 21405710 | 21449454 | 0 | 38 |
| IFNA1 | 1 | 1 | 9 | 21405710 | 21449454 | 0 | 38 |
| MIR31HG | 5 | 5 | 9 | 21449455 | 21562622 | 0 | 45 |
| IFNE | 1 | 1 | 9 | 21479783 | 21493498 | 0 | 40 |
| MIR31 | 1 | 1 | 9 | 21508690 | 21518452 | 0 | 42 |
| MTAP | 8 | 8 | 9 | 21801709 | 21872025 | 0 | 84 |
| CDKN2A-DT | 1 | 1 | 9 | 21965764 | 21968051 | 0 | 108 |
| CDKN2A | 7 | 7 | 9 | 21965764 | 21998559 | 0 | 117 |
| CDKN2B-AS1 | 21 | 21 | 9 | 21991961 | 22124466 | 0 | 106 |
| CDKN2B | 3 | 3 | 9 | 22001639 | 22012853 | 0 | 103 |
| DMRTA1 | 2 | 2 | 9 | 22444685 | 22462834 | 0 | 54 |
| LINC01239 | 6 | 6 | 9 | 22637796 | 22826300 | 0 | 48 |
| LOC101929563 | 28 | 28 | 9 | 23492553 | 23745570 | 0 | 34 |
| ELAVL2 | 16 | 16 | 9 | 23648018 | 24012269 | 0 | 32 |
| IZUMO3 | 9 | 9 | 9 | 24520178 | 24732076 | 0 | 28 |
| TUSC1 | 1 | 1 | 9 | 25493325 | 25845834 | 0 | 22 |
| LINC01241 | 6 | 6 | 9 | 25493325 | 25845834 | 0 | 22 |
| LOC100506422 | 7 | 7 | 9 | 26057761 | 26398905 | 0 | 20 |
| CAAP1 | 7 | 7 | 9 | 26742841 | 26976980 | 0 | 16 |
| PLAA | 16 | 16 | 9 | 26856934 | 26976980 | 0 | 15 |
| IFT74 | 25 | 25 | 9 | 26856934 | 27433669 | 0 | 15 |
| IFT74-AS1 | 2 | 2 | 9 | 26856934 | 26976980 | 0 | 15 |
| LRRC19 | 5 | 5 | 9 | 26976981 | 27433669 | 4.6E-06 | 14 |
| TEK | 26 | 26 | 9 | 26976981 | 27433669 | 4.6E-06 | 14 |
| LINC00032 | 17 | 17 | 9 | 26976981 | 27433669 | 4.6E-06 | 14 |
| EQTN | 9 | 9 | 9 | 26976981 | 27433669 | 4.6E-06 | 14 |
| MOB3B | 4 | 4 | 9 | 26976981 | 27562984 | 0 | 14 |
| IFNK | 2 | 2 | 9 | 27433670 | 27562984 | 0 | 13 |
| C9orf72 | 14 | 14 | 9 | 27433670 | 27757634 | 0 | 13 |
| LINGO2 | 10 | 10 | 9 | 27757635 | 29335491 | 0 | 15 |
| MIR876 | 1 | 1 | 9 | 28770776 | 29041109 | 9.09E-05 | 11 |
| MIR873 | 1 | 1 | 9 | 28770776 | 29041109 | 9.09E-05 | 11 |
| KCNT1 |  |  | 9 | 1.39E+08 | 1.39E+08 | 0.002174 | 7 |
| PAPSS2 | 6 | 13 | 10 | 89475663 | 89513674 | 9.05E-06 | 7 |
| ATAD1 | 13 | 13 | 10 | 89499305 | 89610174 | 0 | 10 |
| CFL1P1 | 4 | 4 | 10 | 89548650 | 89610174 | 2.33E-06 | 10 |
| KLLN | 1 | 1 | 10 | 89618825 | 89625414 | 0 | 13 |
| PTEN | 12 | 12 | 10 | 89618825 | 89776422 | 0 | 22 |
| RNLS | 9 | 9 | 10 | 89849258 | 90357833 | 2.33E-06 | 11 |
| LIPJ | 11 | 11 | 10 | 90249140 | 90385967 | 0 | 10 |
| LIPF | 12 | 12 | 10 | 90413137 | 90562127 | 9.05E-06 | 10 |
| LIPK | 9 | 9 | 10 | 90413137 | 90562127 | 9.05E-06 | 10 |
| LIPN | 9 | 9 | 10 | 90413137 | 90562127 | 9.05E-06 | 10 |
| LIPM | 9 | 9 | 10 | 90562128 | 90596451 | 0 | 9 |
| ANKRD22 | 6 | 6 | 10 | 90562128 | 90809529 | 0 | 9 |
| STAMBPL1 | 11 | 11 | 10 | 90596452 | 90809529 | 0.001486 | 8 |
| ACTA2-AS1 | 5 | 5 | 10 | 90596452 | 90809529 | 0.001486 | 8 |
| ACTA2 | 11 | 11 | 10 | 90596452 | 90809529 | 0.001486 | 8 |
| FAS | 10 | 10 | 10 | 90596452 | 90809529 | 0.001486 | 8 |
| FAS-AS1 | 1 | 1 | 10 | 90596452 | 90809529 | 0.001486 | 8 |
| MIR4679-2 | 1 | 1 | 10 | 90816735 | 90853476 | 0.000194 | 6 |
| MIR4679-1 | 1 | 1 | 10 | 90816735 | 90853476 | 0.000194 | 6 |
| NKX6-2 | 3 | 3 | 10 | 1.35E+08 | 1.35E+08 | 9.73E-05 | 5 |
| CFAP46 | 23 | 58 | 10 | 1.35E+08 | 1.35E+08 | 8.24E-05 | 6 |
| ADGRA1-AS1 | 3 | 3 | 10 | 1.35E+08 | 1.35E+08 | 0.00139 | 5 |
| ADGRA1 | 6 | 8 | 10 | 1.35E+08 | 1.35E+08 | 2.33E-06 | 5 |
| KNDC1 | 33 | 33 | 10 | 1.35E+08 | 1.35E+08 | 6.84E-06 | 5 |
| UTF1 | 2 | 2 | 10 | 1.35E+08 | 1.35E+08 | 0.000106 | 5 |
| VENTX | 3 | 3 | 10 | 1.35E+08 | 1.35E+08 | 0.000106 | 6 |
| MIR202HG | 3 | 3 | 10 | 1.35E+08 | 1.35E+08 | 0.000127 | 5 |
| MIR202 | 1 | 1 | 10 | 1.35E+08 | 1.35E+08 | 0.000127 | 5 |
| OR52N5 | 1 | 1 | 11 | 5787479 | 5804546 | 0 | 18 |
| OR52N1 | 1 | 1 | 11 | 5809050 | 5810167 | 0 | 12 |
| OR4P4 | 1 | 1 | 11 | 55380607 | 55427571 | 2.22E-05 | 5 |
| OR4S2 | 1 | 1 | 11 | 55380607 | 55427571 | 2.22E-05 | 5 |
| PRH1-PRR4 | |  | 12 | 11218334 | 11251512 | 0 | 7 |
| PRH1 |  |  | 12 | 11218334 | 11251512 | 0 | 7 |
| PRH1-TAS2R14 | |  | 12 | 11218334 | 11251512 | 0 | 7 |
| TAS2R43 | 1 | 1 | 12 | 11228845 | 11246378 | 0 | 7 |
| ANHX | 12 | 12 | 12 | 1.34E+08 | 1.34E+08 | 0 | 5 |
| FRY | 48 | 61 | 13 | 32723557 | 32892538 | 5.24E-05 | 5 |
| ZAR1L | 6 | 6 | 13 | 32861810 | 32892538 | 5.24E-05 | 5 |
| BRCA2 | 27 | 27 | 13 | 32861810 | 33056012 | 0 | 6 |
| PDS5B | 35 | 35 | 13 | 33159360 | 33389134 | 0.000616 | 5 |
| LINC00423 | 1 | 5 | 13 | 33159360 | 33431330 | 0.000616 | 5 |
| ITM2B | 6 | 6 | 13 | 48796783 | 48853910 | 2.33E-06 | 6 |
| RB1-DT | 3 | 3 | 13 | 48869322 | 48884943 | 0 | 10 |
| RB1 | 27 | 27 | 13 | 48875658 | 49057901 | 0 | 16 |
| LPAR6 | 9 | 9 | 13 | 48982338 | 49019160 | 0 | 11 |
| RCBTB2 | 20 | 20 | 13 | 49057902 | 49123022 | 0 | 9 |
| LINC00462 | 3 | 3 | 13 | 49123023 | 49189678 | 0 | 8 |
| CYSLTR2 | 7 | 7 | 13 | 49189679 | 49313993 | 0 | 7 |
| FNDC3A | 30 | 30 | 13 | 49544376 | 49789749 | 5.24E-05 | 5 |
| UPF3A | 18 | 18 | 13 | 1.15E+08 | 1.15E+08 | 0 | 7 |
| CHAMP1 | 5 | 5 | 13 | 1.15E+08 | 1.15E+08 | 0 | 9 |
| LINC01054 | 3 | 3 | 13 | 1.15E+08 | 1.15E+08 | 0 | 8 |
| LOC105370401 | 10 | 10 | 14 | 22798784 | 22964322 | 0 | 11 |
| LINC02315 | 1 | 5 | 14 | 41604259 | 41610284 | 0 | 9 |
| ASPG | 15 | 16 | 14 | 1.05E+08 | 1.05E+08 | 0 | 5 |
| MIR203A | 1 | 1 | 14 | 1.05E+08 | 1.05E+08 | 0 | 5 |
| MIR203B | 1 | 1 | 14 | 1.05E+08 | 1.05E+08 | 0 | 5 |
| LOC105370708 | 3 | 3 | 14 | 1.05E+08 | 1.05E+08 | 0 | 6 |
| KIF26A | 15 | 15 | 14 | 1.05E+08 | 1.05E+08 | 0 | 6 |
| MIR4507 | 1 | 1 | 14 | 1.06E+08 | 1.06E+08 | 0 | 5 |
| MIR4538 | 1 | 4 | 14 | 1.06E+08 | 1.06E+08 | 0 | 5 |
| MIR4537 | 2 | 2 | 14 | 1.06E+08 | 1.06E+08 | 0 | 5 |
| MIR4539 | 1 | 2 | 14 | 1.06E+08 | 1.06E+08 | 0 | 5 |
| FAM30A | 6 | 6 | 14 | 1.06E+08 | 1.06E+08 | 0 | 6 |
| LOC101927079 | 2 | 10 | 15 | 22335598 | 22383757 | 0 | 7 |
| B2M | 4 | 4 | 15 | 44989060 | 45031128 | 0 | 5 |
| RBFOX1 | 3 | 24 | 16 | 6288694 | 7004043 | 0 | 15 |
| WWOX | 1 | 12 | 16 | 78367828 | 78908173 | 0 | 21 |
| VPS53 | 1 | 24 | 17 | 478699 | 489523 | 0 | 5 |
| ATP1B2 | 9 | 9 | 17 | 7541724 | 7563500 | 0 | 6 |
| TP53 | 18 | 18 | 17 | 7570798 | 7593227 | 0 | 9 |
| DNAH9 | 67 | 70 | 17 | 11514129 | 11896638 | 0 | 9 |
| ZNF18 | 12 | 12 | 17 | 11844990 | 11930484 | 0 | 8 |
| MAP2K4 | 12 | 12 | 17 | 11896639 | 12077866 | 0 | 12 |
| MIR744 | 1 | 1 | 17 | 11976614 | 12003127 | 0 | 6 |
| LINC00670 | 4 | 4 | 17 | 12401948 | 12581725 | 3.95E-05 | 5 |
| NF1 | 61 | 61 | 17 | 29388421 | 29705018 | 0 | 11 |
| OMG | 2 | 2 | 17 | 29613075 | 29682998 | 2.43E-05 | 5 |
| EVI2B | 2 | 2 | 17 | 29613075 | 29682998 | 2.43E-05 | 5 |
| EVI2A | 3 | 3 | 17 | 29613075 | 29682998 | 2.43E-05 | 5 |
| KRTAP9-6 | 1 | 1 | 17 | 39421405 | 39430869 | 0 | 10 |
| KRTAP9-7 | 1 | 1 | 17 | 39431857 | 39432570 | 0 | 5 |
| MAPK4 | 6 | 6 | 18 | 48018714 | 48327681 | 4.38E-05 | 6 |
| MRO | 9 | 9 | 18 | 48188737 | 48368127 | 0 | 7 |
| ME2 | 16 | 16 | 18 | 48392286 | 48507920 | 0 | 11 |
| ELAC1 | 4 | 4 | 18 | 48457280 | 48518035 | 0 | 12 |
| SMAD4 | 12 | 12 | 18 | 48553523 | 48611791 | 0 | 19 |
| MEX3C | 2 | 2 | 18 | 48699757 | 48768742 | 0 | 11 |
| LINC01630 | 4 | 4 | 18 | 48805384 | 49095861 | 2.33E-06 | 8 |
| DCC | 22 | 29 | 18 | 50628715 | 51216293 | 9.05E-06 | 7 |
| MIR4528 | 1 | 1 | 18 | 50670186 | 50769780 | 0.000229 | 5 |
| ADNP2 | 4 | 4 | 18 | 77844967 | 77936088 | 1.56E-05 | 5 |
| PARD6G-AS1 | 5 | 5 | 18 | 77893655 | 77951273 | 0 | 6 |
| PARD6G | 3 | 3 | 18 | 77893655 | 78005472 | 0 | 16 |
| STK11 | 5 | 10 | 19 | 1220886 | 1229793 | 0 | 6 |
| CBARP | 9 | 9 | 19 | 1227561 | 1238671 | 0 | 7 |
| MIR526A1 | 1 | 1 | 19 | 54209469 | 54301580 | 0.000196 | 5 |
| MIR520C | 1 | 1 | 19 | 54209469 | 54301580 | 0.000196 | 5 |
| MIR518C | 1 | 1 | 19 | 54209469 | 54301580 | 0.000196 | 5 |
| MIR524 | 1 | 1 | 19 | 54209469 | 54301580 | 0.000196 | 5 |
| MIR517A | 1 | 1 | 19 | 54209469 | 54301580 | 0.000196 | 5 |
| MIR519D | 1 | 1 | 19 | 54209469 | 54301580 | 0.000196 | 5 |
| MIR521-2 | 1 | 1 | 19 | 54209469 | 54301580 | 0.000196 | 5 |
| MIR520D | 1 | 1 | 19 | 54209469 | 54301580 | 0.000196 | 5 |
| MIR517B | 1 | 1 | 19 | 54209469 | 54301580 | 0.000196 | 5 |
| MIR520G | 1 | 1 | 19 | 54209469 | 54301580 | 0.000196 | 5 |
| MIR516B2 | 1 | 1 | 19 | 54209469 | 54301580 | 0.000196 | 5 |
| MIR526A2 | 1 | 1 | 19 | 54209469 | 54301580 | 0.000196 | 5 |
| MIR518E | 1 | 1 | 19 | 54209469 | 54301580 | 0.000196 | 5 |
| MIR518A1 | 1 | 1 | 19 | 54209469 | 54301580 | 0.000196 | 5 |
| MIR518D | 1 | 1 | 19 | 54209469 | 54301580 | 0.000196 | 5 |
| MIR516B1 | 1 | 1 | 19 | 54209469 | 54301580 | 0.000196 | 5 |
| MIR518A2 | 1 | 1 | 19 | 54209469 | 54301580 | 0.000196 | 5 |
| MIR517C | 1 | 1 | 19 | 54209469 | 54301580 | 0.000196 | 5 |
| MIR520H | 1 | 1 | 19 | 54209469 | 54301580 | 0.000196 | 5 |
| MIR521-1 | 1 | 1 | 19 | 54209469 | 54301580 | 0.000196 | 5 |
| MIR522 | 1 | 1 | 19 | 54209469 | 54301580 | 0.000196 | 5 |
| MIR519A1 | 1 | 1 | 19 | 54209469 | 54301580 | 0.000196 | 5 |
| MIR527 | 1 | 1 | 19 | 54209469 | 54301580 | 0.000196 | 5 |
| MIR516A1 | 1 | 1 | 19 | 54209469 | 54301580 | 0.000196 | 5 |
| MIR1283-2 | 1 | 1 | 19 | 54209469 | 54301580 | 0.000196 | 5 |
| MIR516A2 | 1 | 1 | 19 | 54209469 | 54301580 | 0.000196 | 5 |
| MIR519A2 | 1 | 1 | 19 | 54209469 | 54301580 | 0.000196 | 5 |
| MIR371A | 1 | 1 | 19 | 54209469 | 54301580 | 0.000196 | 5 |
| MIR371B | 1 | 1 | 19 | 54209469 | 54301580 | 0.000196 | 5 |
| MIR372 | 1 | 1 | 19 | 54209469 | 54301580 | 0.000196 | 5 |
| MIR373 | 1 | 1 | 19 | 54209469 | 54301580 | 0.000196 | 5 |
| NLRP12 | 13 | 13 | 19 | 54209469 | 54359183 | 0.000196 | 5 |
| CENPBD1P1 | 2 | 2 | 19 | 59085429 | 59097681 | 0 | 5 |
| MACROD2 | 3 | 21 | 20 | 14567270 | 15226720 | 0 | 30 |
| MACROD2-AS1 | 4 | 4 | 20 | 14859588 | 14917621 | 0 | 14 |
| PRMT2 | 7 | 15 | 21 | 48070415 | 48090744 | 4.59E-05 | 5 |
| ZDHHC8 | 2 | 12 | 22 | 20134997 | 20136452 | 0.002014 | 5 |
| CCDC188 | 2 | 9 | 22 | 20134998 | 20136452 | 0.009908 | 5 |
| MIR5571 | 1 | 1 | 22 | 23199140 | 23237086 | 0.004215 | 6 |
| IGLL5 | 2 | 3 | 22 | 23199140 | 23237163 | 0.004215 | 6 |
| RSPH14 |  |  | 22 | 23409509 | 23409931 | 0.002092 | 5 |
| GSTT4 |  |  | 22 | 24347762 | 24391059 | 0.004523 | 6 |
| LOC391322 | 3 | 3 | 22 | 24348024 | 24391059 | 0.004523 | 6 |
| GSTT1-AS1 | 1 | 1 | 22 | 24348024 | 24391059 | 0.004523 | 6 |
| GSTT1 | 7 | 7 | 22 | 24348024 | 24391059 | 0.004523 | 6 |
| GSTTP2 | 1 | 5 | 22 | 24348024 | 24391059 | 0.004523 | 6 |
| CABIN1 |  |  | 22 | 24553887 | 24557098 | 0.002005 | 5 |
| LINC02558 |  |  | 22 | 32413522 | 32428266 | 0.002658 | 6 |
| PARVB | 4 | 17 | 22 | 44529841 | 44552636 | 0.002078 | 6 |
| TAFA5 | 3 | 5 | 22 | 48992105 | 49155093 | 0.007799 | 5 |
| C22orf34 | 5 | 11 | 22 | 49989769 | 50043752 | 0.000173 | 6 |
| MAPK12 | 10 | 13 | 22 | 50694019 | 50709128 | 0.00094 | 6 |
| MAPK11 | 14 | 14 | 22 | 50699567 | 50709128 | 0.00094 | 6 |
| PLXNB2 | 50 | 50 | 22 | 50709129 | 50758450 | 0.000741 | 6 |
| DENND6B | 19 | 20 | 22 | 50745332 | 50758450 | 0.005163 | 5 |
| ODF3B | 6 | 7 | 22 | 50969131 | 50976399 | 0.003308 | 5 |
| PUDP | 7 | 7 | X | 6936405 | 7150085 | 2.33E-06 | 6 |
| STS | 19 | 19 | X | 7045619 | 7290374 | 3.3E-05 | 6 |
| MIR4767 | 1 | 1 | X | 7045619 | 7150085 | 5.02E-05 | 5 |
| VCX | 3 | 3 | X | 7730601 | 7982238 | 0.000285 | 5 |
| PNPLA4 | 9 | 9 | X | 7730601 | 7982238 | 0.000285 | 5 |
| VCX2 | 3 | 3 | X | 8125047 | 8395823 | 0.000311 | 5 |
| DMD | 30 | 91 | X | 31673069 | 33374633 | 9.05E-06 | 14 |
| MIR3915 | 1 | 1 | X | 32575651 | 32638408 | 9.05E-06 | 5 |

**Supplementary Table 6.** Recurrently homozygously deleted genes in the POG cohort of metastatic cancers. Blank exon values indicate that no exons of that gene were affected by recurrent homozygous deletion. False-discovery rate corrected P-values are shown.
